# A chromosome level reference genome of Diviner’s sage (*Salvia divinorum*) provides insight into salvinorin A biosynthesis

**DOI:** 10.1101/2024.09.08.611878

**Authors:** Scott A. Ford, Rob W. Ness, Moonhyuk Kwon, Dae-Kyun Ro, Michael A. Phillips

## Abstract

**Background:** Diviner’s sage (*Salvia divinorum*; Lamiaceae) is the source of the powerful hallucinogen salvinorin A (SalA). This neoclerodane diterpenoid is an agonist of the human Κ opioid receptor with potential medical applications in the treatment of chronic pain, addiction, and post-traumatic stress disorder. Only two steps of the approximately twelve step biosynthetic sequence leading to SalA have been resolved to date.

**Results:** To facilitate pathway elucidation in this ethnomedicinal plant species, here we report a chromosome level genome assembly. A high-quality genome sequence was assembled with an N50 value of 41.4 Mb and a BUSCO completeness score of 98.4%. The diploid (2n = 22) genome of ∼541 Mb is comparable in size and ploidy to most other members of this genus. Two diterpene biosynthetic gene clusters were identified and are highly enriched in previously unidentified cytochrome P450s as well as crotonolide G synthase, which forms the dihydrofuran ring early in the SalA pathway. Coding sequences for other enzyme classes with likely involvement in downstream steps of the SalA pathway (BAHD acyl transferases, alcohol dehydrogenases, and O-methyl transferases) were scattered throughout the genome with no clear indication of clustering. Differential gene expression analysis suggests that most of these genes are not inducible by methyl jasmonate treatment.

**Conclusions:** This genome sequence and associated gene annotation are among the highest resolution in *Salvia*, a genus well known for the medicinal properties of its members. Here we have identified the cohort of genes responsible for the remaining steps in the SalA pathway. This genome sequence and associated candidate genes will facilitate the elucidation of SalA biosynthesis and enable an exploration of its full clinical potential.

## Introduction

*Salvia divinorum* (Epling and Játiva-M.), also known as Ska María Pastora or Diviner’s sage, is a hallucinogenic plant used in ritualistic ceremonies by the Mazatec people of Oaxaca, Mexico [1]. Its hallucinogenic properties are due to salvinorin A (SalA), a neoclerodane diterpenoid produced in glandular trichomes on the surfaces of leaves. SalA is a highly selective and potent K-opioid receptor (KOR) agonist and the first known nitrogen-free ligand for this receptor family [2]. Its powerful dissociative properties and popularity as a recreational drug have led to its criminalization in Canada, much of western Europe, Japan, South Korea, and 33 states of the United States. However, it remains legal in many countries, including Mexico, home to its only natural endemic population.

SalA belongs to the neoclerodane subfamily of labdane diterpenoid secondary metabolites. Diterpenoids represent enormous chemical diversity in the plant kingdom numbering close to 13,000, including the salvinorins unique to *S. divinorum*. [3–5]. Although widespread in the Viridiplantae, diterpenoids are especially abundant in the Lamiaceae, which are the source of approximately one quarter (∼3,000) of this total.

There are at least 22 diterpenes in the neoclerodane diterpenoid group that have now been isolated from *S. divinorum* [6], but SalA is the most abundant and the only natural product in this species thus far to demonstrate pharmacological activity [7]. It is the first reported diterpene to possess psychoactive properties [8]. It displays high affinity for the KOR (Ki of ∼ 4 nM), one of five related opioid receptors in the human body that modulate pain, emotional control, memory, learning, and mood (Schwarzer 2009). It has thus been at the center of research efforts to understand the medical potential of this plant species. The selectivity and potency of SalA towards KOR suggest therapeutic potential in the treatment of pain, inflammation, and depression [9–11]. Pain relief has traditionally relied on agonism of the related μ opioid receptor (MOR), which leads to dependency and addiction. In contrast, agonism of KOR is not associated with addiction and therefore represents a critical alternative in the development of pharmaceuticals to treat chronic pain. Indeed, SalA agonism of KOR may exert anti-addictive effects [7, 12].

Other clerodane diterpenoids of *S. divinorum* such as salvinorin B and divinatorin D and E are also KOR agonists with Ki values ranging from 65 to 418 nM [13]. The structural diversity of bioactive diterpenoids in this species has led to new insights into opioid receptor function [2] as well as synthetic analogs whose potency, selectivity, and bioavailability exceed SalA [14–16]. However, most synthetic protocols depend on semi-synthesis from advanced precursors [17, 18] and are economically unfeasible from an industrial standpoint. Thus, a detailed understanding of the biosynthetic pathway responsible for the bioactive diterpenoids from *S. divinorum* is essential to explore their full clinical potential.

The biosynthesis of typical labdane diterpenes, including the neoclerodane SalA, commences with an initial cyclization of GGDP by a class II diterpene synthase to a bicyclic product which retains the diphosphate group [4]. This is followed by a type I diterpene synthase, which removes the diphosphate group and may also introduce additional rings, double bonds, or hydroxyl groups [19]. In the case of SalA, GGDP is cyclized to kolavenyl diphosphate (KDP) by the type II diterpene synthase kolavenyl (or clerodienyl) diphosphate synthase (KDS) (Figure 1; step 1) [20, 21]. KDP is subsequently dephosphorylated to kolavenol by an as-of-yet unidentified enzyme or enzymes (step 2). The dihydrofuran ring is next introduced by crotonolide G synthase (step 3) [22]. This cytochrome P450 introduces an O atom at C16 which initiates a nucleophilic attack to form the dihydrofuran ring, displacing the C15 alcohol as water. It is presently unclear whether the furan ring, which confers the KOR binding activity of SalA, is formed in the next biosynthetic step or later in the pathway. In case of the former, desaturation of the dihydrofuran ring of crotonolide G would produce *trans*-annonene (Figure 1, step 4), but this intermediate has not been detected in *S. divinorum* to date. Alternatively, the furan ring may be completed after oxidation at C18 (step 4a). Evidence for the latter view consists of a partially characterized C1 (CYP728D25) and C18 hydroxylase (CYP728D26) from *S. divinorum* which act on crotonolide G as substrate [23]. This scenario may implicate salvidivins or salvinicins as intermediates in the SalA pathway, which naturally occur in *S. divinorum* [6]. However, they are not necessarily required for SalA biosynthesis and may constitute side products.

**Figure 1.**
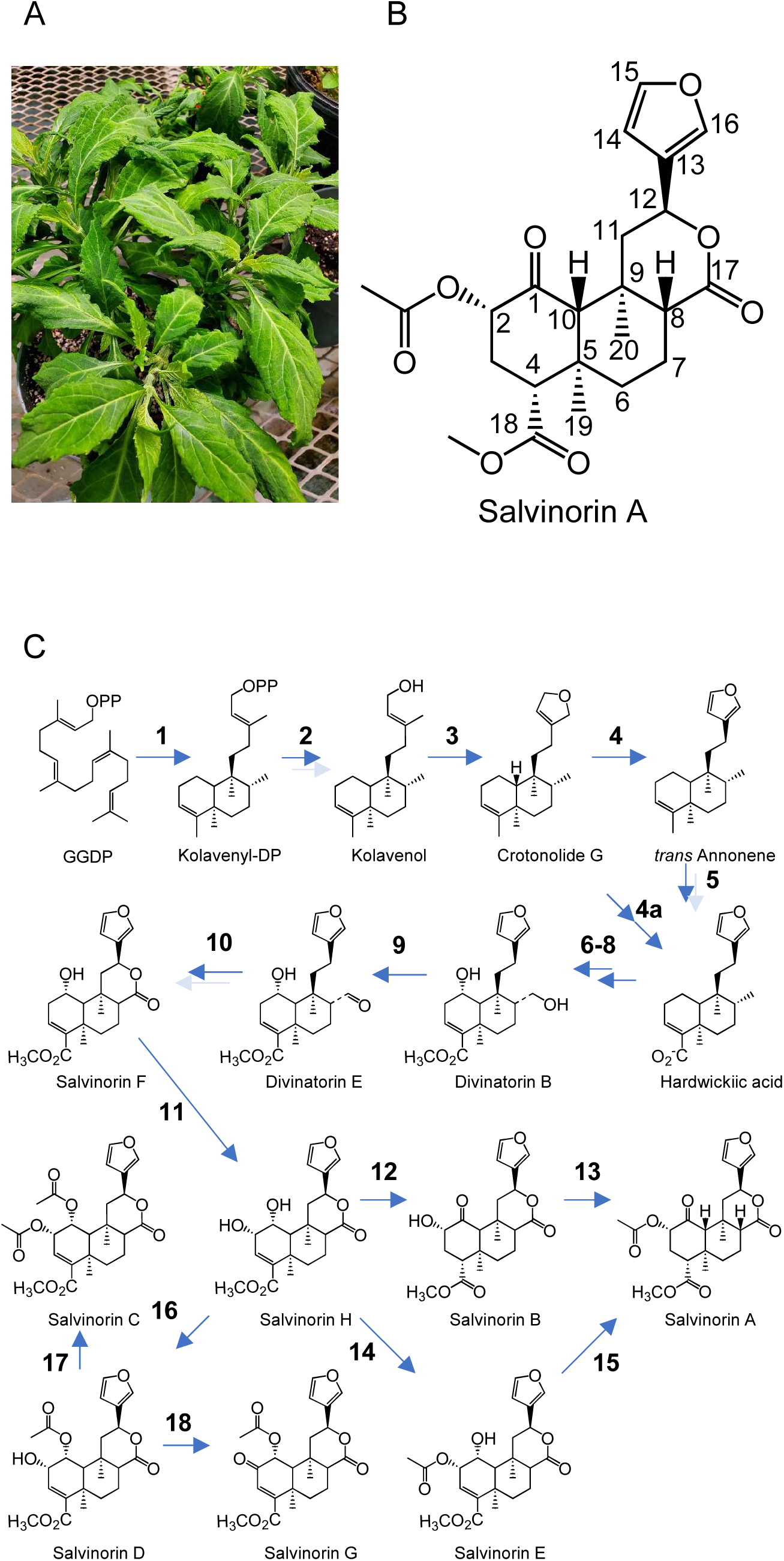
Diviner’s sage (*Salvia divinorum*) plants and salvinorin A proposed biosynthetic route. A, *S. divinorum;* B, Structure of salvinorin A showing numbering used in text. C, Proposed biosynthetic pathway based on biochemical characterization and intermediates isolated from *S. divinorum* leaves. The enzyme catalyzing reaction **1** has been described by Chen et al. 2017 and Pelot et al. 2017 while **3** was described in Kwon et al. 2022. Two dark arrows signify multi-enzyme sequences whose order is uncertain. Mixed light and dark arrows indicate sequences which may be catalyzed by single or multiple enzymes.

As originally proposed by Chen et al. (2017), the remainder of the SalA pathway may be postulated by structures of diterpenoids isolated from *S. divinorum* leaves [6, 13, 24, 25] (Fig. 1). Following (or perhaps preceding) assembly of the furan ring (steps 1-4), oxidation at C18 leads to hardwickiic acid (step 5). Hydroxylation at C1 and C17 and methylation C18 yield divinatorin B (steps 6-8), although the exact order is uncertain. Subsequent dehydrogenation at C17 produces the aldehyde divinatorin E, and lactonization between C12 and C17, leading to the salvinorins, presumably stems from oxidations at C12 and C17 followed by elimination of water (steps 9-10). A hydroxylation at C2 of salvinorin F arrives at the vicinal alcohol salvinorin H, which may proceed to SalA by dehydrogenation at C1 (salvinorin B) and acetylation at C2 or by the same steps in reverse, passing instead through salvinorin E (steps 12-15). Acetylation at C1 instead of dehydrogenation to a ketone leads to salvinorins C, D, and G (steps 16-18). A reference genome of *S. divinorum* would not only facilitate identification of candidate genes for the steps outlined above but also provide valuable insight into the diterpenoid biosynthetic gene complement of one of nature’s most medically important plant genera.

*Salvia* (Lamiaceae:Nepetoideae: Mentheae: Salviinae) is the largest genus in the mint family with its over 1,000 species making up ∼15% of the family. The intrageneric relationships within the *Salvia* genus have been refined using concatenated, coalescent, and network-based nuclear phylogenies [26], which support a common backbone topology of ∼10 monophyletic subgenera. Assembled genome sequences for six economically important *Salvia* species have been reported. These include *danshen* (*S. miltiorrhiza*) [27, 28] and *nan danshen* (*S. bowleyana*) [29], sources of tanshinones, rosemary (*S. Rosmarinus*) [30], source of rosmarinic acid, chia (*S. hispanica*) [31], a rich source of several dietary supplements, and common sage (*S. officinalis*) [32], which is widely used in cooking. These genomes tend to be diploid and range in size from 400 to 600 Mbp except for the ornamental tetraploid *S. splendens* (807 Mbp) [33, 34].

The closest sequenced species which produces diterpenes similar to those found in *S. divinorum* is the traditional Chinese medicine herb skullcap (*Scutellaria barbata*) [35]. Skullcap accumulates clerodane diterpenes with anti-cancer applications [36], and its genome may offer clues to SalA biosynthesis. In the Lamiaceae, genes for diterpene biosynthesis tend to organize into discrete gene clusters consisting of microsyntenic blocks [37, 38] also referred to as biosynthetic gene clusters (BGC) [32]. Here we report a chromosome level reference genome of *S. divinorum*, an examination of the structure of its BGCs, and a detailed inventory of genes identified as candidates for involvement in the SalA pathway.

## Results

### Assembly of the *S. divinorum* genome

A preparation of genomic DNA purified from *S. divinorum* leaves showed an average length of >50 kb according to DNA tape station analysis. A total of 62 Gb of consensus HiFi reads were generated on the PacBio Revio platform (N50 read length 14,336 bp; (Supplementary Figure 1). *K*-mer distribution analysis of the reads predicted a genome size of ∼546 Mb, low levels of heterozygosity (0.192%), ∼53% repeat content, and a homozygous coverage of ∼30x (Supplementary Figure 2). Raw HiFi reads were assembled using hifiasm [39] and haplotypic duplication was removed from the haploid draft assembly with purge_dups [40]. Since many highly fragmented plastidic and mitochondrial DNA contigs were produced in the draft assembly, alternative approaches were taken to assemble the organellar genomes. Using ptGAUL [41], we assembled a complete, 150,899 bp circular chloroplast genome (Supplementary Figure 3). Two circular mitochondrial genomes were assembled (272,775 bp and 44,943 bp) by providing hifiasm with subsampled mitochondrial DNA (Supplementary Figure 4). A mitochondrial genome conformation of two circular chromosomes is a known feature of the *Salvia* genus [42, 43].

After filtering mitochondrial and plastidic contigs from the assembly, the resulting nuclear genome was 541 Mb (Figure 2), consistent with *K*-mer predictions and genome sizes of closely related species. General assembly statistics are summarized in Table 1. The final assembly featured an N50 score of 42.4 Mb (L50 = 5) with a maximum contig length over 72 Mb and a median length of ∼8.8 Mb. The BUSCO completeness of the genome was 98.4% when scored against the Embryophyta database [44]. Together, these statistics confirm that the assembly is of high quality with contiguity and completeness comparable to or exceeding the most recently sequenced *Salvia* genomes [29, 30, 32, 45].

**Figure 2.**
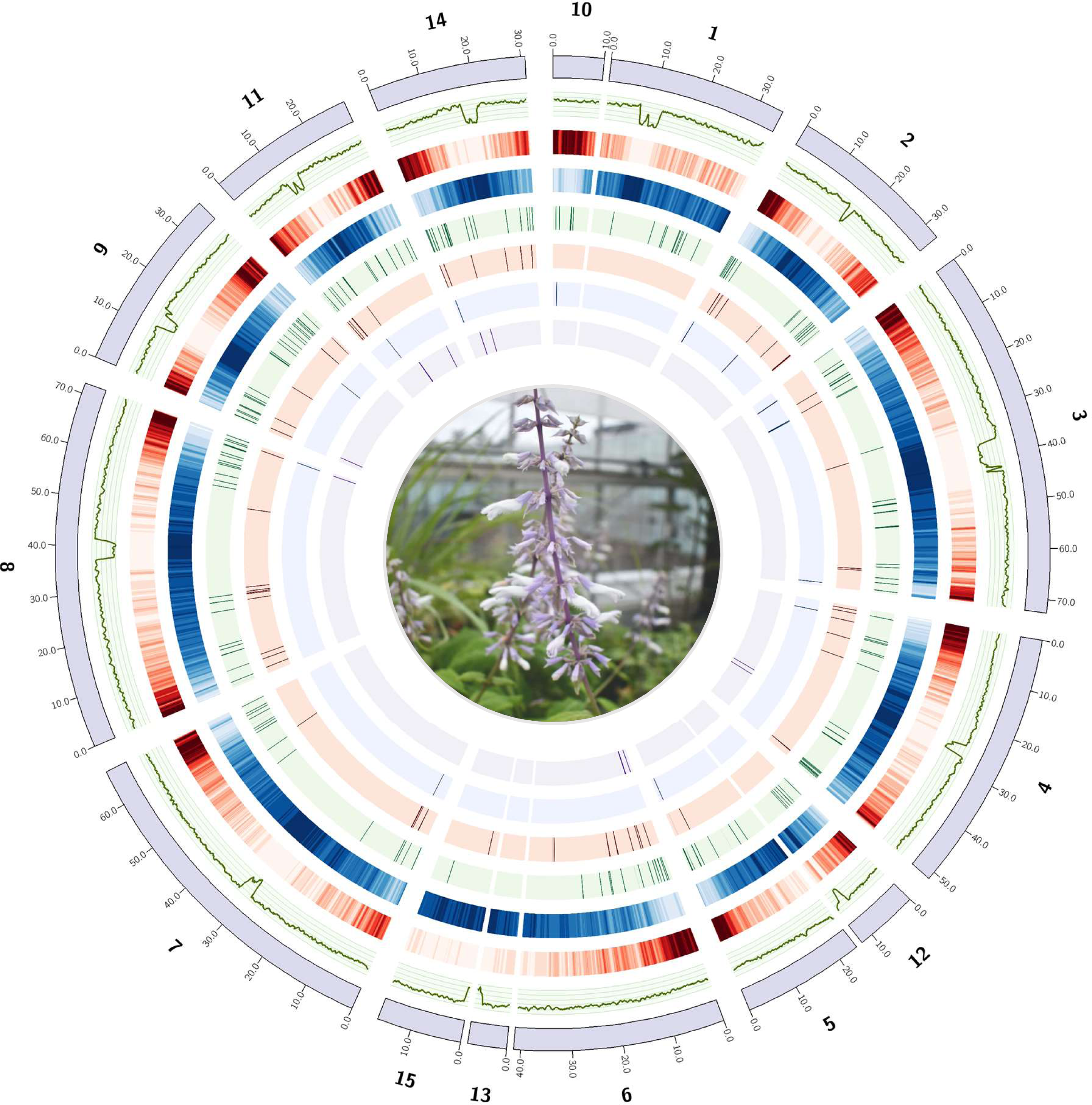
Circos plot of *Salvia divinorum* genome assembly. From outside to inside: scaffolds greater than 1Mb (purple ideograms), GC% (green line, range displayed; 20-45%), gene density (500Kb windows with 100Kb step size), repeat content (500Kb windows with 100Kb step size), position of putative diterpene biosynthetic genes, including cytochrome P450s (light green), BAHD acyltransferase (light red), O-methyl transferases (light blue), diterpene synthases (light purple).

**Table 1.** Genome assembly statistics for Diviner’s sage

| Assembly statistics |  | Repeat content |  | Annotation statistics |  |
| --- | --- | --- | --- | --- | --- |
| Total length | 541,447,152 bp | Repeat content (%) | 67.63 | Gene number | 34,514 |
| Total contigs | 28 | Retroelements (%) | 37.28 | Mean gene length (bp) | 2542.05 |
| N50 | 41,401,808 bp | SINES (%) | 0 | Mean exon length (bp) | 196.56 |
| L50 | 5 | LINES (%) | 0.34 | BUSCO<sup>1</sup> (protein) | 98.2% |
| Max contig length | 72,546,039 bp | LTR (%) | 36.94 | [S:85.7, D:12.5] |  |
| Median contig length | 8,837,828 bp | Transposons (%) | 3.17 |  |  |
| Contigs > 1Mb (%) | 50 | Unclassified (%) | 26.22 |  |  |
| Contigs > 100Kb (%) | 83.3 |  |  |  |  |
| BUSCO<sup>1</sup> (genome) | 98.4% |  |  |  |  |
| [S:85.5, D:12..9] |  |  |  |  |  |

<sup>1</sup> Using embryophyta\_odb10 database, n = 1614 genes

nThe repeat content in *S. divinorum* was 67.63%, with long terminal repeat (LTR) retrotransposons (RTs) accounting for 36.94% of the genome. Transposable elements constitute 3.17% while long interspersed nuclear elements (LINES) compose only 0.34%. As expected, regions of high repeat content were generally inversely associated with gene density, and the most repetitive regions of each chromosome were characterized by steep declines in GC content from ∼39% to ∼20% on average, likely indicative of centromeres (Figure 2). It is also noteworthy that repeat content was much higher throughout the inner parts of each chromosome and gene density higher in the telomere proximal regions.

### Genome annotation

Based on *ab initio* prediction as well as evidence from mRNA sequencing and protein homology, the *S. divinorum* nuclear genome was predicted to have 34,514 genes, with an average gene length of 2,542 bp and exon length of 197 bp (Table 1). Among the predicted genes, 92.7% (31,995) were functionally annotated with protein products, Gene Ontology terms, and/or identifiers from external protein databases queried using InterProScan [46], eggNOG-Mapper [47] and funannotate [48]. The gene content of the *S. divonorum* genome was comparable to similarly sized diploid genomes of *S. miltiorrhiza* (530.97 Mb; 32,191 genes) [45] and *S. officinalis* (480 Mb; 31,713 genes) [32], but considerably lower than *S. bowleyana* (462.44 Mb; 44,044 genes) [29].

Within the 150,899 bp chloroplast genome of *S. divinorum*, we annotated 116 unique genes, 4 encoding rRNA, 30 encoding tRNA and 82 encoding proteins (Supplementary Figure 3). The inverted repeat, small single copy and large single copy regions were 25,527 bp, 17,553 bp and 82,289 bp respectively. The structure of the *S. divinorum* chloroplast genome is consistent with the high conservation of plastid genome features within the genus [49]. In contrast, the structure of the mitochondrial genome is known to be diverse amongst *Salvia*, even at the intraspecies level [49]. In the two mitochondrial chromosomes of *S. divinorum*, we annotated 42 unique protein coding genes, 22 tRNA genes, and 3 rRNA genes (Supplementary Figure 4). In spite of >100-fold variation in mtDNA size across the plant kingdom, mitochondrial gene content varies little with about 60-70 genes found in mitogenomes of most plant species [50].

### Chromosomal synteny with *S. splendens*

We analyzed orthologous gene order using GENESPACE [51] and observed a high degree of synteny between the genomes of the diploid *S. divinorum* [52] and the tetraploid *S. splendens* (Figure 3), suggesting few large-scale chromosomal rearrangements in spite of the whole genome duplication in *S. splendens*. The shared synteny between these genomes also permitted the identification of smaller contigs in our *S. divinorum* assembly that can likely be grouped with larger contigs in chromosomes, but have been broken either at repeats near the centromere (contigs 5/12, 15/13) or at other chromosomal regions (contigs 10/1; 6/13) (Figure 2). Grouping these contigs together as chromosomes, our *S. divinorum* assembly has 11 nuclear chromosomes, as Reisfield reports (1993). Analyses of GC content, repeat density, and gene density indicate the presence of 11 centromeres (Figure 2). Though these results strongly suggest linkages between the aforementioned contigs, we have elected to keep them separate in our assembly since we have no direct sequence evidence to scaffold contigs together.

**Figure 3.**
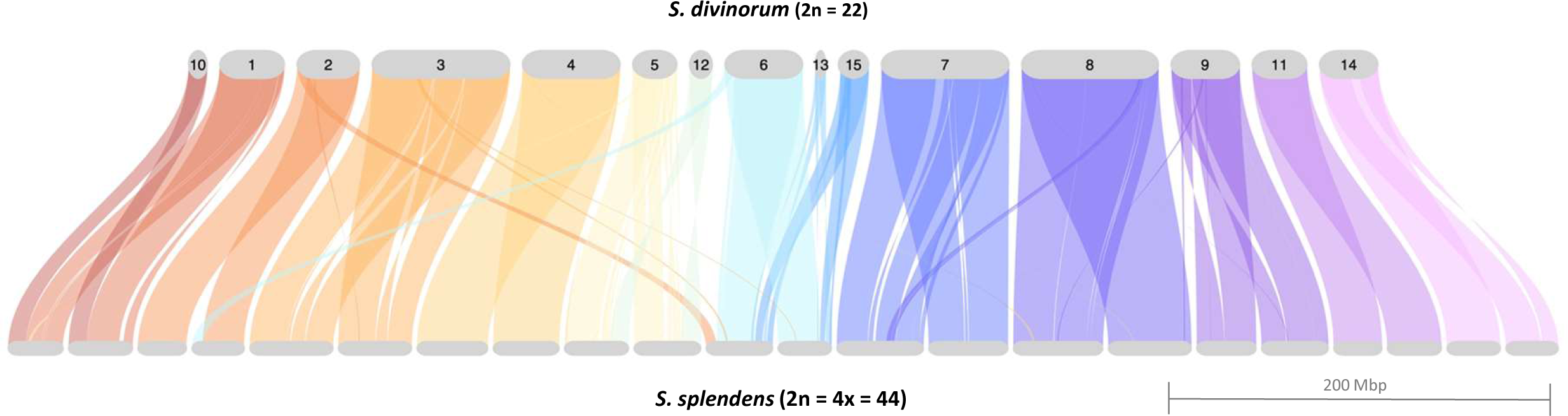
Riparian plot displaying syntenic relationship of Salvia divinorum and the tetraploid Salvia splendens [33]. Syntenic regions were identified using GENESPACE [84] which combines evidence from gene collinearity and sequence similarity.

### Genome evolution and phylogeny

To infer the evolutionary history of *S. divinorum*, we constructed a maximum likelihood phylogeny (Figure 4) of *S. divinorum* and six related species (*S. splendens*, *S. hispanica*, *S. rosmarinus*, *S. bowleyana*, *S. miltiorrhiza*, and *Sesamum indicum*) from protein orthologs identified with OrthoFinder [53]. The divergence of the monophyletic *Salvia* clade from the common ancestor of *Salvia* and the outgroup *Sesamum indicum* was estimated to occur ∼55 MYA based on previous studies [30, 45]. Among *Salvia* species with available whole-genome sequences, *S. divinorum* was most closely related to *S. splendens* and *S. hispanica*, which are sister to each other, and diverged from their common ancestor with *S. divinorum* at an estimated ∼8 MYA. The clade of *S. divinorum*, *S. hispanica* and *S. splendens* diverged from its common ancestor with the clade of *S. miltiorrhiza*, and *S. bowleyana* approximately 20 MYA. We estimate that the most distantly related *Salvia* species included in the analysis, *S. rosmarinus*, diverged from its common ancestor with other Salvia species ∼23 MYA.

**Figure 4.**
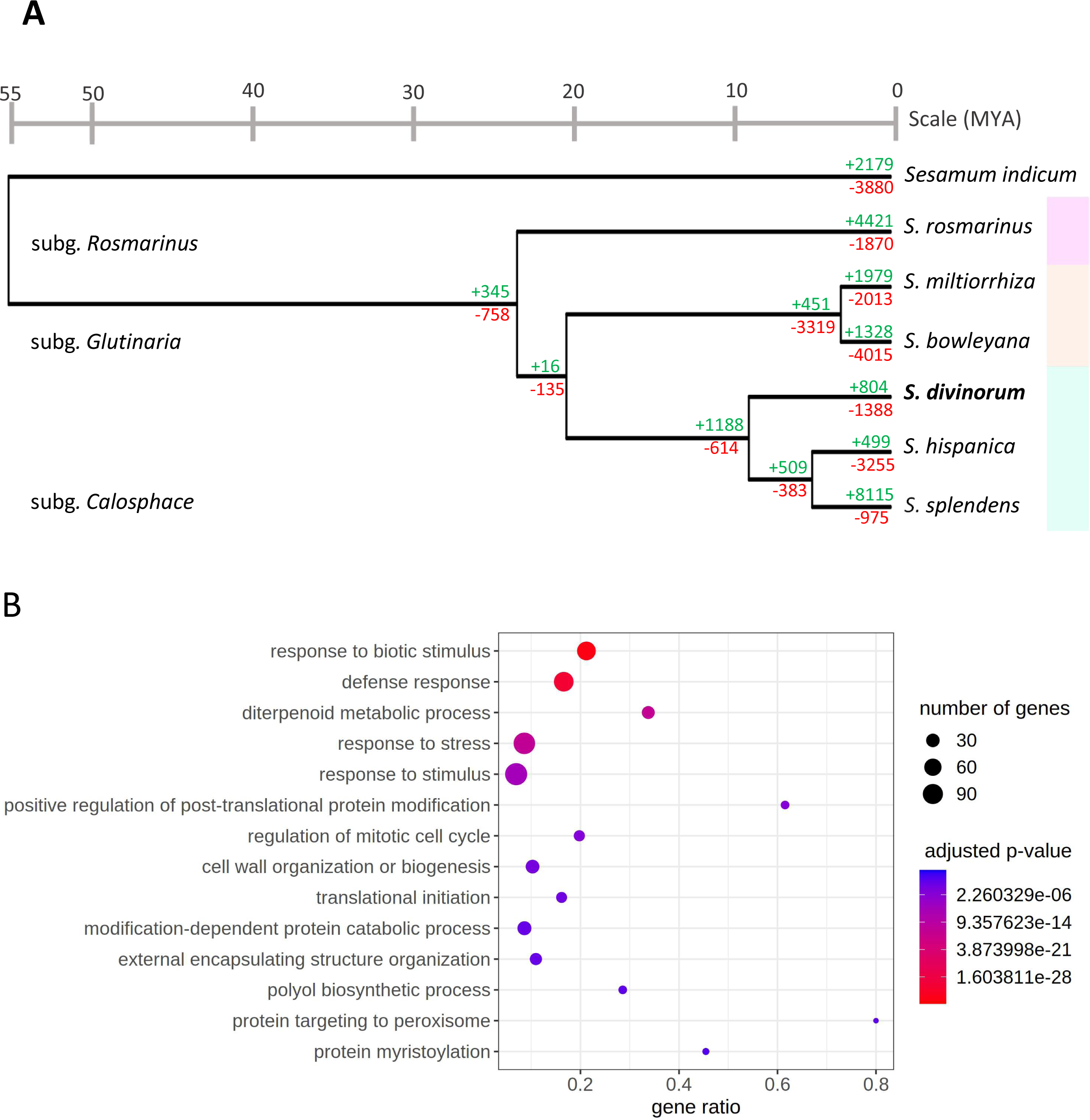
Genome evolution of *S. divinorum* and six related species and their divergence times. A, Maximum likelihood phylogeny constructed from a common set of protein orthologs (Emms and Kelly, 2019). Number of expanded and contracted gene families are shown in green and red, respectively, based on comparison of their orthogroups. The phylogeny was assembled using a JTT+CAT substitution model. *Sesamum indicus* served as outgroup. The colored boxes (right) indicate the Rosmarinus (pink), Glutinaria (orange), and Calosphace (green) subgenera. B, Significantly (p < 0.05) enriched GO Biological Process annotations in expanded gene families of *S. divinorum*.

A total of 27,469 orthogroups were identified in the 7 plant species, and gene family evolutionary analysis was perfomed using CAFE 5. Within the *Salvia* genus, *S. splendens* showed the greatest expansion of these gene families, likely owing to its polyploidization, while *S. hispanica* showed the least (Figure 4). Gene family expansion and contraction of *S. divinorum* was intermediate relative to other *Salvia* species, with 804 expansions and 1388 contractions. GO enrichment analysis demonstrated that significantly (p<0.05) expanded orthogroups in *S. divinorum* were enriched in genes involved in biological processes such defense against other organisms, stress response, and diterpene metabolism (Figure 4B). All 26 genes within an expanded family, annotated by the GO term ‘diterpene metabolic process’, are class I terpene synthases. Twenty four belong to the tps-*a* subclade with similarity to sesquiterpene synthase-like enzymes while the other 2 are kaurene synthases in the tps-*e*/*f* subclade (Supplementary Data Set 1A). Considering reports of sesquiterpene synthases acquiring plastid signal peptides and becoming active in diterpene pathways [54, 55], we used TargetP 2.0 [56] to predict the presence of N-terminal presequences in all terpene synthase proteins. However, none of the sesquiterpene synthase-like enzymes in expanded gene families were predicted to be plastid localized (Supplementary Data Set 1B). Enriched molecular functions within expanded orthogroups included binding to ADP, ions and heterocyclic compounds, as well as terpene synthase and O-methyltransferase activity (Supplementary Figure 5).

### Diterpene biosynthetic genes in the *S. divinorum* genome

We observed a total of 107 terpene synthase genes in the genome (TPS) that consisted of 84 tps-a/b/g, 12 tps-c (type II), and 11 tps-e/f [57–60] (Supplementary Data Set 1B). Among these, 17 putative diterpene synthases were detected which included 12 class II and five class I (Supplementary Data Set 1C). We constructed a neighbor-joining tree of diterpene synthases in the genomes of five *Salvia* species, using the ancestral bifunctional class I/ II diTPS ent-kaurene/kaurenol synthase of *Physcomitrium patens* as an outgroup (Accession: XP_024380398.1) (Figure 5).

**Figure 5.**
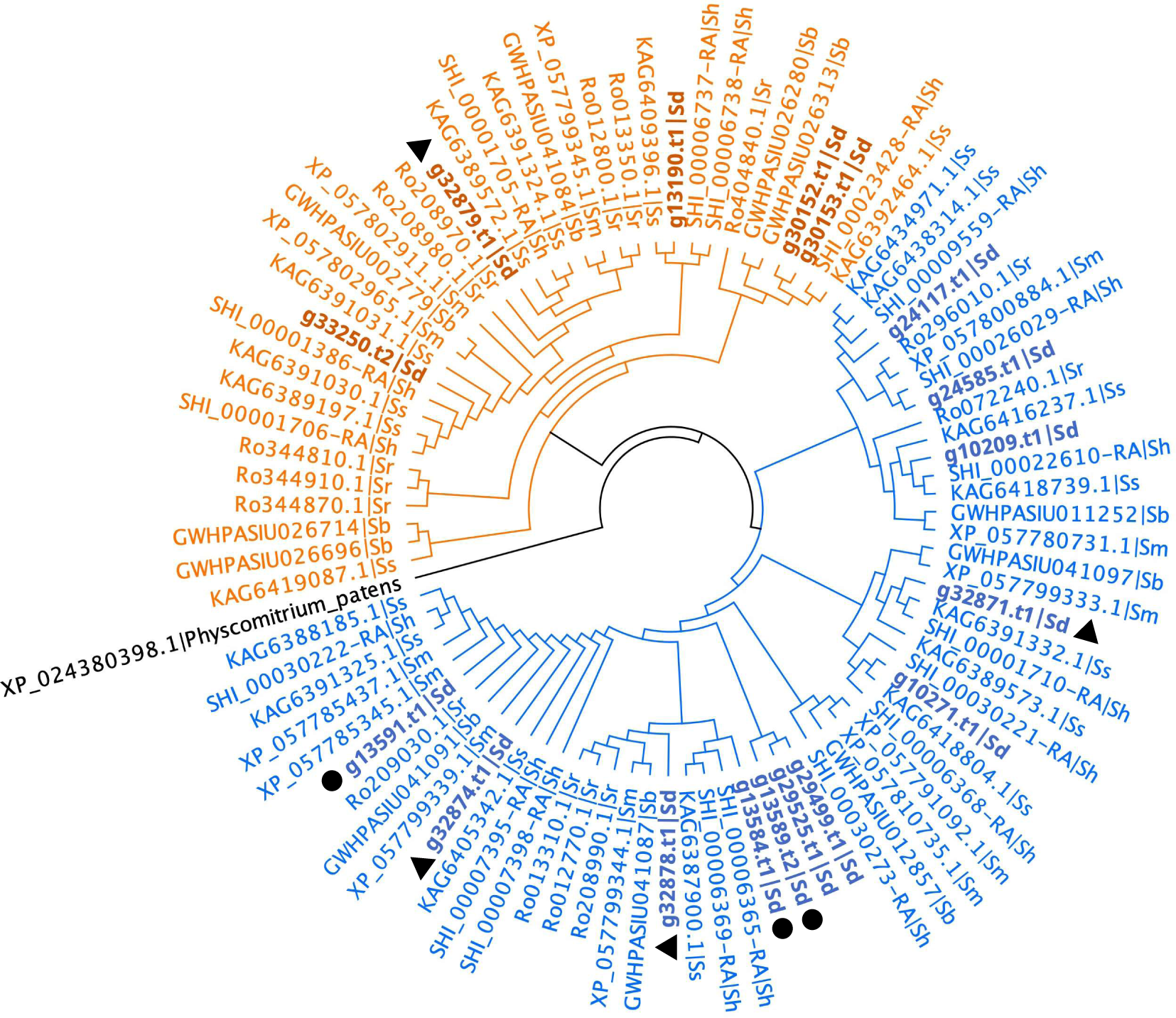
Neighbor-joining phylogeny of class II (blue) and class I (orange) diterpene synthases in *S. divinorum* (Sd; bold), *S. rosmarinus* (Sr), *S. miltiorrhiza* (Sm), *S. splendens* (Ss) and *S. bowleyana* (Sb). Rooted on the outgroup, ancestral bifunctional class I/ II diTPS *Physcomitrium patens* ent-kaurene/kaurenol synthase. Circles and triangles indicate presence in the contig 6 and contig 14 biosynthetic gene clusters, respectively as shown in Figure 6.

Cytochrome P450s (CYPs), which introduce oxygen functional groups into the core structure, are expected to play multiple roles in the SalA pathway. A total of 392 CYP genes were detected (Supplementary Data Set 2) that encode 422 predicted transcripts including isoforms (Supplementary Data Set 1D). Nine CYP families were represented (Supplementary Figure 6). The majority of CYPs were in the CYP71 clan, which is known for its involvement in diterpenoid biosynthesis [5].

We examined the genome for additional enzymes likely to be involved in the SalA pathway, predicted 69 putative alcohol dehydrogenases (ADHs) (Supplementary Data Set 1E), 39 SAM-dependent O-methyl transferases (OMTs) (Supplementary Data Set 1F), and 105 BAHD acyl transferases (BATs) (Supplementary Data Set 1G). These classes of enzymes were found on each chromosome in the *S. divinorum* genome, showing no strong positional bias beyond the large-scale tendency for genes to be at the ends of chromosomes (Figure 2).

There is evidence in the mint family that genes in the same secondary metabolic pathway cluster together in the genome [37]. Two BGCs identified on contigs 14 and 6 were enriched in diterpene synthases and CYP450s (Figure 6; Supplementary Data Sets 1H and 1I). Among the CYPs in the BGC on contig 14 was crotonolide G synthase [22], an enzyme highly expressed in glandular trichomes which catalyzes the formation of crotonolide G (dihydrofuran neoclerodane) from (-)-kolavenol. Other CYPs in the cluster included one CYP76B (g32876), one CYP71BE (g32875) and one CYP71D (g32873).

**Figure 6.**
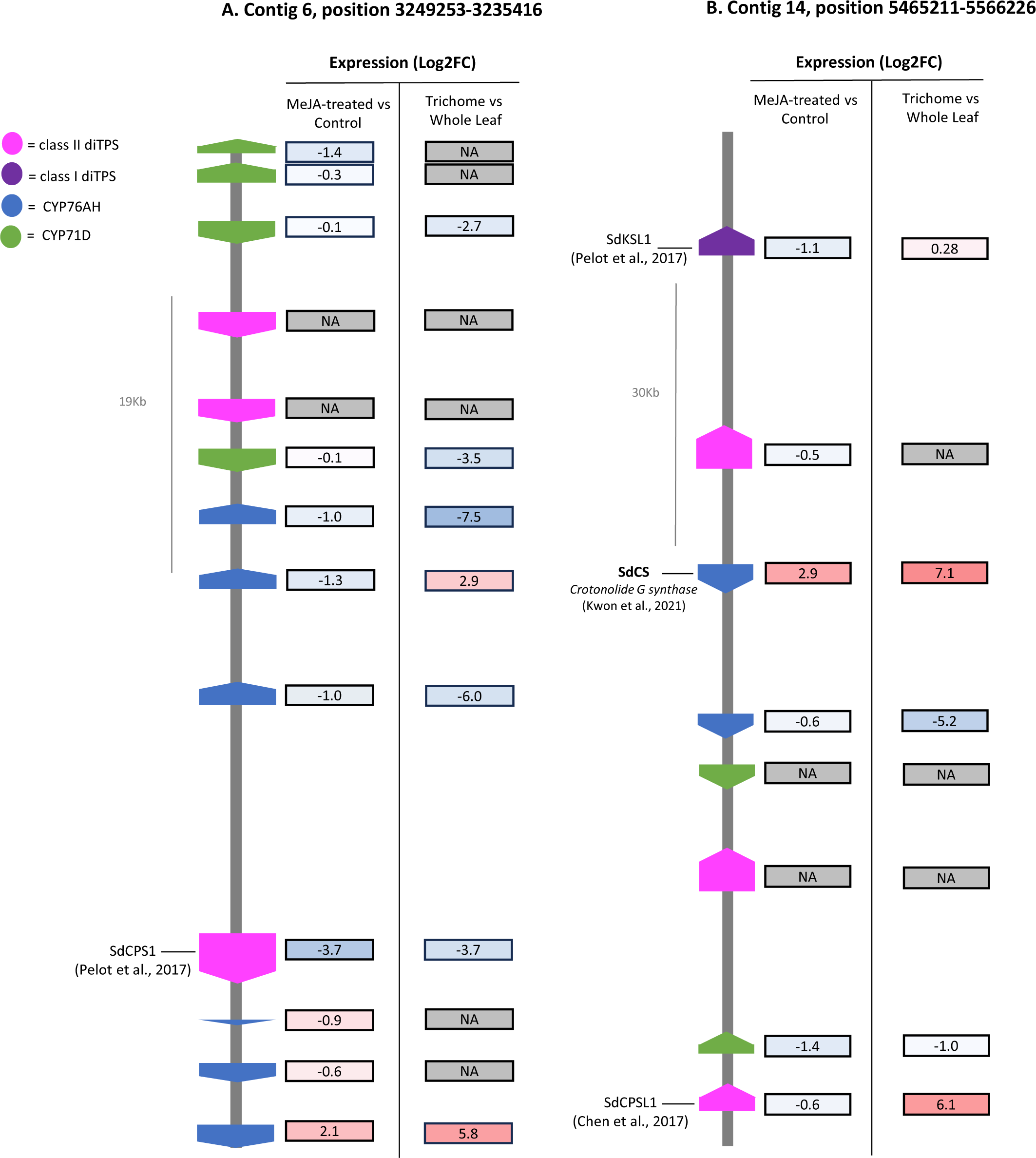
Diterpene biosynthetic gene cluster on contigs 6 (a) and 14 (b) in the *Salvia divinorum* genome. Genetic map shows position and class of diterpene biosynthetic genes. MeJA-treated vs control represents the log2fold change in expression of methyl jasmonate-treated plants relative to control plants, while trichrome vs whole leaf shows log2fold change in expression in trichomes [20] relative to whole leaves. Colors are scaled independently for each RNA-sequencing data set and depict the magnitude of expression change relative to all genes in enzyme classes postulated to participate in SalA biosynthesis (Supplementary Data Set 1J).

Putative diterpene synthases identified in BGCs included six class I and one class II enzyme. Despite clustering with crotonolide G synthase on contig 14, the class I diTPS kaurene synthase-like 1 (SdKSL1) was reportedly unable to convert CLDP into (-)-kolvaenol for SalA biosynthesis [21]. The class II diTPS, copalyl-diphosphate synthase-like 1 (SdCPSL1), located in the same BGC, was previously predicted not to produce (-)-kolavenol diphosphate [(-)-KPP] in the SalA pathway based on its similarity to class II diTPSs that synthesize (+)-copalyl diphosphate [21]. However, copalyl-diphosphate synthase 1 (SdCPS1), found in the BGC on contig 6, grouped phylogenetically with (+)-KPP synthases (Figure 6; g13584) but was shown to have (-)-KPP synthase activity [20]. Though this finding provides incentive for functional investigation of SdCPSL1, SdCPS1 is predominantly expressed in roots and is not thought to be involved in SalA biosynthesis [20]. Other diterpene synthases and CYPs within the biosynthetic gene clusters on contigs 6 and 14 have not yet been characterized, but do not appear to be highly enriched in trichomes (Supplementary Data Set 1J). Notably, copalyl-diphosphate synthase 2 (SdCPS2), which catalyzes the first step in the proposed SalA biosynthetic pathway [20, 21], is isolated on chromosome 4, and no coding sequence for biosynthetic enzymes was found within 200 Kb of this gene.

### Glandular trichome-specific and MeJA-responsive transcript expression

The assembled *S. divinorum* genome allowed us to profile the enrichment of RNA transcripts putatively involved in SalA biosynthesis in enriched tissue (glandular trichomes) or in response to induction from the defense hormone MeJA. RNA-seq analysis from all tissue sources corroborated 93% of the predicted proteins. In response to MeJA induction, 7% of annotated TPS genes were significantly upregulated (p<0.05); however, none of them was predicted to be a diterpene synthase (Supplementary Data Set 1B, 1C, and 1J). Among CYP450 genes, 7% were also enriched following MeJA induction. Significant upregulation was also observed for 9, 14, and 15% of ADH, BAT and SAM-dependent OMT genes, respectively (Supplementary Data Set 1E, 1F, 1G, and 1J). Confirming the effectiveness of MeJA treatment, we observed the upregulation of fatty acid hydroperoxide lyase (g11225), allene oxide cyclase (g206), and the JASMONATE ZIM-domain (JAZ) transcriptional repressor, JAZ1 (g116446), which aligns with the common response in MeJA-treated plants. Consistently, the BTB/POZ domain-containing, nonexpressor of pathogenesis-related genes 1 (NPR1; g30536), a key activator of salicylic acid (SA)-mediated immune responses was significantly downregulated in MeJA treated plants .[61]

Expression of genes in diterpene metabolic gene clusters was not clearly affected by MeJA treatment. Among genes in the two identified clusters (Figure 6), only crotonolide G synthase on chromosome 14 (which synthesizes crotonolide G in the SalA pathway) was significantly upregulated (Log2FC = 3.0; p <0.05). However, three CYP450 genes in the cluster on contig 6 were significantly downregulated (Log2FC = -3.7, -1.3, -1.0; p < 0.05). Despite induction of crotonolide G synthase and another CYP450 implicated in the SalA pathway (CYP728D26 or g5204) [23] with Log2FC = 2.2 (p ∼ 0.01), expression of SdCPS2, the class II diTPS catalyzing the first committed step in the SalA pathway, was not significantly altered by MeJA (Log2FC = 0.6). Among the four genes with an established or putative role in the SalA pathway, only crotonolide G synthase demonstrates statistically significant induction by MeJA (Supplementary Table 1), indicating that not all enzymes in the SalA pathway are uniformly co-expressed following MeJA treatment.

Similarly, clustered diterpene biosynthetic genes were not uniformly enriched in the trichomes (Supplementary Data Set 1J). The majority of genes in these clusters were downregulated in trichomes. RNA-seq analysis showed that aside from aside from crotonolide G synthase, no CYP in this cluster was upregulated in trichomes (Supplementary Data Set 1J). Two CYP76Bs of unknown function in the contig 6 BGC were enriched in trichomes (g13581 and g13586; Log2 FC > 2). However, genes implicated in the SalA pathway, regardless of position in the genome, were uniformly highly expressed in trichomes (Supplementary Table 1). These results corroborate the observations above that SalA pathway genes are neither uniformly induced by MeJA treatment nor clustered together in the genome.

## Discussion

### Most SalA genes are not associated with BGCs

*S. divinorum* is a medicinal plant with a long history in shamanistic healing ceremonies of the Mazatec people of Oaxaca, Mexico. Its use in such rituals is due to the psychoactive diterpene SalA, which accumulates in glandular trichomes of *S. divinorum* aerial tissues. Despite its potential medical applications in the treatment of chronic pain, addiction, and depression, only two steps in the SalA biosynthetic pathway are currently known. Here we have generated a high-quality reference genome sequence which will serve as a guide to facilitate the complete elucidation of the SalA pathway and better understand the unique secondary metabolism of the *Salvia* genus.

Our genome assembly is highly contiguous with N50 values that mark a substantial improvement upon other recently published *Salvia* genomes [30, 33]. The high BUSCO completeness scores of this assembled genome (98.4% and 98.2% at the genome and protein levels, respectively) also signify that the assembly is of high quality with significant value for the goal of elucidating the SalA pathway. This assembled genome sequence is among the highest contiguity of any *Salvia* genome published to date.

The repeat content of *S. divinorum* was noticeably higher than that of *S. splendens* (47.99%) [33], *S. miltiorrhiza* (between 54.44% [28] and 56.65% [45] ), or *S. bowleyana* (58.7%) [29] but similar to *S. officinalis* (61.67%) [32]. Another report placed the repeat content of *S. miltiorrhiza* even closer to *S. divinorum* (64.84%) [27]. Within the genus, the genome of *S. rosmarinus* currently possesses the highest repeat content at 72.72% [30].

Genes for diterpene metabolism have been known to form clusters in the Lamiaceae [37], and a recent comparison of several *Salvia* genomes confirmed that BGCs are a typical feature of genomes in this family [30]. In the *S. divinorum* genome, we identified two BGCs located on contig 6 and contig 14, each enriched in CYPs and diterpene synthases. Notably, the BGC on contig 14 contains crotonolide G synthase, the CYP which forms the dihydrofuran ring on C16 of the neoclerodane backbone in salvinorin biosynthesis [22]. In addition to crotonolide G synthase, sequences for three other CYPs were identified in this BGC, but only two were expressed. CYPs within the cluster were differentially regulated, as only crotonolide G synthase showed a significant increase in expression after MeJA treatment, while the other two expressed CYPs were slightly downregulated. Similarly, two of the three diTPSs in this cluster were expressed, and both showed slight downregulation in response to MeJA. The other identified BGC, on contig 6, contained 10 CYPs, 3 of which were significantly downregulated by MeJA, while one was strongly upregulated in response to MeJA (g13581; Log2FC = 2.12; p=0.07). This transcript was also significantly enriched in trichomes (Log2FC = 6.14) and therefore represents a promising candidate for involvement in the SalA pathway. Only one diTPS in this cluster (SdCPS1; g13584) was expressed, and it was significantly downregulated both in trichomes (Log2FC = -3.74) and after jasmonate treatment (Log2FC = -3.71).

Genes in BGCs are often co-regulated. For instance, the BGCs involved in paclitaxel synthesis in *Taxus chinensis* showed tightly coordinated expression patterns in tissues and in response to jasmonate [62]. Differential regulation of genes within diterpene gene clusters has also been observed in *S. officinalis* [32]. In the latter, a single BGC contained two pairs of class I and class II diTPS, and each was independently co-expressed in roots and shoots despite membership in separate diterpene biosynthetic pathways in these organs [32]. We speculate that in *S. divinorum*, the pair of class I (SdKSL1) and class II diTPSs (g32873) in the BGC of contig 14 may be part of a diterpenoid pathway in roots, as SdKSL1 was shown to be expressed in roots [20] and the pair showed coordinated downregulation in response to MeJA (Supplementary Data Set 1J). This would suggest that crotonolide G synthase may have dual roles in biosynthesis of SalA and root diterpenoids, as has been previously documented for CYP76AHs in the BGC of *S. officinalis* [32]. SdCPS2 in contrast, which catalyzes the committing step in the SalA pathway, is isolated on chromosome 4. These observations lead us to conclude that genes encoding enzymes in the SalA pathway are not exclusively found in diterpene metabolic gene clusters. Similarly, paclitaxel biosynthesis involves two discrete BGCs as well as many genes not found in BGCs [62]. But unlike the known genes for SalA biosynthesis, most are found in a small region of a single chromosome. Thus, the organization of clerodane biosynthetic genes in *S. divinorum* more closely resembles that of skullcap, wherein many clerodane biosynthetic genes are not associated with BGCs but appear instead to be the products of gene duplications of genes that are [35]. This is consistent with the assertion that BGCs may serve as toolboxes for recruitment of new catalysts in emergent secondary metabolic pathways.

### Insights into kolavenol formation

Class II diTPS enzymes catalyze the rearrangement of geranylgeranyl diphosphate (GGDP) with retention of the diphosphate group. They are all derived from an ancestral diTPS sequence which produces *ent-*copalyl diphosphate, a precursor to gibberellin [59]. Examples which produce copalyl diphosphate, clerodienyl diphosphate, and isokolavenyl diphosphate have been described [19, 63, 64]. However, no class I diTPS participating in a clerodane diterpenoid biosynthetic pathway has been identified to date [35]. The conversion of KDP to kolavenol (Figure 1; step 2) therefore remains unclear. Several possibilities have been proposed. Extensive transcript mining has not thus far not produced a type I diTPS that carries out the conversion of KPP to kolavenol. Three type I diTPS (SdKSl1-3) were previously identified [21]. We found evidence for two additional type I diTPS sequences in the *S. divinorum* genome. One does not appear to be expressed (g33250). The other, g30152, is 97% identical to SdKSl and shows low expression relative to other diTPSs. However, it does appear to be predominantly expressed in trichomes, warranting future functional characterization.

An alternative explanation for conversion of KDP to kolavenol is the Nudix hydrolase family of enzymes, which have found an unexpected place in terpene metabolism over the past several years [65–69]. Nudix hydrolases act on diphosphate substrates through a phosphohydrolase mechanism, removing a single phosphate group [70]. Although mostly involved in housekeeping metabolism, one subclade of Nudix hydrolases preferentially acts on terpenoid substrates [71]. It has been suggested that conversion of prenyl diphosphate substrates to their monophosphate analogs increases their susceptibility to non-specific phosphatases to yield the corresponding alcohol [65]. We searched the *S. divinorum* genome for Nudix hydrolases which might convert KPP to KP and identified three genes which clustered within the terpene modifying subclade (g224, g225 and g7372) (Supplementary Figure 7 and Supplementary Data Set 1K). Two Nudix sequences, g224 and g7372, were highly upregulated following MeJA treatment (Log2FC=2.07 and 2.16, respectively), while no other Nudix hydrolase showed a Log2FC greater than 1.17 (Supplementary Data Set 1J). Additionally, g225 was highly expressed in glandular trichomes relative to whole leaves (Log2FC = 6.67), and predicted to have a chloroplast transit peptide. Together, these results provide a strong rationale for further investigating a possible role for Nudix hydrolases in the SalA pathway.

## Conclusions

We have presented the first assembly and annotation of the *S. divinorum* genome. This high quality genome will serve as a valuable resource for future studies aiming to elucidate steps in the biosynthetic pathway of the potent *K*-opioid receptor agonist, salvinorin A. The sequenced genome will provide a framework for leveraging the medicinal potential of *S. divinorum*.

## Methods

### Genomic DNA isolation, sequencing, and assembly

High molecular weight DNA was extracted from frozen, dark-treated *S. divinorum* leaf tissue using a standard cetyltrimethylammonium bromide (CTAB) / phenyl:chloroform:isoamyl alcohol method high molecular weight (HMW). Frozen *S. divinorum* leaves stored at -80 °C were obtained from the University of Toronto -Mississauga greenhouse collection. DNA was extracted from frozen, dark-treated *S. divinorum* leaf tissue using a standard cetyltrimethylammonium bromide (CTAB) method. Briefly, ∼0.5 g of ground leaf tissue was suspended in 2 mL of 65° C CTAB buffer with 5 μL each of proteinase K (20 mg·mL^-1^ stock) and RNAse A (100 mg·mL^-1^ stock), and incubated at 65° C for 30 minutes. After cooling to room temperature, 2 mL phenol:chloroform:isoamyl alcohol (25:24:1) were added to the solution and mixed by inversion for 5 minutes at 4° C, then centrifuged at 1500 × g for 5 minutes. The aqueous phase was isolated, and re-extracted as above, this time with an equal volume of chloroform:isoamyl alcohol (49:1). The aqueous phase was collected, and DNA was precipitated by the addition of 2× volume of ice cold 100% ethanol. DNA was pelleted by centrifugation at 20000 × g for 1 minute, and washed with 70% ethanol, before a final resuspension in ultrapure ddH2O. A 1× volume of SPRIselect bead-based reagent (Beckman Coulter, Inc.) was used for DNA cleanup and purification prior to library preparation. SMRTbell library preparation was performed by The Centre for Applied Genomics at SickKids Hospital (Toronto, Canada) and Pacbio Hifi sequencing was conducted on the Revio SMRTcell platform.

To estimate genome size, Jellyfish v2.2.10 [72] was used to count canonical (-C) 21 bp k-mers from the PacBio HiFi reads, and the resulting k-mer frequency histogram was provided to GenomeScope [73] to fit a model for the prediction of the genome length, heterozygosity, and repetitiveness.

Raw Pacbio HiFi reads were assembled using two validated softwares: (1) hifiasm v0.16.1 [39] and (2) Flye v2.8.1 [74] with default parameters. The haploid assembly generated by hifiasm (458 contigs; N50=41Mb) was more contiguous than the Flye assembly (1760 contigs; N50=11.1 Mb), and was therefore selected for further processing and subsequent analyses.

To remove erroneous allelic duplication and low coverage contigs from the assembly purge_dups v1.2.5 [40] was used after self-alignment using Minimap2 v2.22 [75]. A single circular chloroplast chromosome was assembled from HiFi reads using ptGAUL [41]. ptGAUL uses a reference chloroplast genome from a related species to select HiFi reads originating from plastid DNA, downsamples the reads to ∼50x coverage, and assembles them with Flye. We used a reference chloroplast genome of *S. madrensis,* a close relative of *S. divinorum*, with an estimated chloroplast genome size of 150 Kb. A similar approach was used to assemble the mitochondrial genome; we first mapped HiFi reads to a concatenated FASTA file of *S. splendens* and *S. miltiorrhiza* mitochondrial genomes using Minimap2, and filtered mappings for length >1,000 bp and number of residue matches/length >0.7. After downsampling the mapped reads to ∼50x coverage (using an estimated mitochondrial genome size of 300 Kb), we assembled them with hifiasm, which produced two non-redundant circular mitochondrial DNA chromosomes. To filter redundant organellar DNA contigs from the final assembly, we mapped assembled contigs to the organellular genomes and removed those which had a mapping length of >99% contig length. To assess completeness, the assembly was queried against the embryophyta_odb10 database using BUSCO v5.6.1 [76] in genome mode.

### Structural annotation

For improved gene-model prediction, we first softmasked the assembled genome. A species-specific repeat library was generated using RepeatModeler v2.0.5 [77] and merged with eudicot repeat database from RepBase v. 20240126. To softmask repeats, the joined library was provided as input to RepeatMasker v4.1.6 [78].

To predict gene models, we combined evidence from species-specific RNA-seq and proteins from related plant species. First, we ran BRAKER v3.0.8 [79] in RNA-only mode using raw *S. divinorum* RNA-seq reads obtained from whole leaves and isolated trichomes [20] (Supplementary Data Set 1L), as well as the control and methyl jasmonate (MeJA)-induced leaves generated in this study. Next, we created a protein database consisting of all Viridiplantae proteins from OrthoDB v11 [80] merged with Lamiaceae proteins downloaded from UniProt (20240220). The resulting 5,401,793 plant proteins were used as input to BRAKER, running in protein-only mode. We then combined evidence from protein-only and RNA-only BRAKER modes using TSEBRA [79] and applied the script `agat_convert_gxf2gxf.pl` from AGAT v1.0.0–pl5321hdfd78af_0 [81] (https://zenodo.org/records/5834795) to create a GFF3 file from the TSEBRA-generated GTF. To evaluate gene prediction, we queried the predicted protein sequences against embryophyta_odb10 using BUSCO v5.6.1 in protein mode.

We used GeSeq to annotate the organellular genomes [82] and produced graphical maps using OGDRAW v1.3.1 [83]. We used BLAT search for prediction of coding sequence, rRNA and tRNA, with additional tRNA annotation by tRNAscan-SE v2.0.7. We provided GeSeq with the reference mitochondrial genome of *Arabidopsis thaliana*, and reference plastid genomes of *S. splendens*, *S. miltiorrhiza*, and *S. hispanica* (Supplementary Data Set 1L). For plastome annotation, we used profile searches of Embryophyta chloroplasts by HMMER and Chloe v0.1.0 to supplement BLAST and predictions of coding sequences and rRNA.

#### Functional annotation and biosynthetic gene cluster analysis

Functional annotation of genes was performed using eggNOG-Mapper v2 [47] and InterProScan v5.66-98.0 [46] which queried 14 protein databases and assigned Gene Ontology (GO) terms. EggNOG-Mapper and InterProScan outputs were used as input to funannotate v1.8.14 [48], which combined the annotations and queried additional databases CAZy, MEROPS and BUSCO. As described by Santangelo et al., [84] (https://github.com/James-S-Santangelo/dcg/tree/main), if a protein was annotated as ‘hypothetical protein’, but was assigned a fully resolved enzyme commission (EC) number, the annotation was replaced with the EC number’s product in the ExPASSY Enzyme database [85].

Based on the expected enzyme classes participating in the SalA pathway (Figure 1), candidate terpene synthases, alcohol dehydrogenases, CYP450s, BAHD acyltransferases, SAM-dependent O-methyltransferases and NUDIX hydrolases in *S. divinorum* were identified using representative Pfam and InterPro domains (Supplementary Data Set 1L). CYPs were putatively classified into clans using a protein-protein BLAST v2.5.0 [86] search of predicted *S. divinorum* CYPs against a database of 9739 plant CYPs previously classified by Dr. David Nelson (https://drnelson.uthsc.edu/plants) (Supplementary Data Set 2). To identify putative diterpene synthases, we performed a BLAST search a class II diTPS, SdCPS2, (Accession: A0A1S5RW73.1) class I diTPS, SmKSL1 (Accession: XP_057799345.1), and a bergamotene (sesquiterpene) synthase (Accession: XM_048123031.1) and against a database of all predicted *S. divinorum* proteins. To obtain the list of diTPSs, we filtered genes in which the sesquiterpene synthase was the top hit, and applied a bit score cutoff of 45, which corresponded to a maximum e-value of 1.4e-21. Neighbor-joining trees were assembled using Geneious Prime 2021.2.2 (https://www.geneious.com). Presence of transit peptides was predicted using TargetP-2.0 [56].

To complement the class-specific search above, we further refined our list of candidate genes in the SalA pathway by searching for BGCs. We supplied PlantiSMASH [87] with our *S. divinorum* genome assembly fasta file and GFF file using default settings to identify putative BGCs that might contain genes for secondary metabolite biosynthesis.

### Genome evolution and gene family expansion and contraction

Published protein sequences of five *Salvia spp.* and *Sesasmum indicum* (Supplementary Data Set 1L) were obtained and used to make evolutionary inferences. After filtering protein sequences to include only the longest isoforms, orthogroups of *S. divinorum* and the six additional plants were identified using OrthoFinder v2.5.5 [53] with the following parameters: `- M msa -S diamond -A mafft -T fasttree`. A maximum likelihood species tree rooted on *Sesamum indicum* was inferred by FastTree [88] using a JTT+CAT substitution model. The tree was made ultrametric using the OrthoFinder script `make_ultrametric.py` with a root divergence time (`-r`) of 55 MYA, as determined previously [30].

Based on the obtained tree file and orthogroup gene counts, CAFE 5 [89] was used for determination of gene family expansions and contractions. Prior to analysis, the python script `clade_and_size_filter.py`, included with CAFE 5, was used to remove 22 of the 22,234 total orthogroups in which a single species had >100 gene copies. Significance of gene family expansions and contractions was determined using a threshold of P<0.05. Gene ontology (GO) enrichment analysis was performed for genes within significantly expanded gene families in *S. divinorum* using GOATOOLS [90] with GO annotations obtained from InterProScan [46].

We assessed pairwise synteny between *S. divinorum* and *S. splendens* using GENESPACE v1.3.1 [51] which employs MCScanX [91] to detect regions with conserved gene order and OrthoFinder [53] to identify paralogs and orthologs within these syntenic blocks.

### RNA-seq analysis

RNA transcript enrichment was calculated for glandular trichomes and, on the whole leaf level, in response to induction by MeJA. In the first experiment, RNA from isolated trichomes was compared to that of leaves. Preparation of trichome-enriched RNA-Seq analysis was performed as previously described [20] and deposited in NCBI (Accession number: SRR3716680). Transcripts in trichomes vs leaves was available only as a single data set, and fold change calculations were based on sequencing a single library of each type.

In the second, *S. divinorum* plants were sprayed with aqueous 1 mM MeJA containing 0.1% (*v*/*v*) Tween-20 (or a control solution containing only Tween-20). The total application volume was 10 mL per plant. Control and experimental plants were covered with plastic bags for 4 hours after spraying. Young leaves of approximately the same size were collected for RNA preparation three weeks after treatments. Total RNA samples were prepared by Trizol (Sigma-Aldrich) and further purified by EZNA RNA purification kit (Omega Biotech), followed by evaluations of RNA integrity by Bio-analyzer (Agilent Technologies). Libraries of four biological replicates were prepared and sequenced on an Illumina NovaSeq 6000 with paired-end reads of 100 bp at the McGill Genome Centre (Montreal, QC, Canada).

Reads from each library were trimmed using Trimmomatic [92] and mapped to the assembled *S. divinorum* genome presented here using STAR [93]. The resulting BAMs were post processed to sort reads by coordinate and add read groups using Samtools [94]. Differential gene expression analysis was conducted between Methyl Jasmonate-treated and control samples using the R package DESeq2 [95]. All code used for this analysis is provided in Supplementary Methods file 1.

To identify genes over-represented in glandular trichomes as candidates in SalA biosynthesis we analyzed a publicly available mRNA library sequenced with 454 GS FLX technology (SRA accession SRX1900686). We followed the same read trimming and mapping protocol outlined above. Exploratory expression analysis was conducted using the R package EdgeR [96] where differential expression can be characterized without replication using an exact test.

## Declarations

### Ethics approval and consent to participate

Not applicable

### Consent for publication

Not applicable

### Availability of data and materials

All data supporting the conclusions of this article are included within the article and its Supplementary files and datasets. The *S. divinorum* genome sequence described herein is available at NCBI (https://www.ncbi.nlm.nih.gov/) under the BioProject PRJNA1104206.

Detailed methods, including sample code for performing major analyses is available in Supplementary Methods File 2.

### Author Contributions

S.A.F. performed major experiments, analyzed data, and wrote the manuscript. R.W.N. and M.A.P. directed the research, analyzed data, and wrote the manuscript. M.K. performed minor experiments. D.K.R. assisted in preparation of the manuscript.

## Acknowledgements

The authors acknowledge a generous NSERC CGSM graduate scholarship supporting S.A.F.

---

**Supplementary Figure 1.**
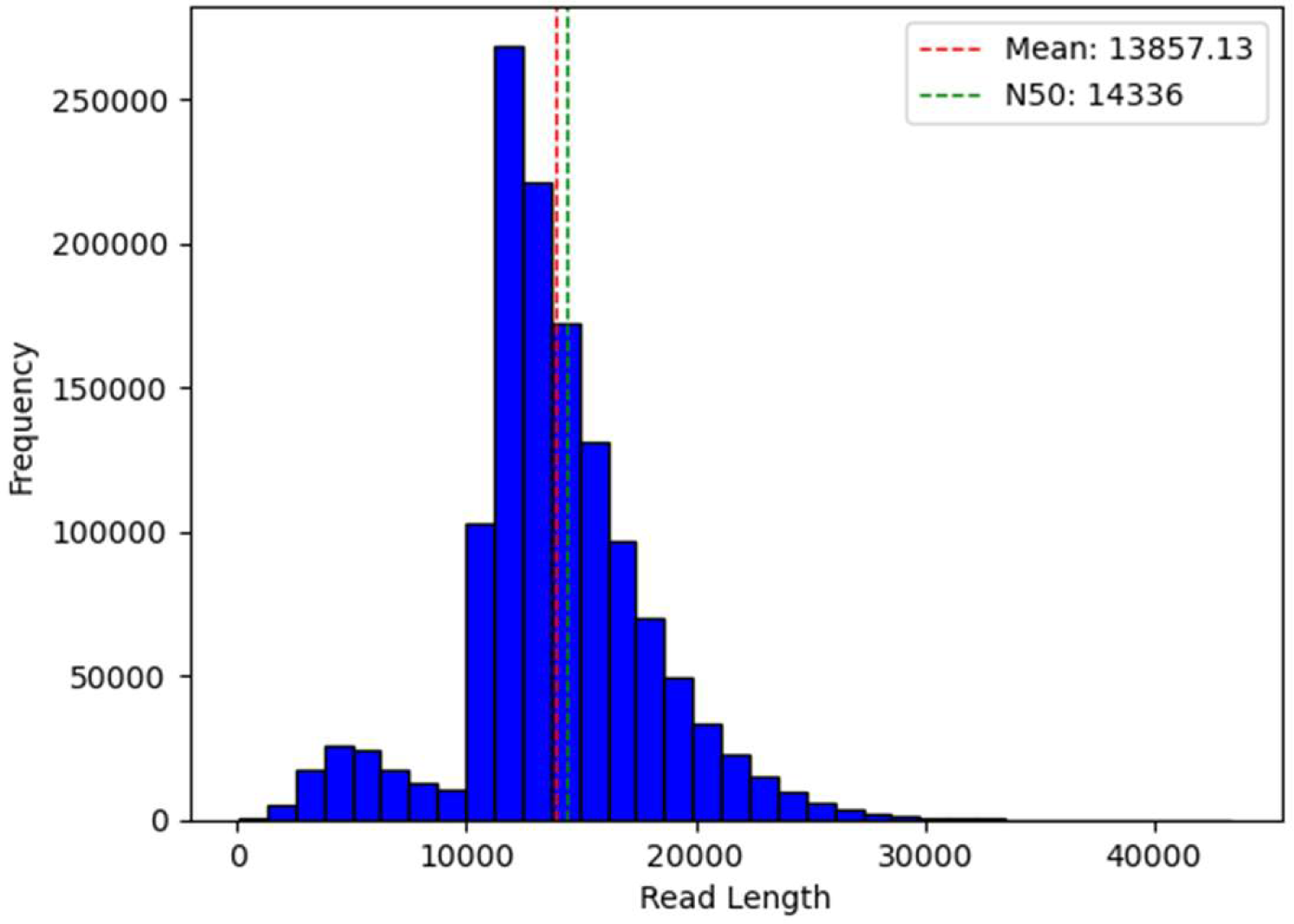
Distribution of pacbio hifi read lengths used in the genome assembly described in this study.

**Supplementary Figure 2.**
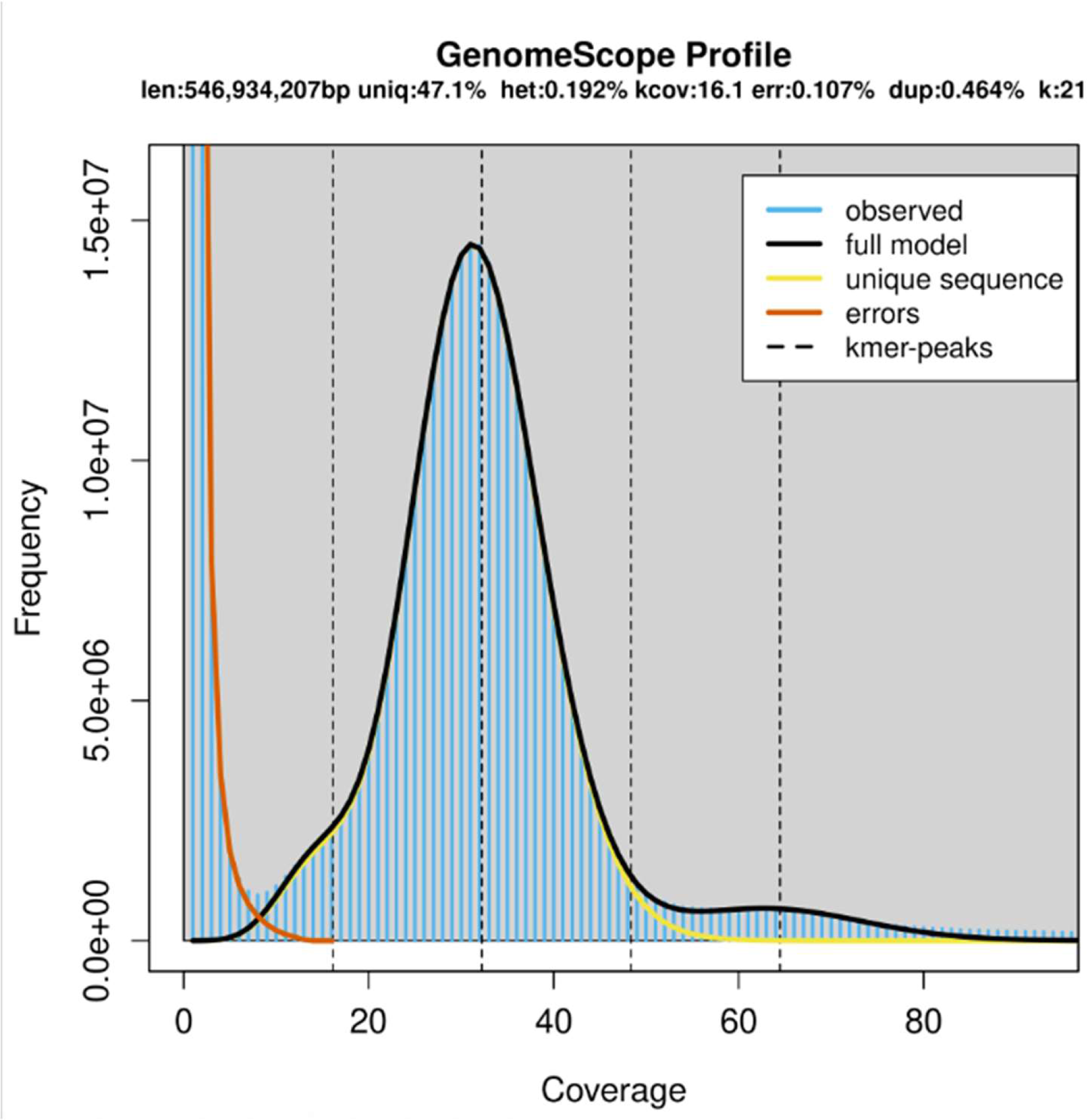
Genome size, heterozygosity, repeat content and coverage estimation by GenomeScope using 21-mer counting

**Supplementary Figure 3.**
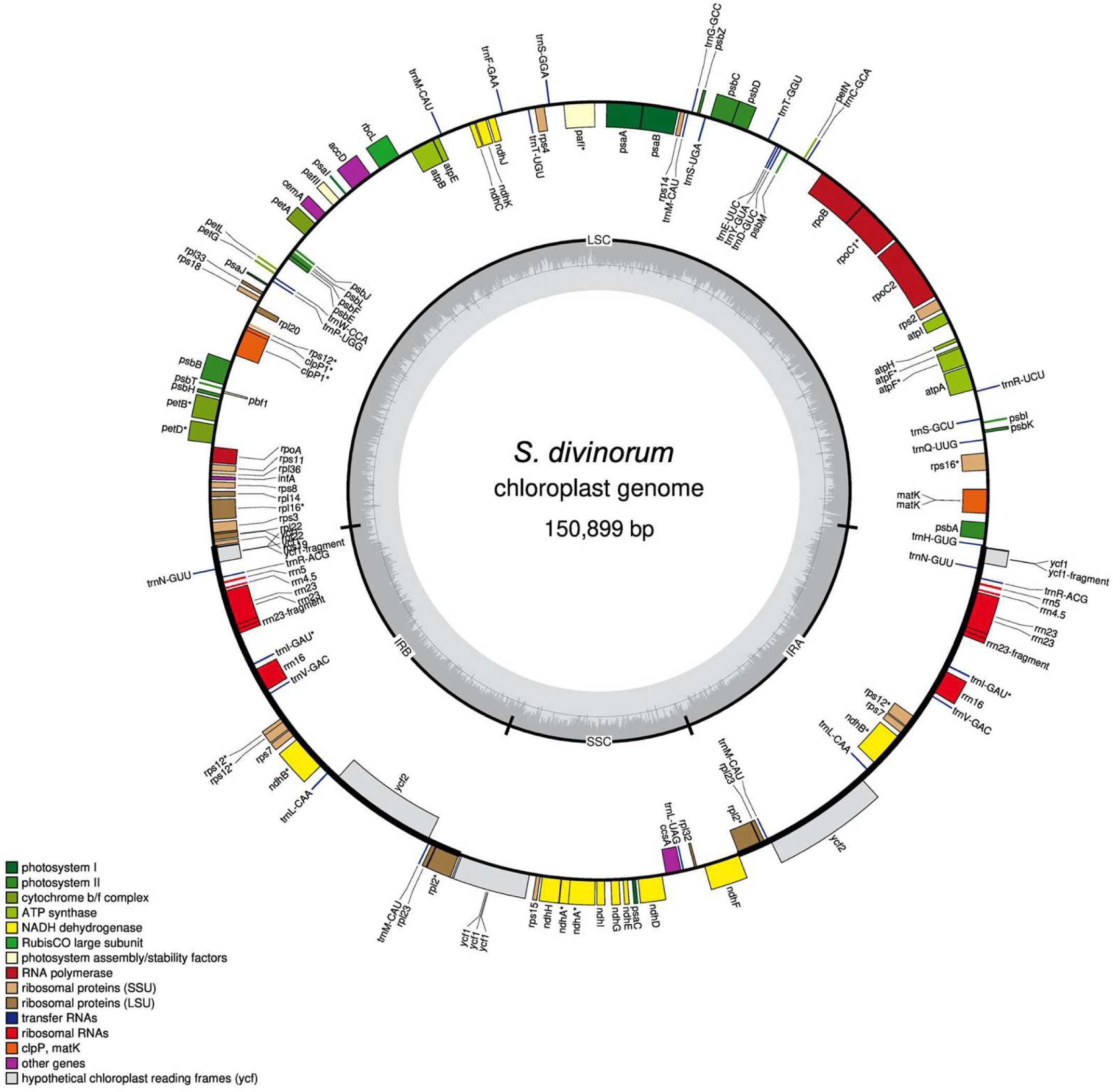
S. divinorum chloroplast genome structure and gene annotations. Inner grey ring displays GC content.

**Supplementary Figure 4.**
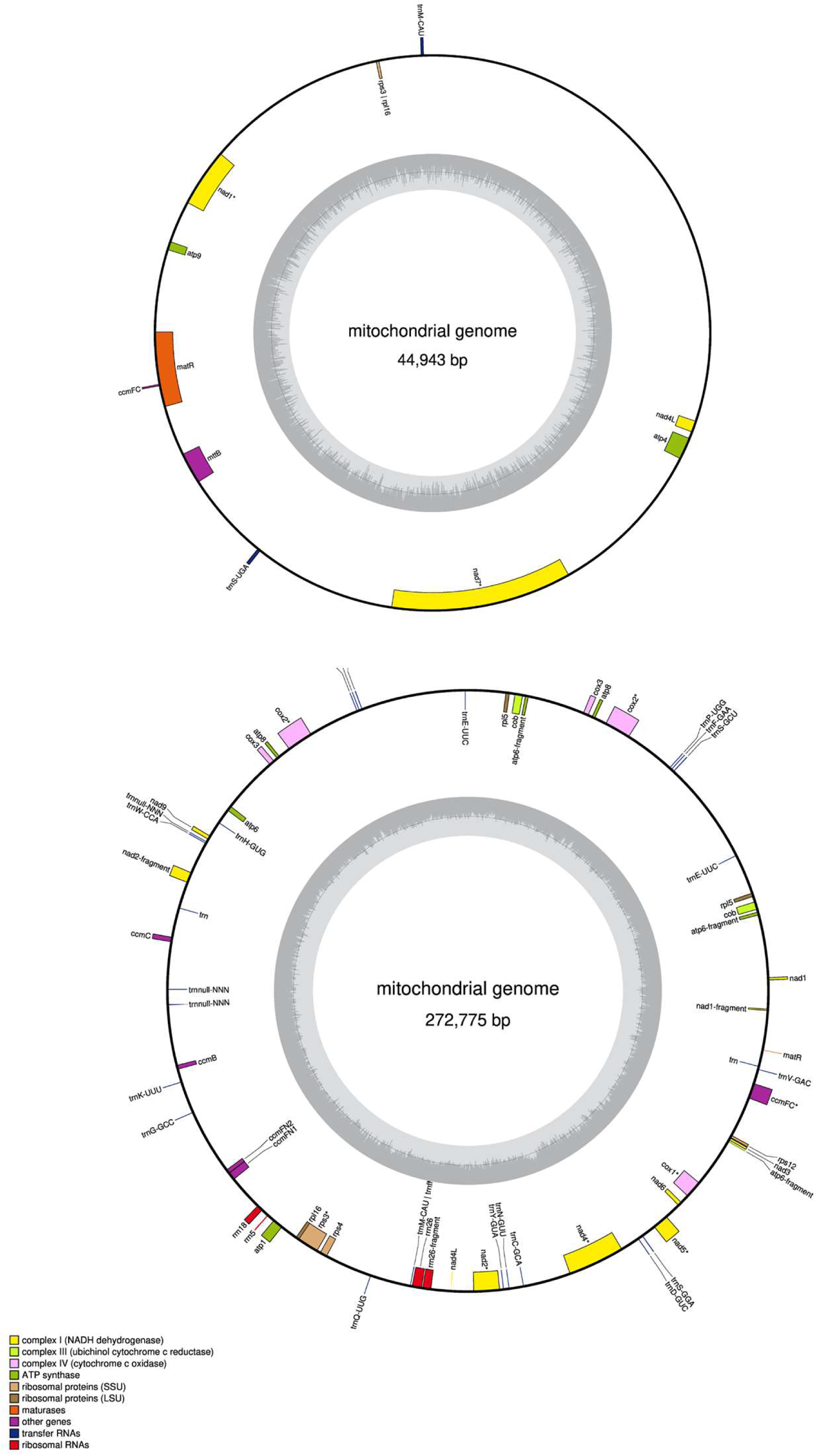
S. divinorum mitochondrial genome structure and gene annotations. Inner grey ring displays GC content.

**Supplementary Figure 5.**
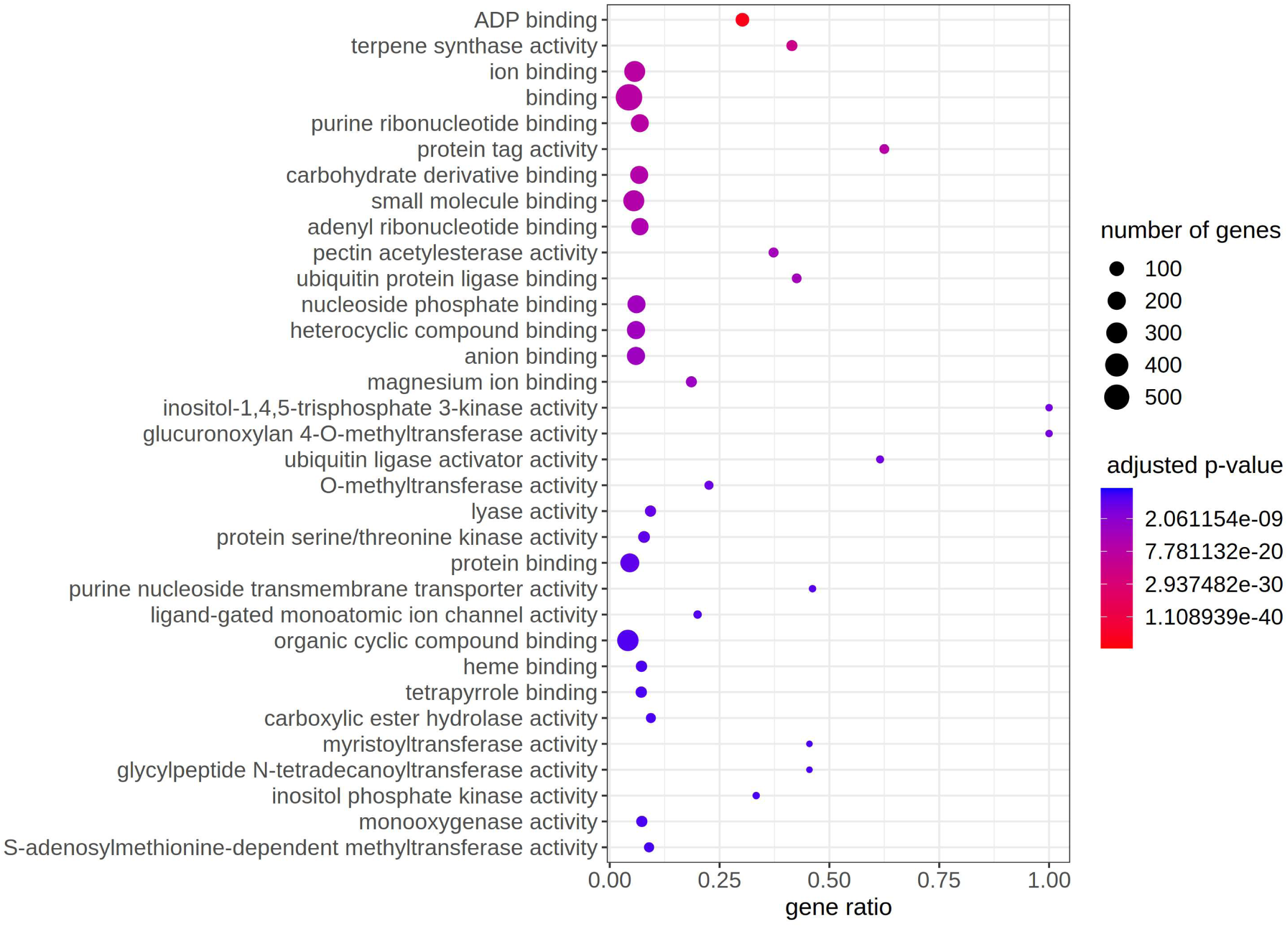
Significantly (p<0.05) enriched GO molecular function annotations in expanded gene families of S. divinorum.

**Supplementary Figure 6.**
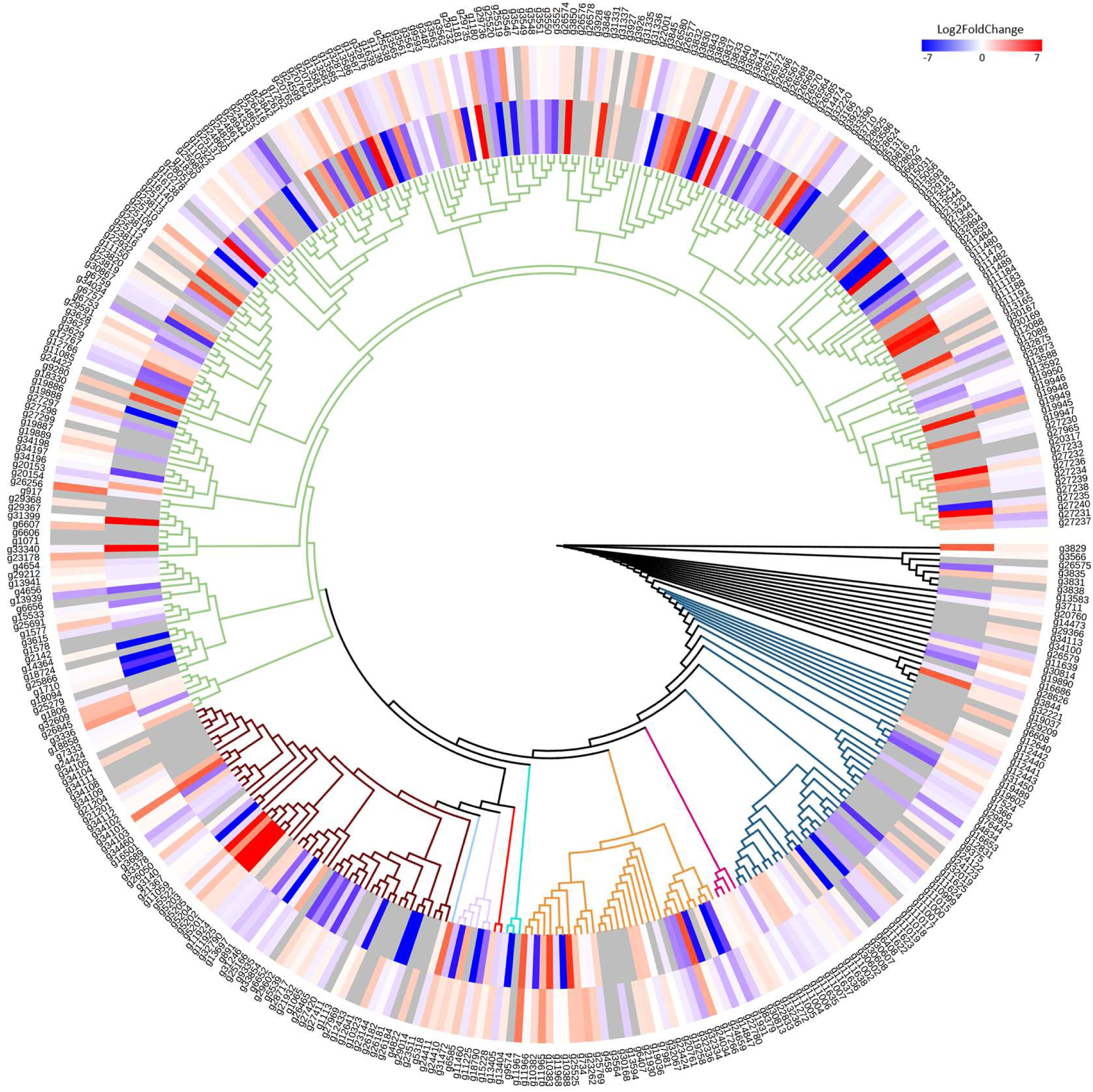
Neighbor-joining tree of S. divinorum cytochrome p450 genes. Branch colors depict CYP clans (green, CYP71; brown, CYP85; light blue, CYP710; light purple, CYP74; red, CYP51; turquoise, CYP711; orange, CYP96; purple, CYP97; navy, CYP72; black, unclassified). Inner heatmap ring represents log2 fold change of expression in trichomes compared to whole leaf; outer heatmap represents log2fold change in expression in methyl jasmonate induced compared to control leaves.

**Supplementary Figure 7.**
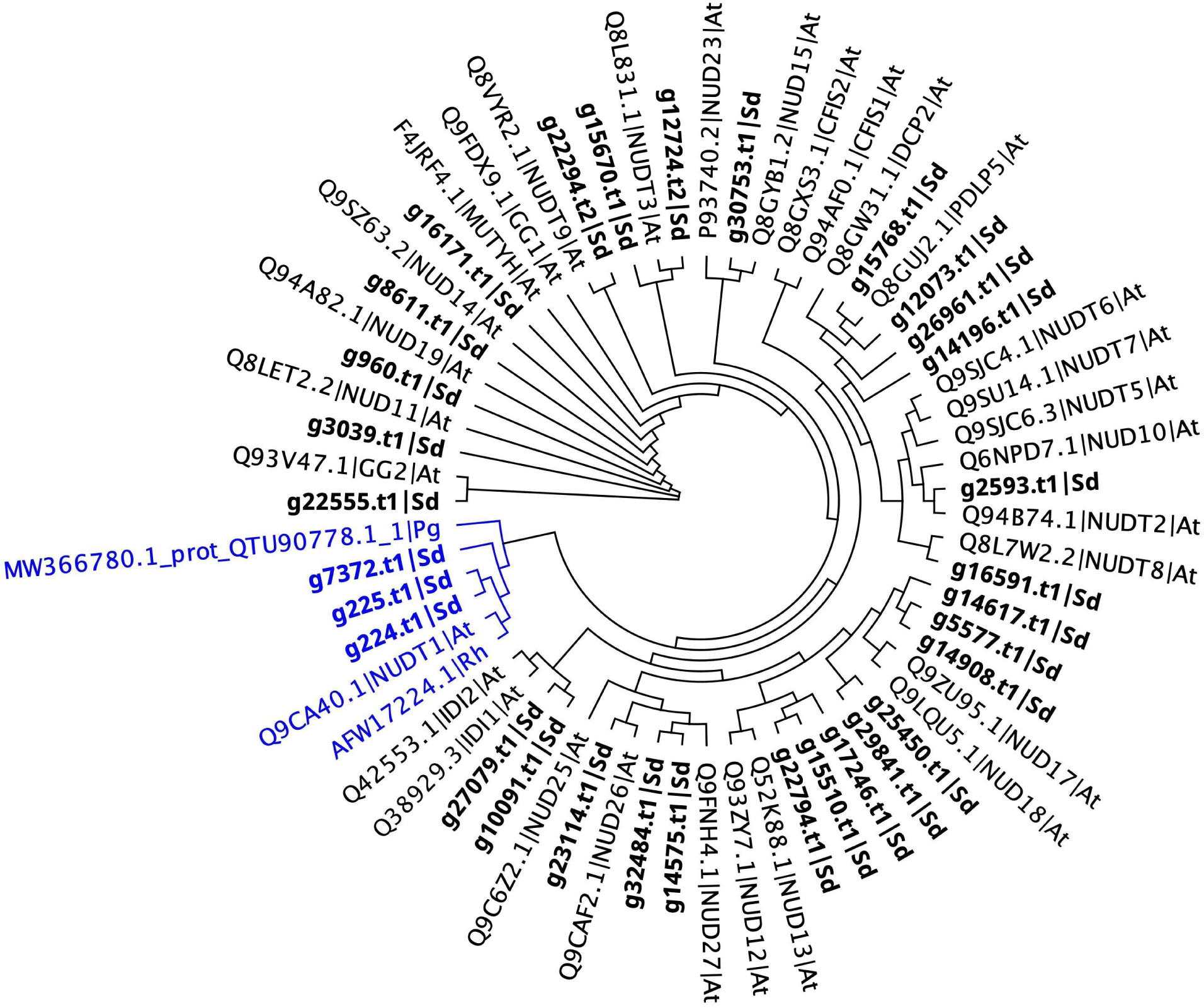
Unrooted neighbor-joining phylogeny of Nudix hydrolases in Arabidopsis thaliana (At) and S. divinorum (Sd; bold). The terpene modifying clade of Nudix hydrolases (Bergmann et al., 2021) is shown in blue and includes sequences from Rosa hybrida (Rh) and Pelargonium graveolens (Pg) in addition to S. divinorum and A. thaliana.

**Supplementary Table 1.** Differential expression of genes implicated in SalA biosynthesis

| Gene<br>name | Annotation<br>numbers | Expression<br>levels by Base<br>Mean | Log<sub>2</sub> FC<sup>a</sup><br>(MeJa<br>induction) | Adjusted *p*-<br>value (MeJa<br>induction) | Log<sub>2</sub> FC in<br>trichome |
| --- | --- | --- | --- | --- | --- |
| *SdCPS2* | g10209 | 55 | 0.59 | 0.76 | 7.71 |
| *SdCS* | g32877 | 2,991 | 2.95 | 0.02 | 7.15 |
| *SdC1H* | g5201 | 1,809 | 1.01 | 0.59 | 8.08 |
| *SdC18H* | g5204 | 1,130 | 2.17 | 0.01 | 7.07 |

<sup>a</sup> FC, fold change

Supplemental Table 1. Differential expression of genes implicated in SalA biosynthesis
| Gene name | Annotation numbers | Expression levels by Base Mean | Log<sub>2</sub> FC<sup>a</sup> (MeJa induction) | Adjusted *p*-value (MeJa induction) | Log<sub>2</sub> FC in trichome |
| --- | --- | --- | --- | --- | --- |
| *SdCPS2* | g10209 | 55 | 0.59 | 0.76 | 7.71 |
| *SdCS* | g32877 | 2,991 | 2.95 | 0.02 | 7.15 |
| *SdC1H* | g5201 | 1,809 | 1.01 | 0.59 | 8.08 |
| *SdC18H* | g5204 | 1,130 | 2.17 | 0.01 | 7.07 |

<sup>a</sup> FC, fold change

## Supplementary Methods file 1

### Trimmomatic

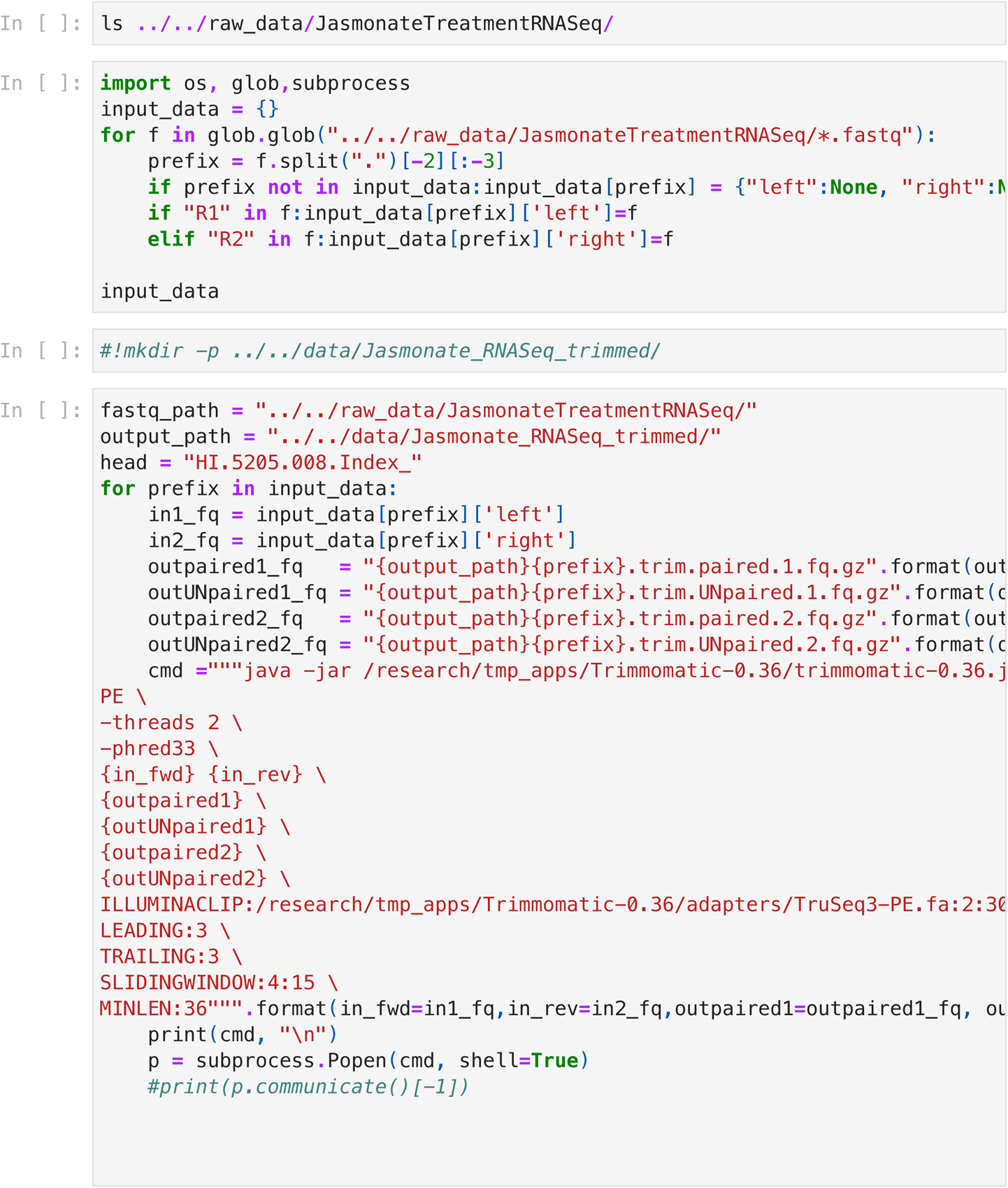

### Organize genome files

Draft genome copied from here

/research/projects/chlamydomonas/lipid_BulkSegregant/Pacbio/analysis/working_assemblie

Draft Annotation copied from here

/research/projects/chlamydomonas/lipid_BulkSegregant/Pacbio/analysis/BRAKER_test/Sd_a

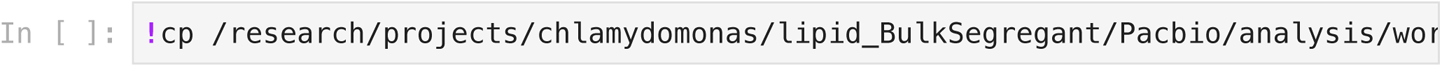

### Build STAR index

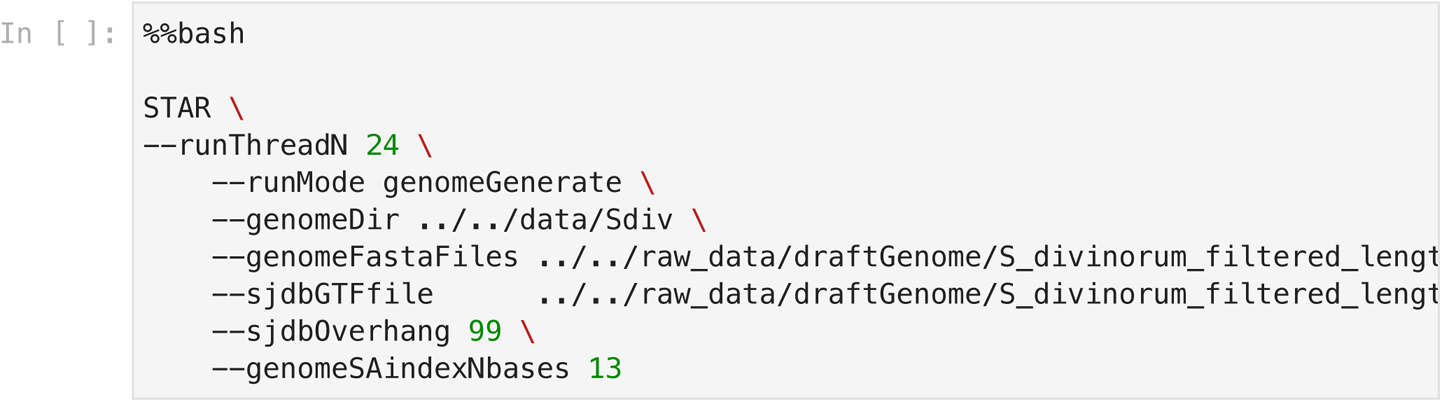

### STAR Alignment

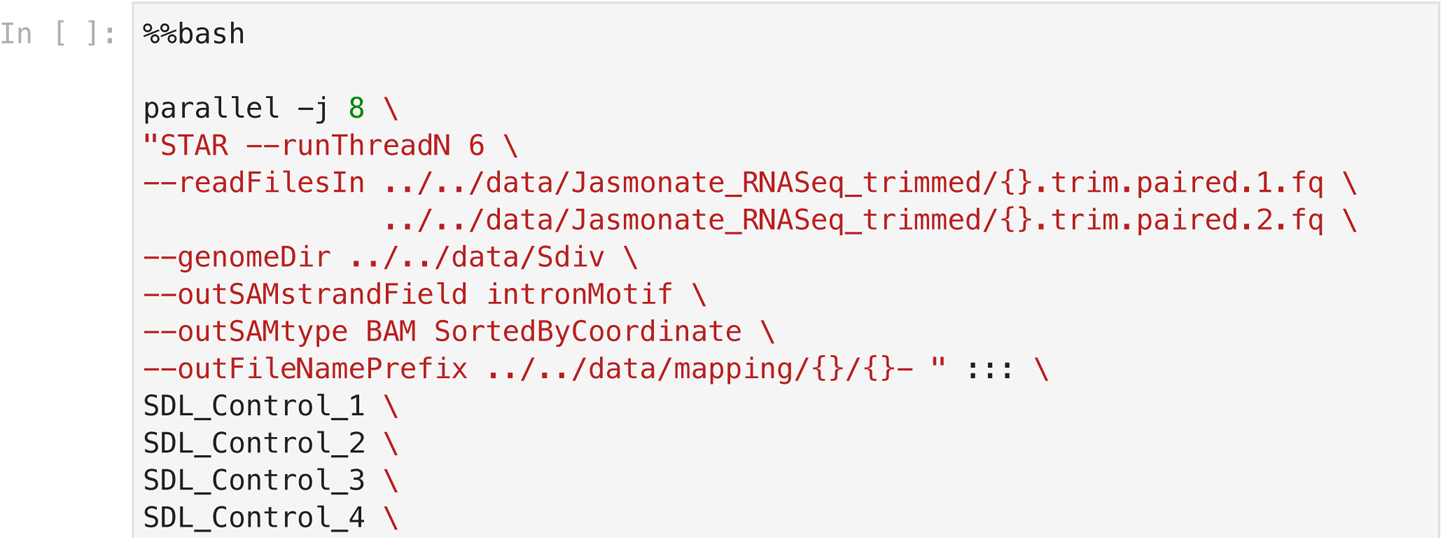

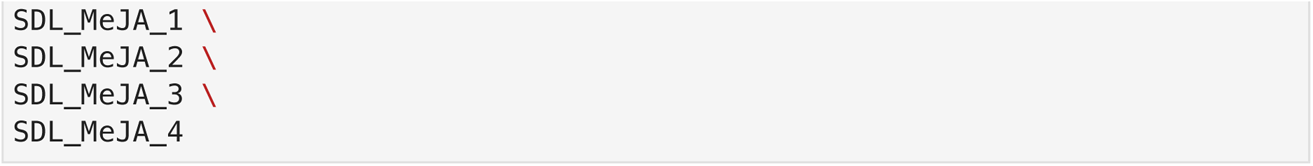

### AddOrReplaceReadGroups

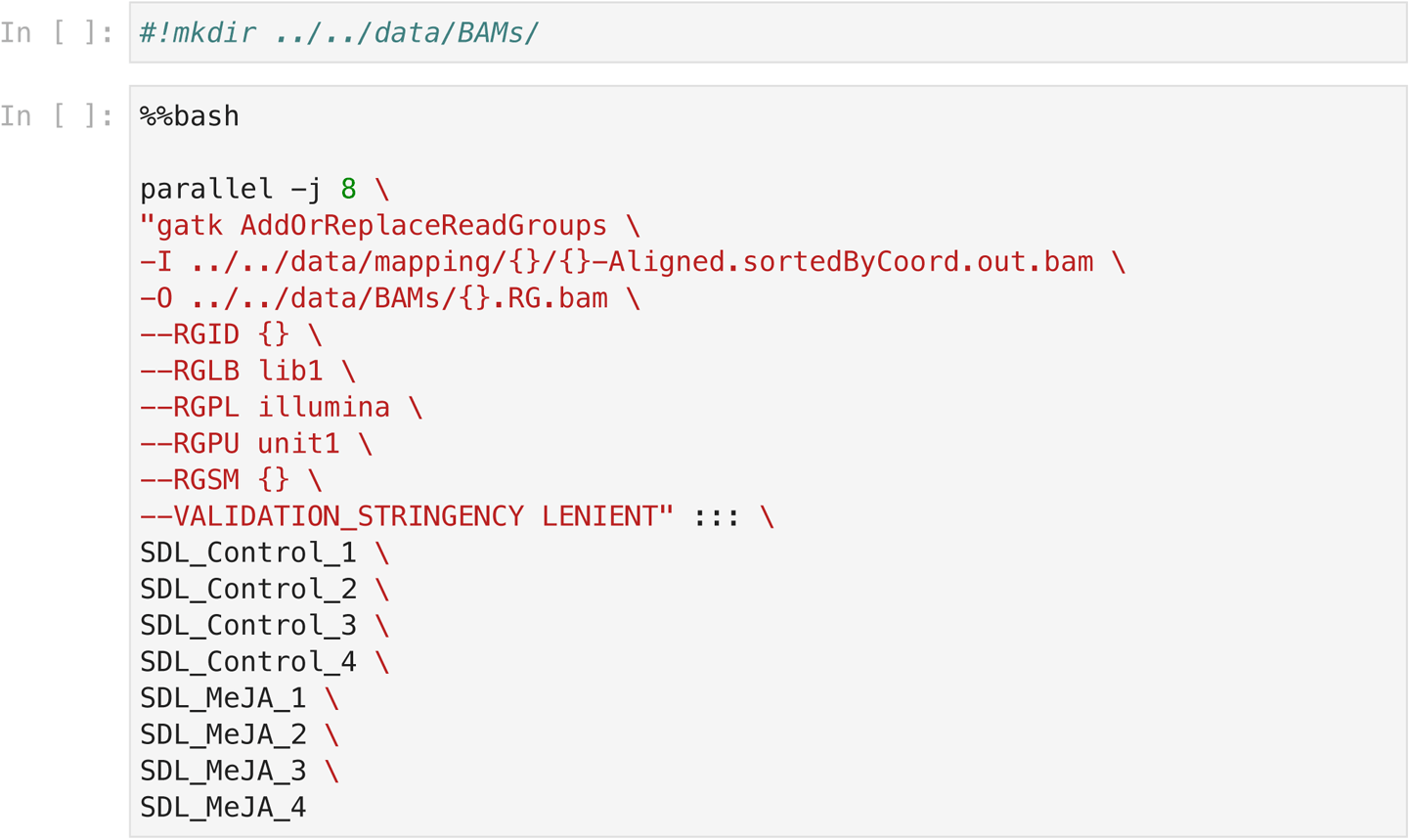

### Index BAMs

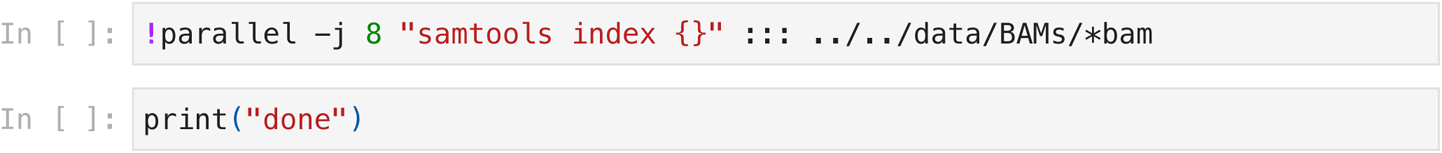

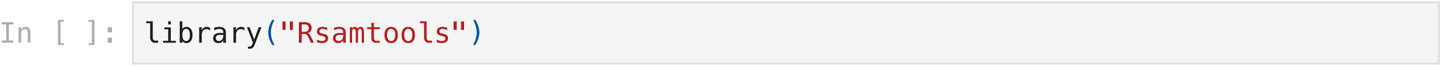

### Assemble information

* include BAMs from Mapping.*.RNASeq.ipynb

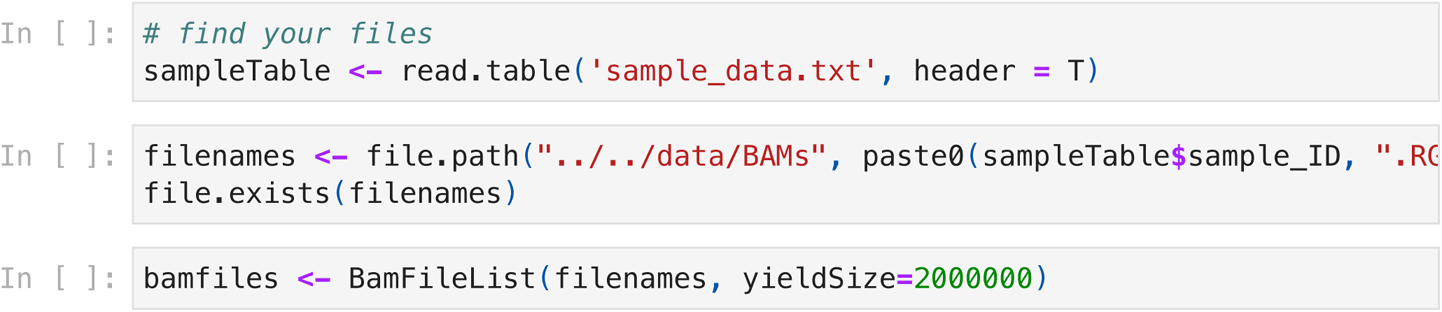

### Annotate genes from de novo gene models

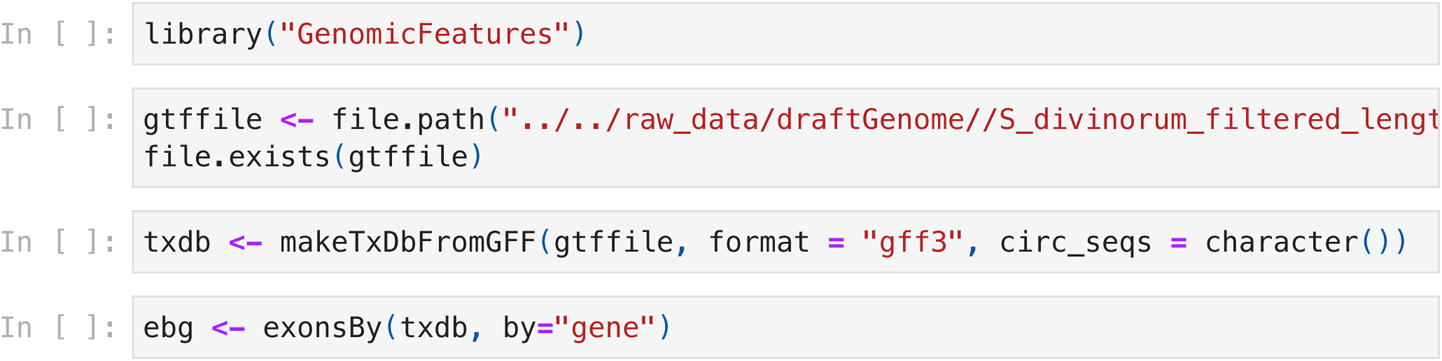

### Summarize read overlaps on genes

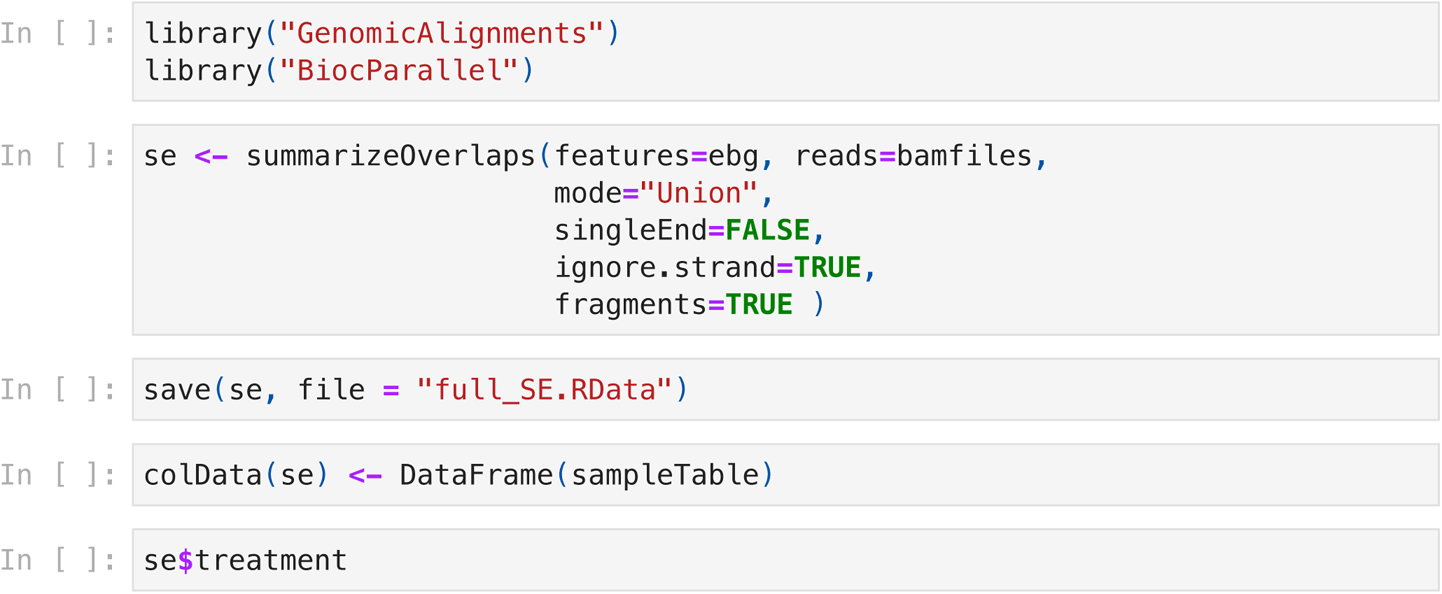

### Analyse differential selection between control and jasmonate treatments

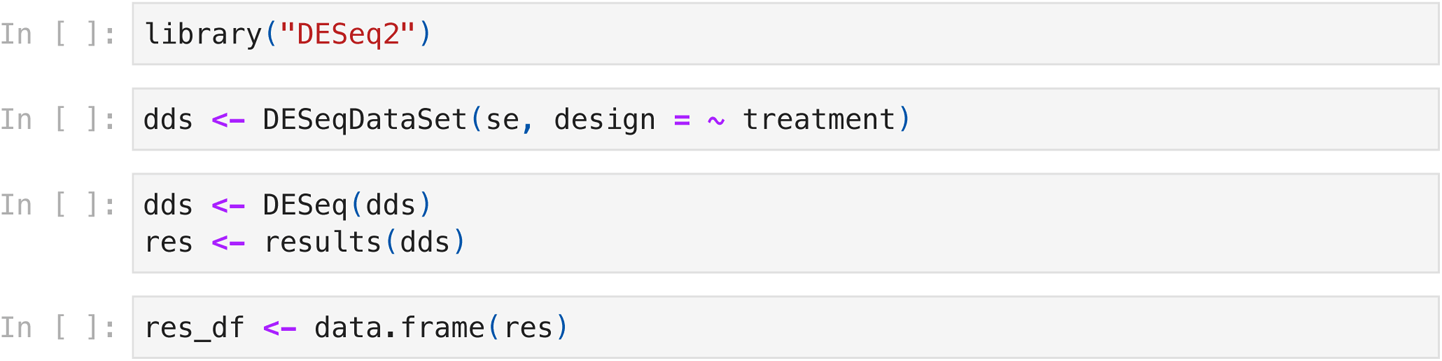

### Write differential expression to file

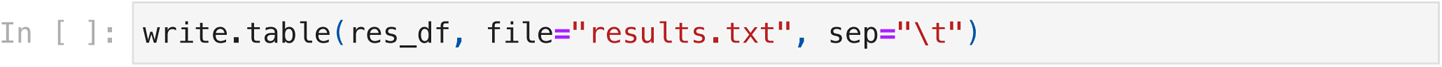

## Mapping Trichome Reads

### Trimmomatic to trim Reads

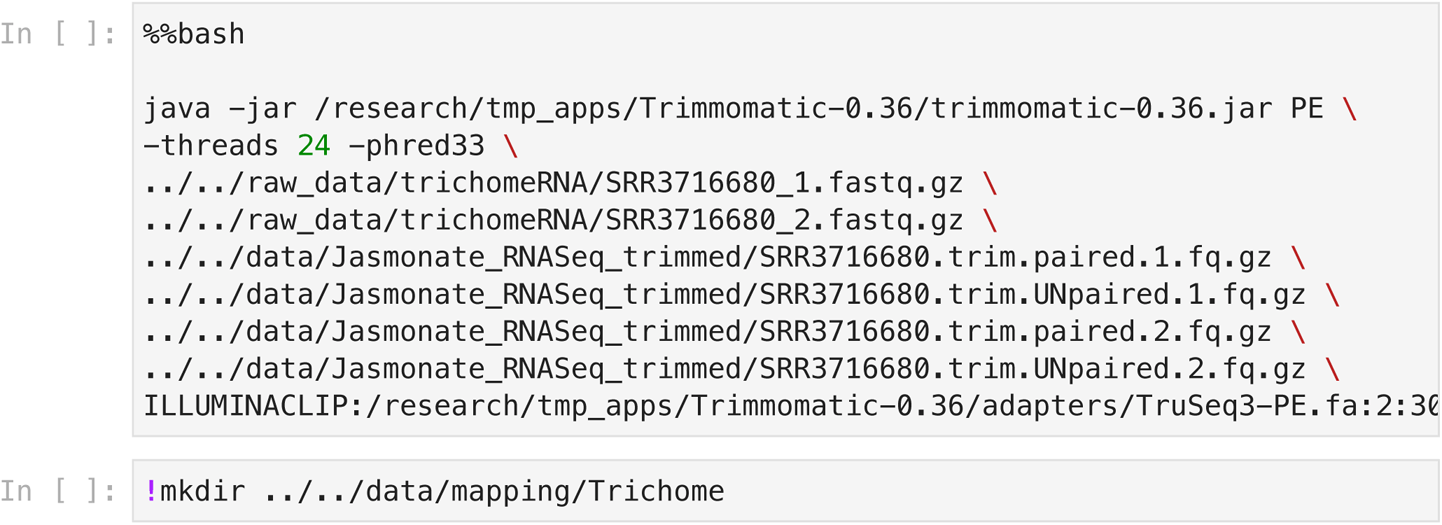

### Unzip Reads

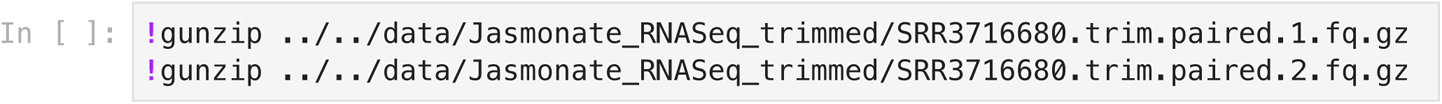

### Align reads to S. divinorum assembly using STAR

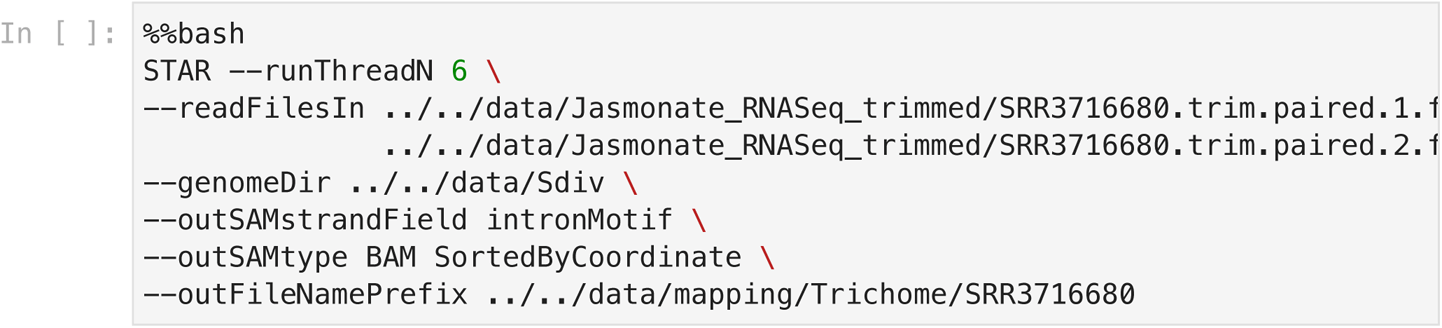

### Add Read Group Labels

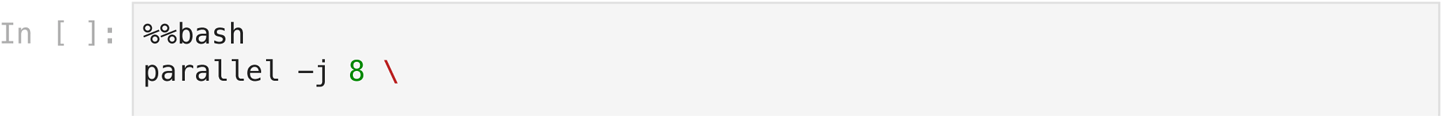

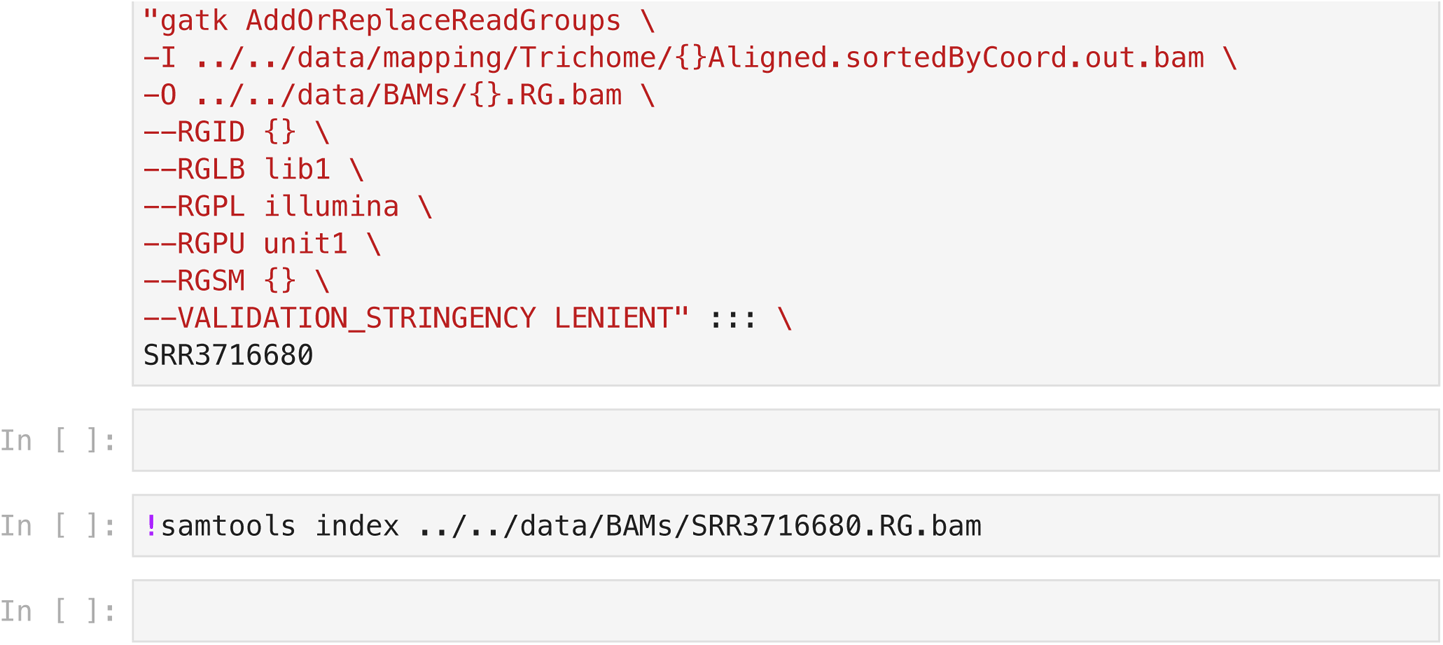

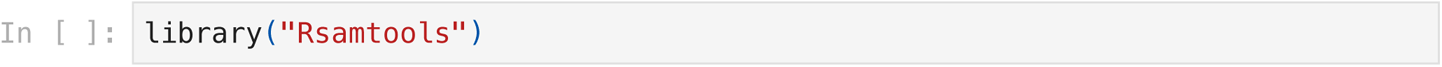

### Assemble information

- include BAMs from Mapping.Trichome.RNASeq.ipynb and Mapping.Jasmonate.RNASeq.ipynb

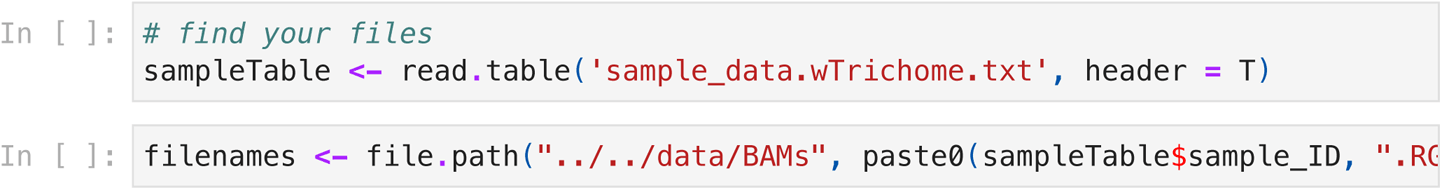

### First step

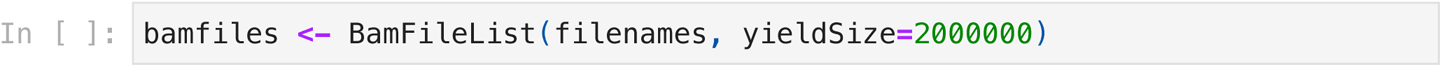

### Annotate genes from de novo gene models

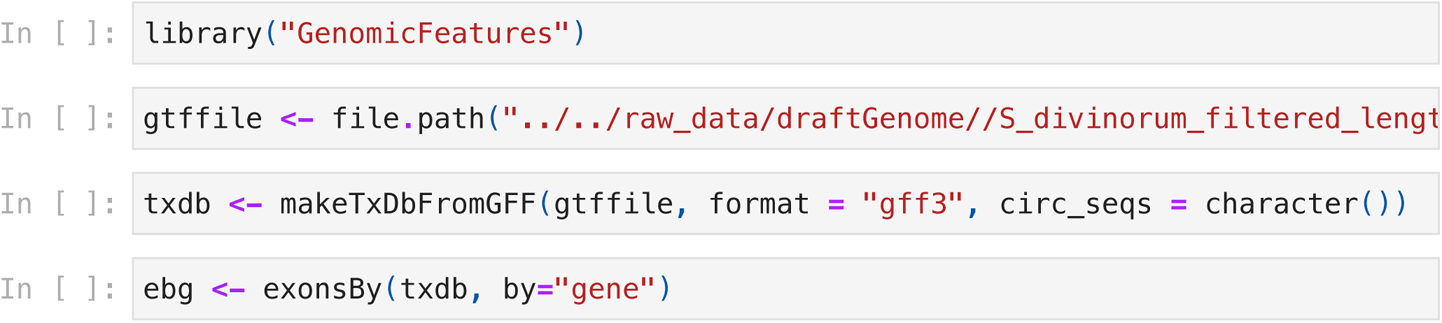

### Summarize read overlaps on genes

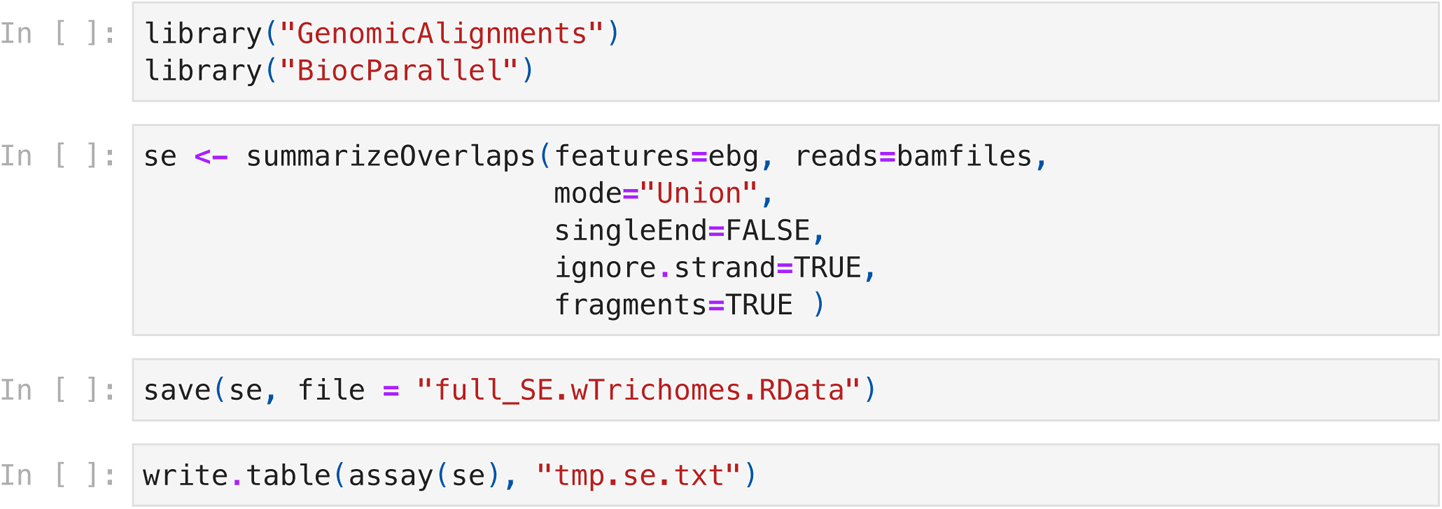

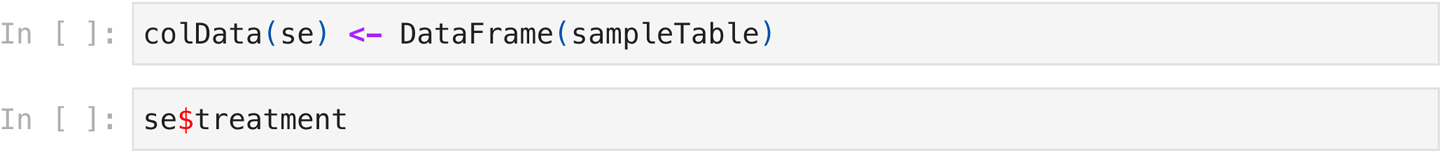

### Analyze fold change without replication following exploratory guidelines in EdgeR

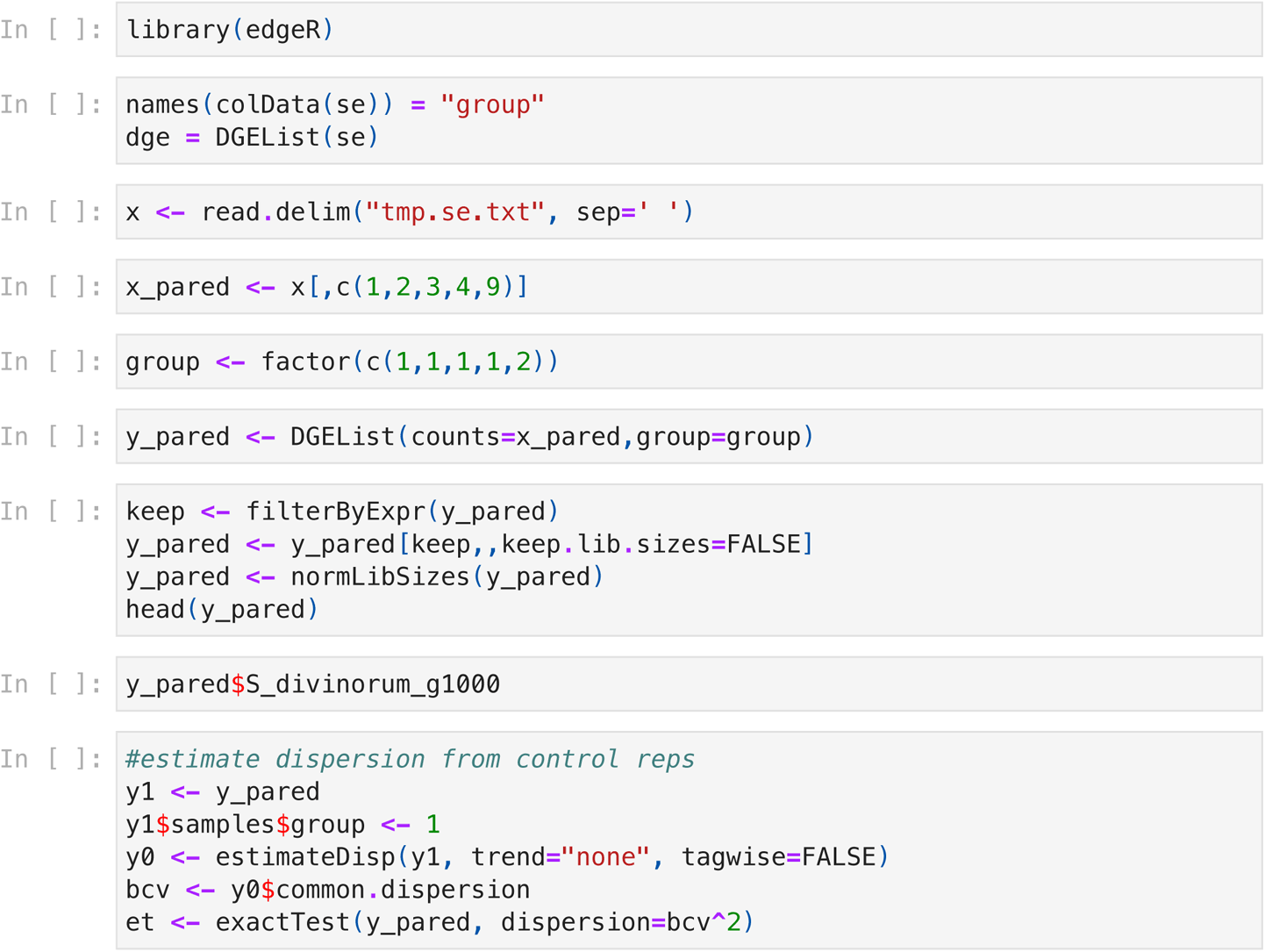

### Write fold change of trichome specific sequence vs leaf

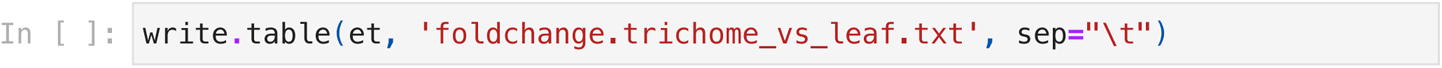

## Supplementary Methods file 2

## A Genome assembly

Text within [ ] represents the names of files that have been simplified for clarity.

### 1. Estimate genome size

Software: Jellyfish v2.2.10

`jellyfish count <(zcat [reads.fasta.gz]) -m 21 -t 10 -s 800M -C jellyfish histo -t 10 --high=100000 mer_counts.jf > reads.histo`

Software: GenomeScope online tool http://genomescope.org/

`Kmer length = 21`

`Read length = 14000`

`Max kmer coverage = 100000`

### 2. Draft genome assembly

Software: hifiasm

`hifiasm -o assembly.asm -t 32 [reads.fastq.gz] 2> assembly.log`

### 3. Purge erroneous allelic duplication and low coverage contigs (https://github.com/dfguan/purge_dups)

i. Map HiFi reads to hifiasm assembly Software: Minimap2 v2.22

`minimap2 -x map-hifi [assembly] [reads.fastq.gz] > [asm.paf.gz]`

ii From alignment, get PB.base.cov, PB.stat files and caculate cutoffs for purge_dups Software: purge_dups v1.2.5

`pbcstat [asm.paf.gz] # produces PB.base.cov and PB.stat files calcuts PB.stat > [cutoffs] 2>calcults.log # calculates cutoffs`

iii Split assembly; perform self-alignment Software: purge_dups v1.2.5

`split_fa [assembly] > [asm.split]`

Software: Minimap2 v2.22

`minimap2 -xasm5 -DP [asm.split] [asm.split] | gzip -c > [asm.split.self.paf.gz]`

Software: purge_dups v1.2.5

`#command below is based on calculated cutoffs, but cutoffs can be adjusted manually. See [calcults.log] to determine if manual adjustment is required. See documentation for tutorial`

`purge_dups -2 -T [cutoffs] -c [PB.base.cov] [asm.split.self.paf.gz] > [dups.bed] 2> [purge_dups.log]`

iv. Get sequences back following after haplotigs are removed Software: purge_dups v1.2.5

`get_seqs -e [dups.bed] [assembly] > [asm.purged.fa] 2> [asm.hap.fa] #haplotig fasta`

### 4. Assemble organelle genomes

Software: ptGAUL

i. CpDNA

`ptGAUL.sh -t 16 -r [relatedCpDNA.fasta] -l [reads.fastq.gz] -o [chloro.out]`

ii. MitoDNA

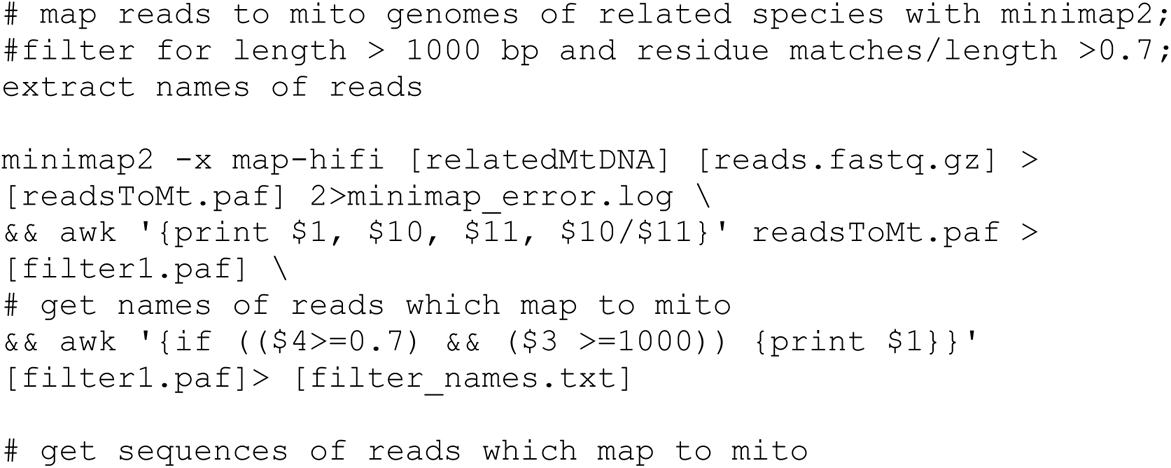

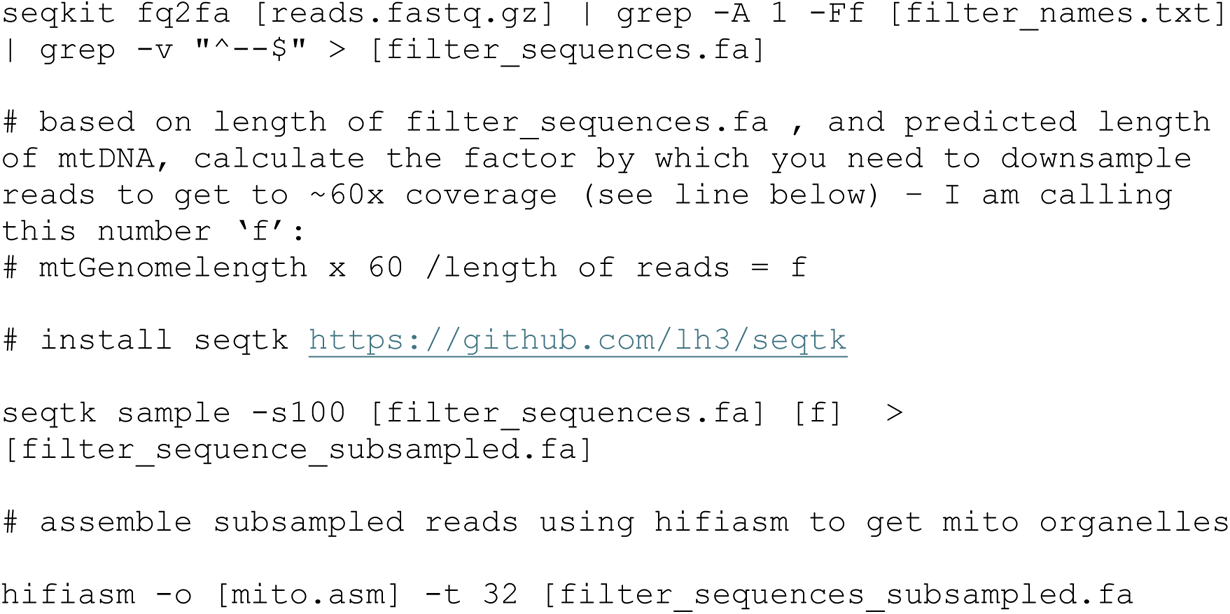

### 5. Filter redundant organellular DNA contigs from final assembly

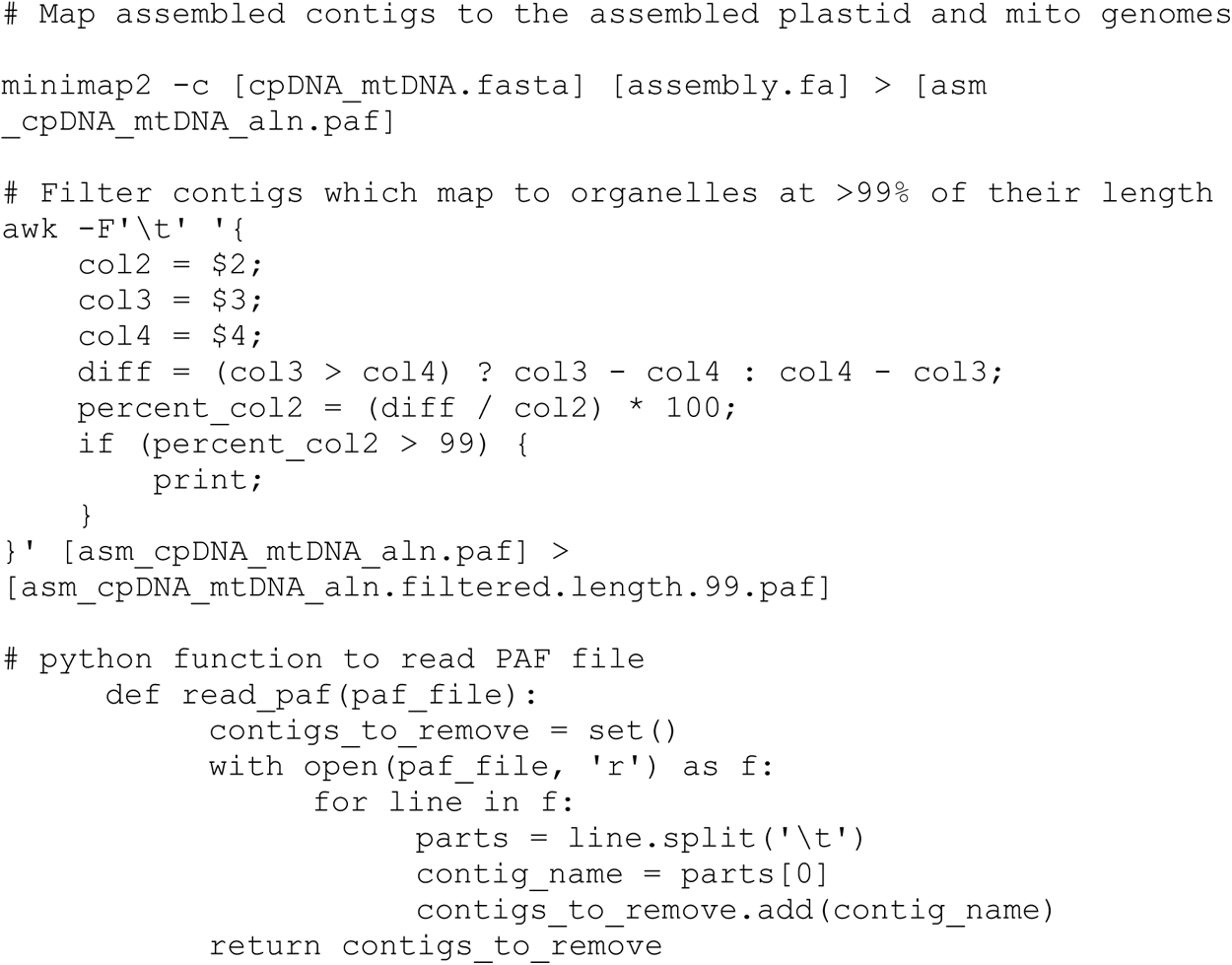

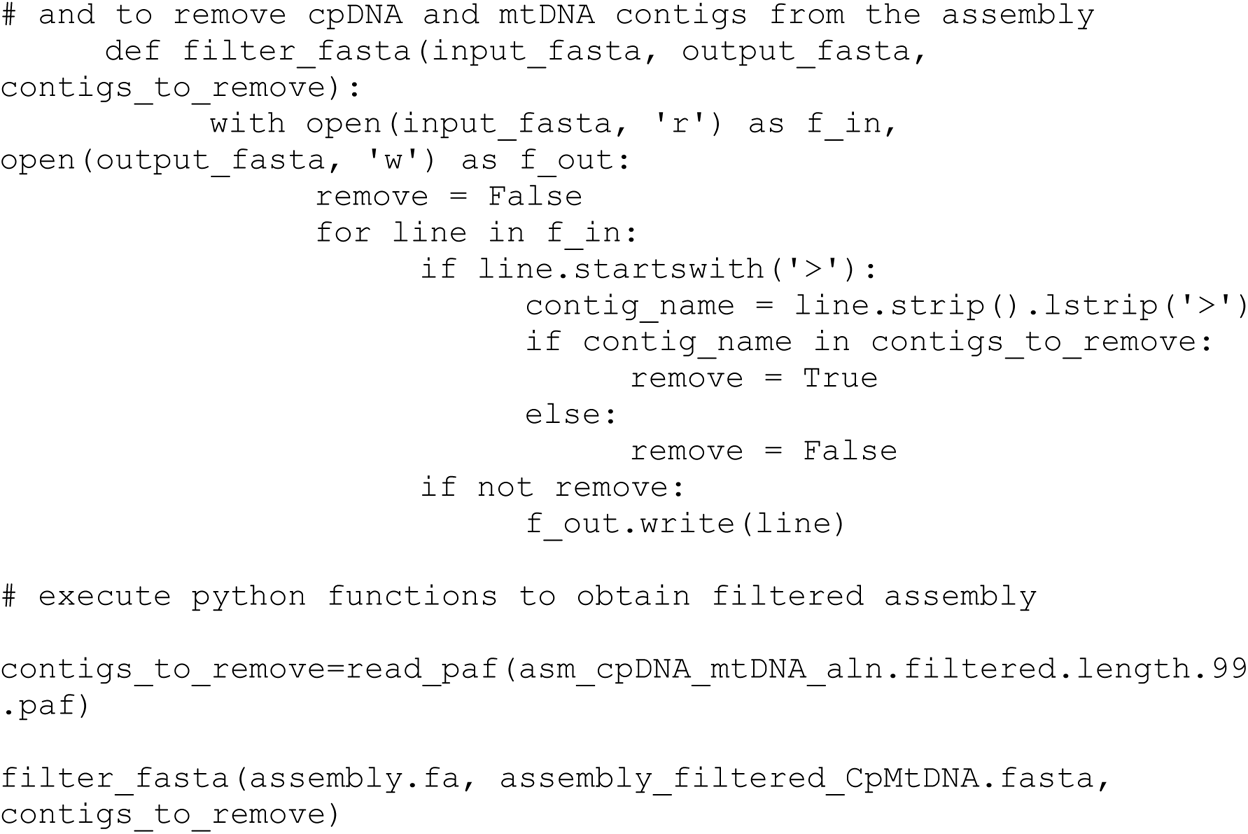

## B Organelle Genome Annotation

Functional annotation of organelle genomes was done using the webtool GeSeq (https://chlorobox.mpimp-golm.mpg.de/geseq.html), using the reference mitochondrial genome of *Arabidopsis thaliana*, and reference plastid genomes of *S. splendens*, *S. miltiorrhiza*, and *S. hispanica* (See Methods section). The following options were used.

For mtDNA annotation:

- BLAT - CDS, tRNA, rRNA
- ARGAGON - default
- tRNAscan – default

For cpDNA annotation:

- BLAT - CDS, tRNA, rRNA
- ARGAGON - default
- tRNAscan – default
- HMMER profile search
- Support annotation by Chloë
- Annotate plastid Inverted Repeat (IR)
- Annotate plastid trans-spliced *rps12*

## C Structural Annotation

### 1. Generate species-specific repeat library, combine with RepBase

Software: RepeatModeler v2.0.5

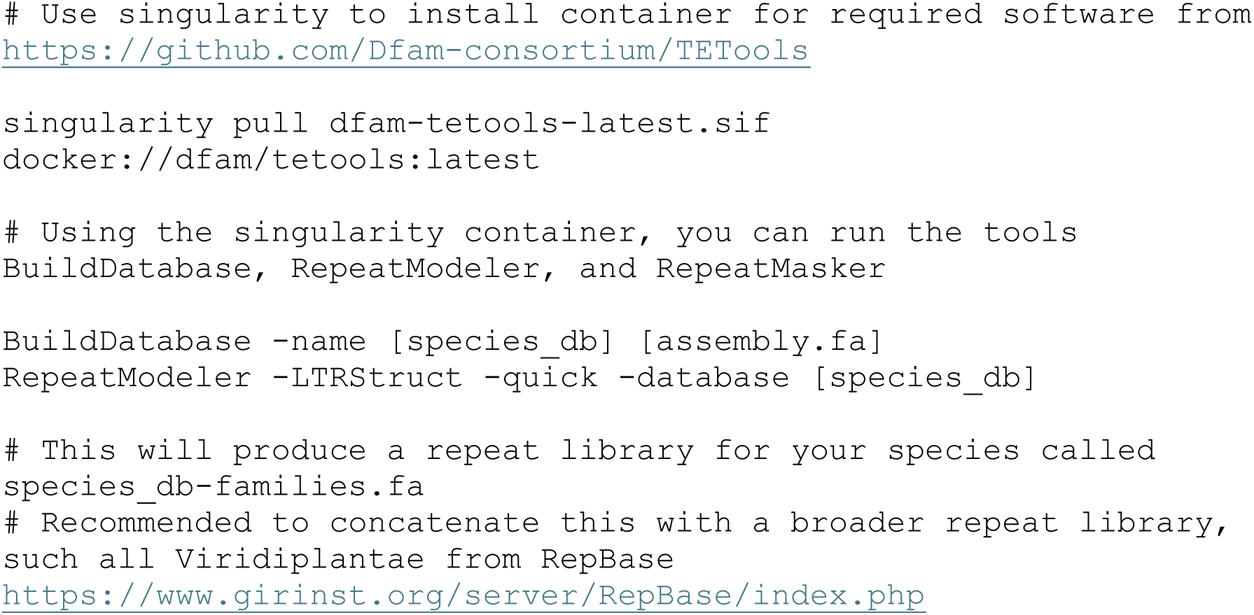

### 2. Mask repeats using repeat library

Software: RepeatMasker v4.1.6

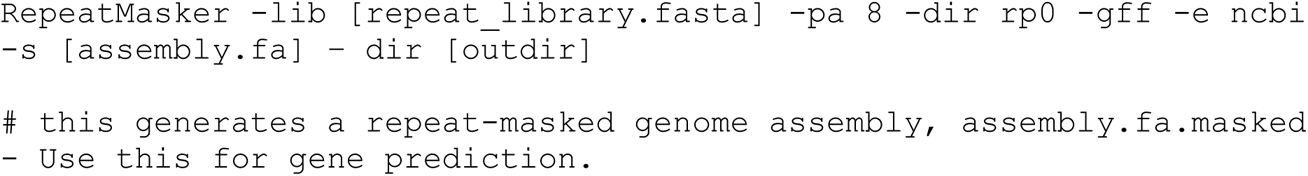

### 3. Predict genes using RNAseq and protein evidence

Software: BRAKER v3.0.8

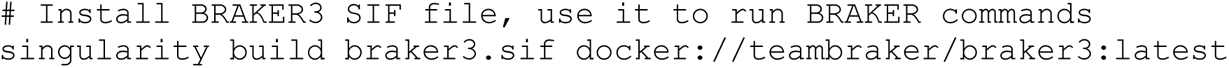

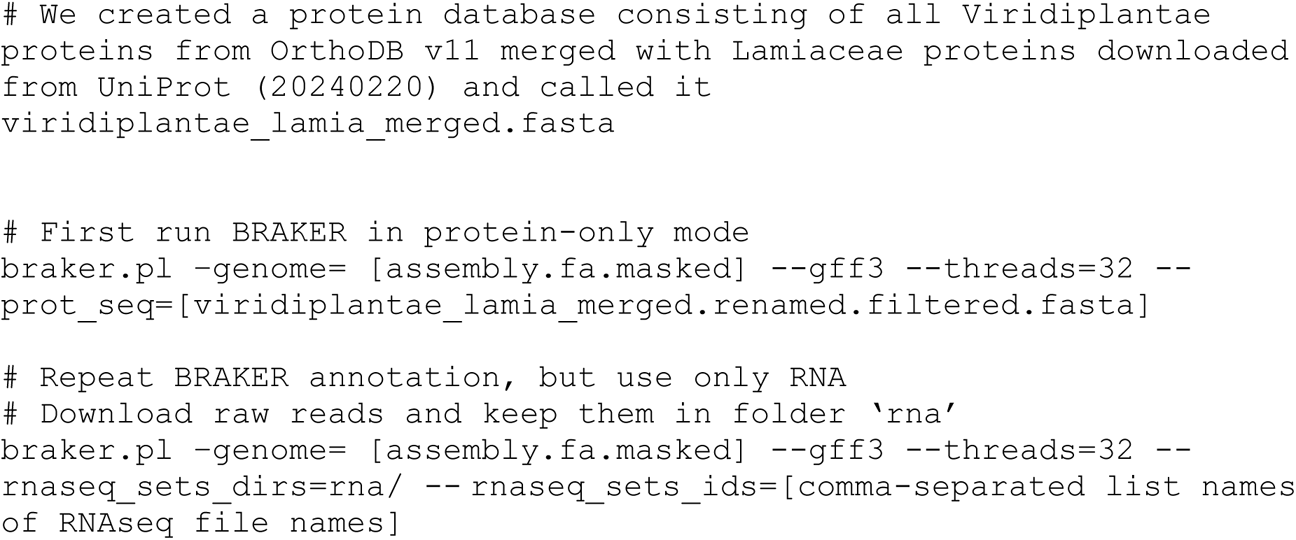

### 4. Predict genes using RNAseq and protein evidence

Software: TSEBRA (access from BRAKER3 SIF file)

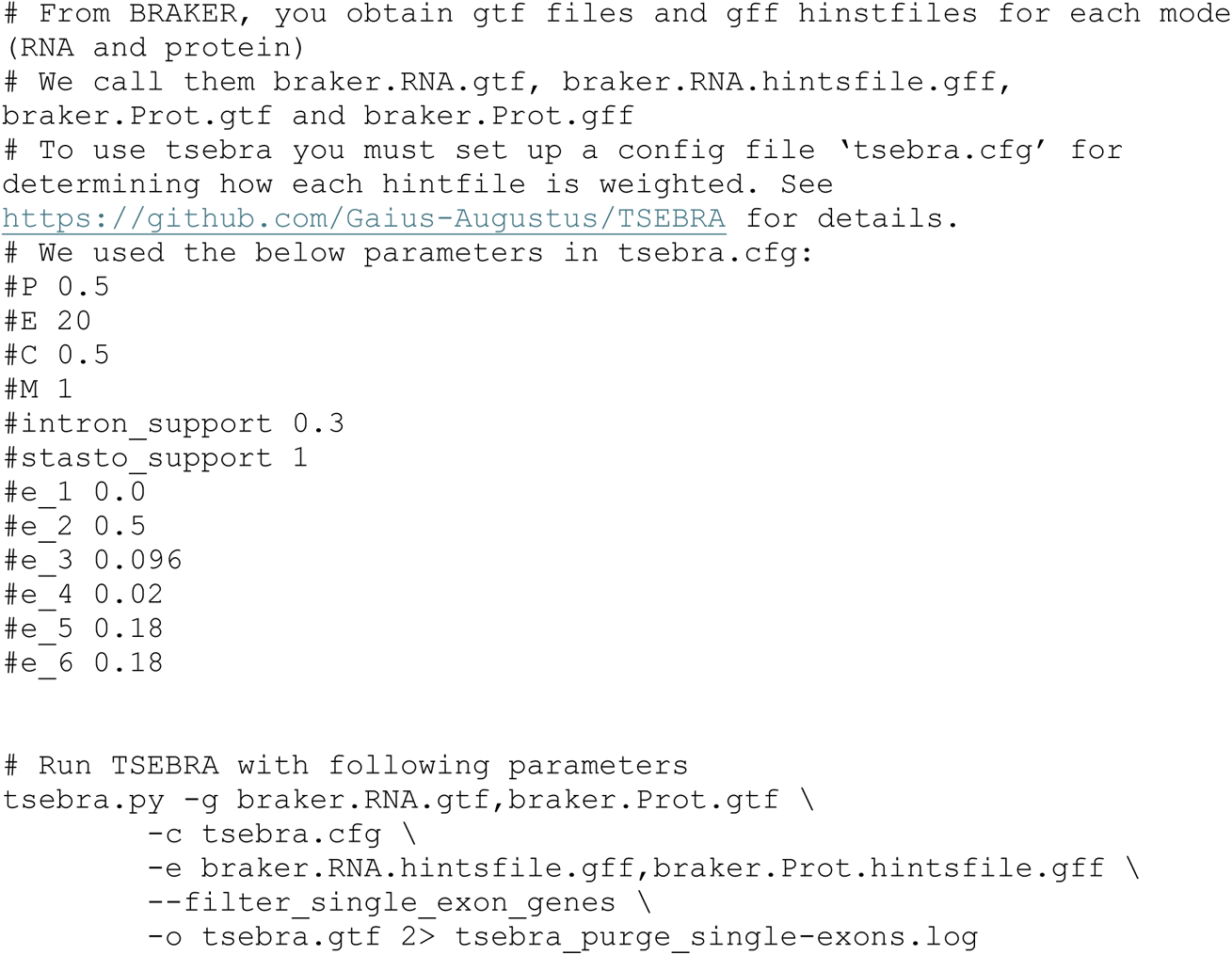

### 5. Format TSEBRA output

The following is based on the pipeline by J. Santangelo (https://github.com/James-S-Santangelo/dcg/tree/main)

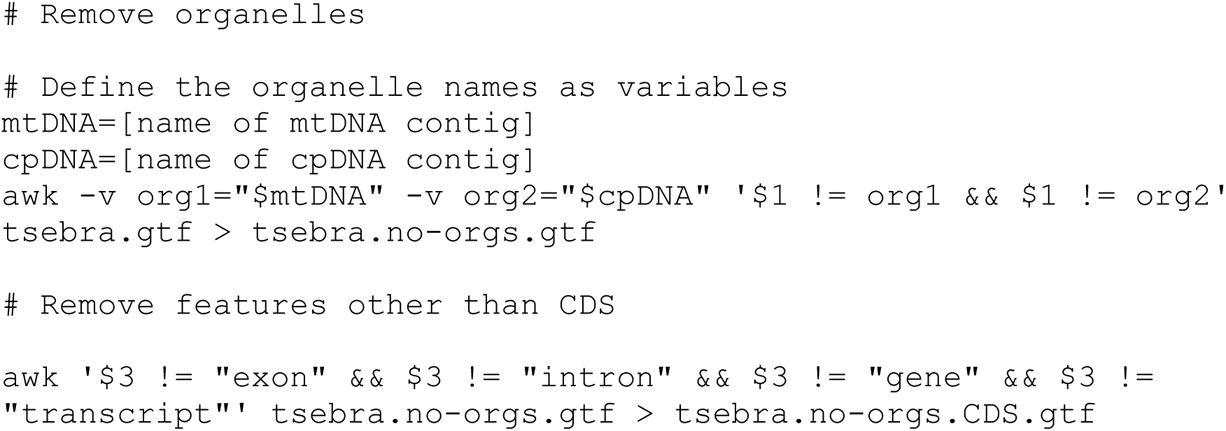

Software: genometools

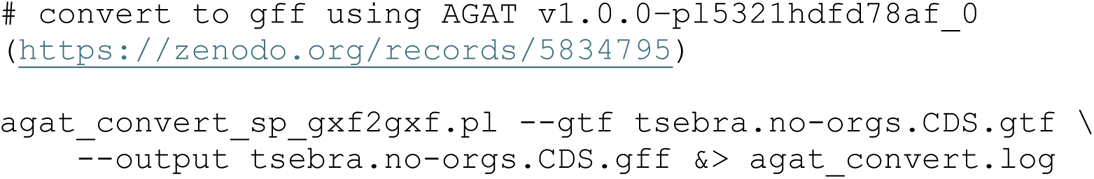

Software: genometools
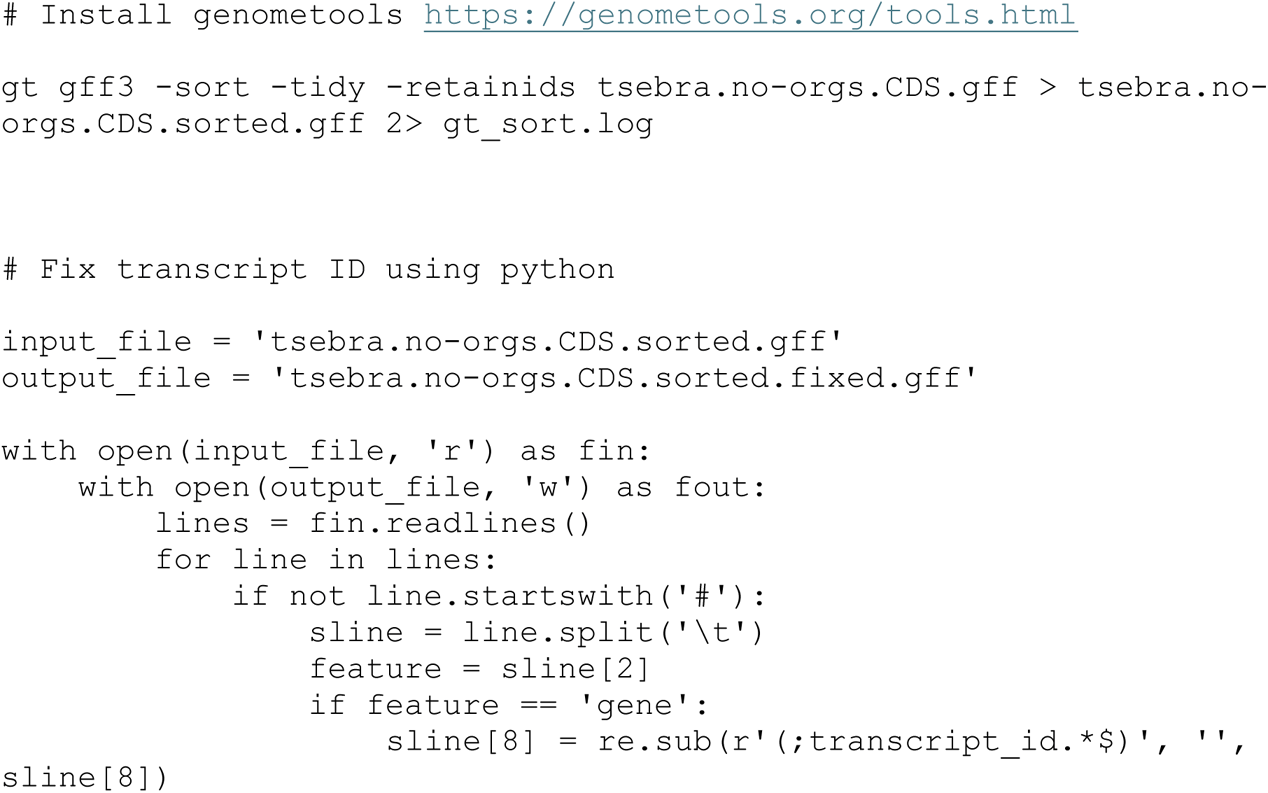

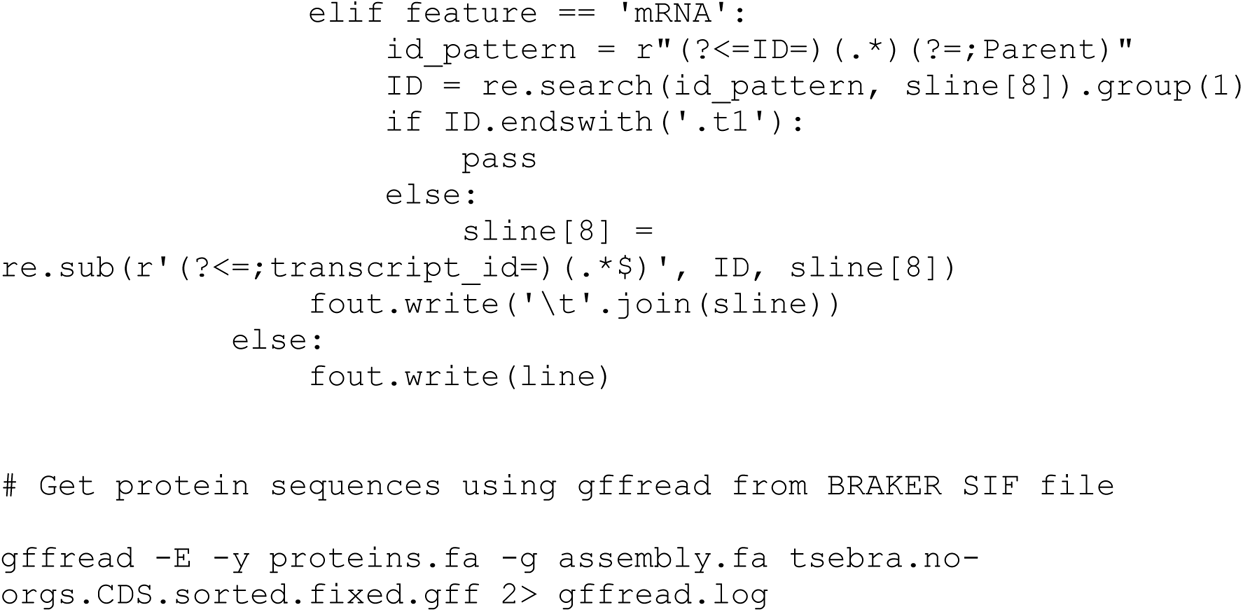

## D Functional Annotation

We combined functional annotations from eggNOG-Mapper, InterProScan and funannotate.

Software: eggNOG-Mapper
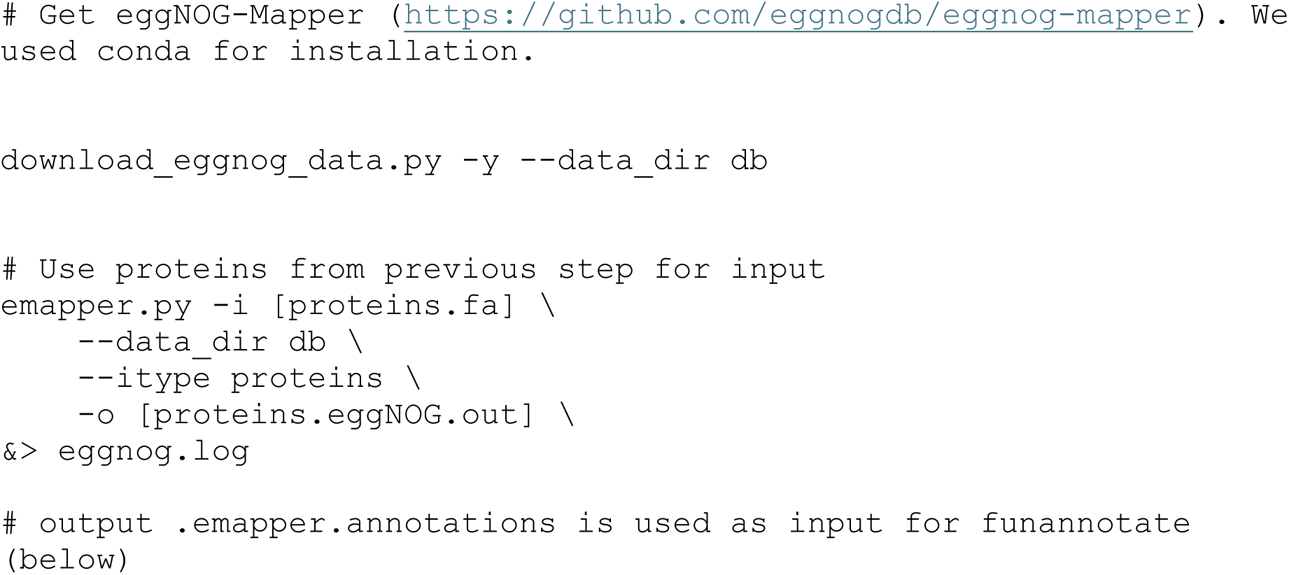

Software: InterProScan
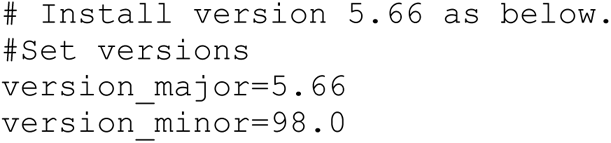

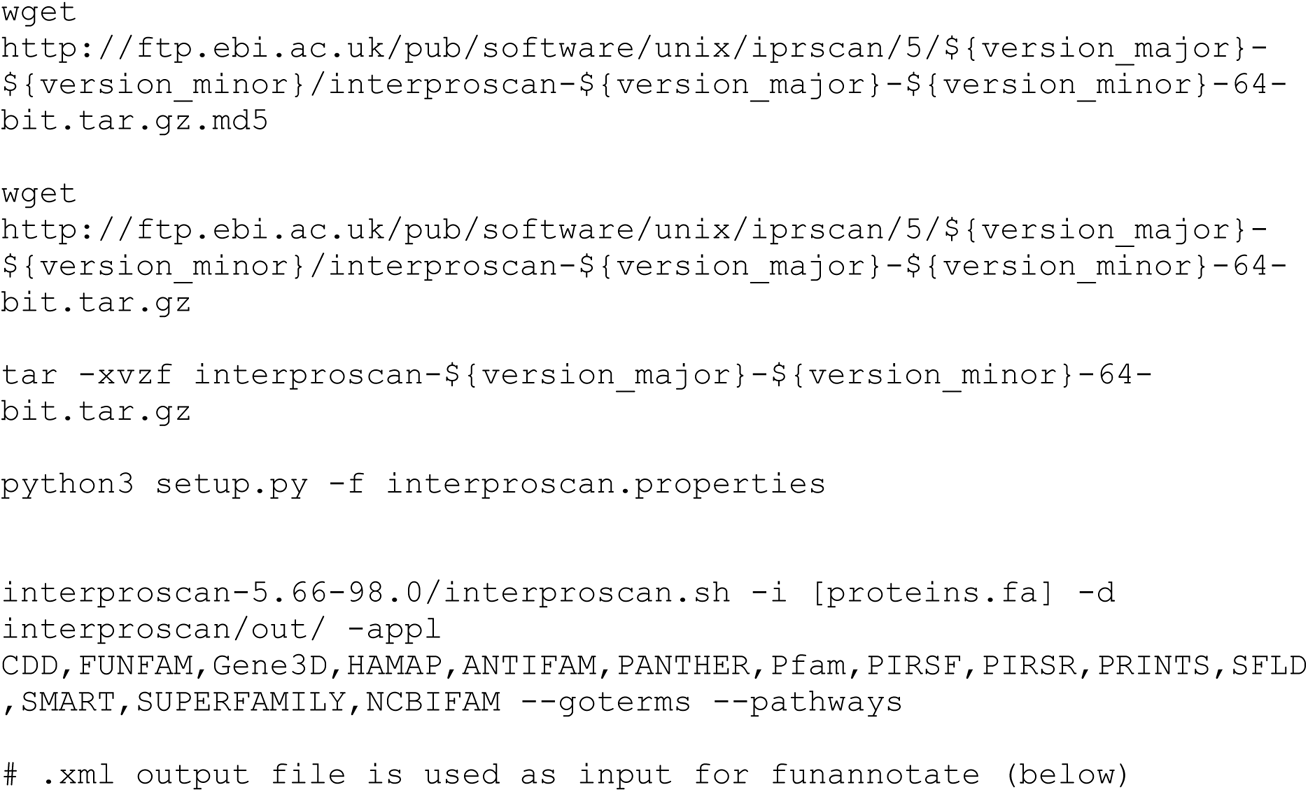

Software: Funannotate
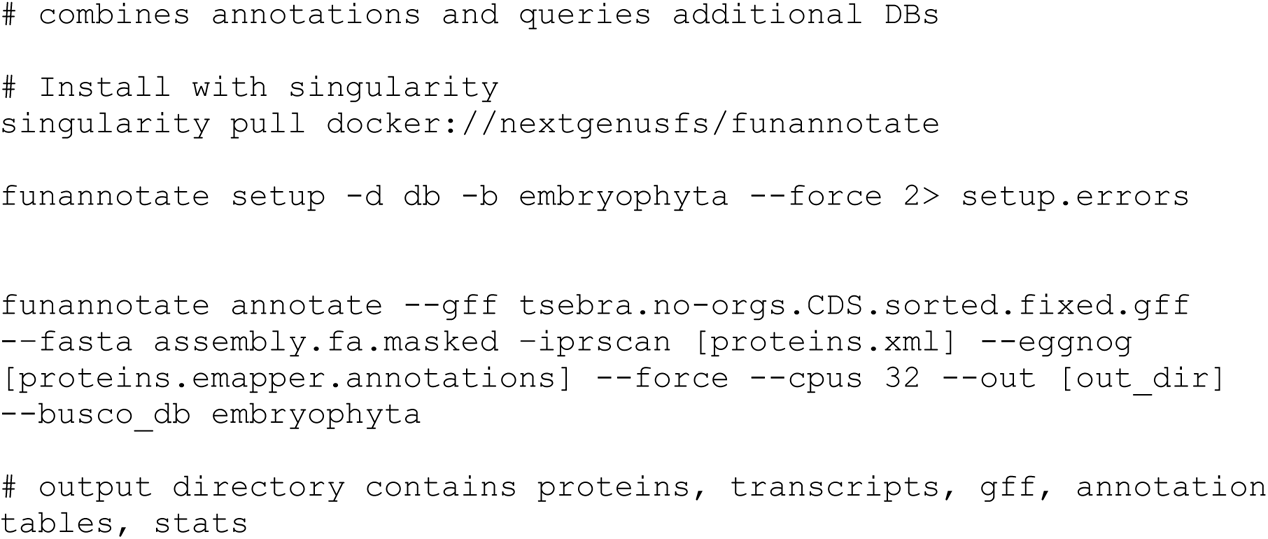

The gff file from funannotate is annotated with pfam and InterPro domains which were used to identify enzymes in classes involved to SalA synthesis (See Methods; Supplementary Data Set 1L for details). As described in the Methods, protein-protein BLAST searches against characterized proteins from closely related species were used to identify diterpene synthases. CYPs were putatively classified into clans and subfamilies similarly using a protein-protein BLAST search of predicted *S. divinorum* (based on pfam and InterPro domains) CYPs against a database of 9739 plant CYPs previously classified by Dr. David Nelson (https://drnelson.uthsc.edu/plants) (See Extended Data Set 2).

Based on the pipeline provided by J. Santangelo (https://github.com/James-S-Santangelo/dcg), if a protein was annotated as ‘hypothetical protein’, but was assigned a fully resolved enzyme commission (EC) number, the annotation was replaced with the EC number’s product in the ExPASSY Enzyme database.

## E Identification of diterpene biosynthetic gene clusters (BGCs)

The PlantiSMASH (https://github.com/plantismash/plantismash) webserver was used with default settings to detect signatures of BGCs. Terpene synthesis clusters were further classified as diterpene clusters based on overlap with predicted diterpene synthase genes (See above).

## F Genome evolution

### 1. Pairwise synteny analysis with GENESPACE

The protein fasta file and gff file for *S. splendens* was downloaded from NCBI (Supplementary Data Set 1L for accession number). It needs to be converted to a bed file for GENESPACE analysis.
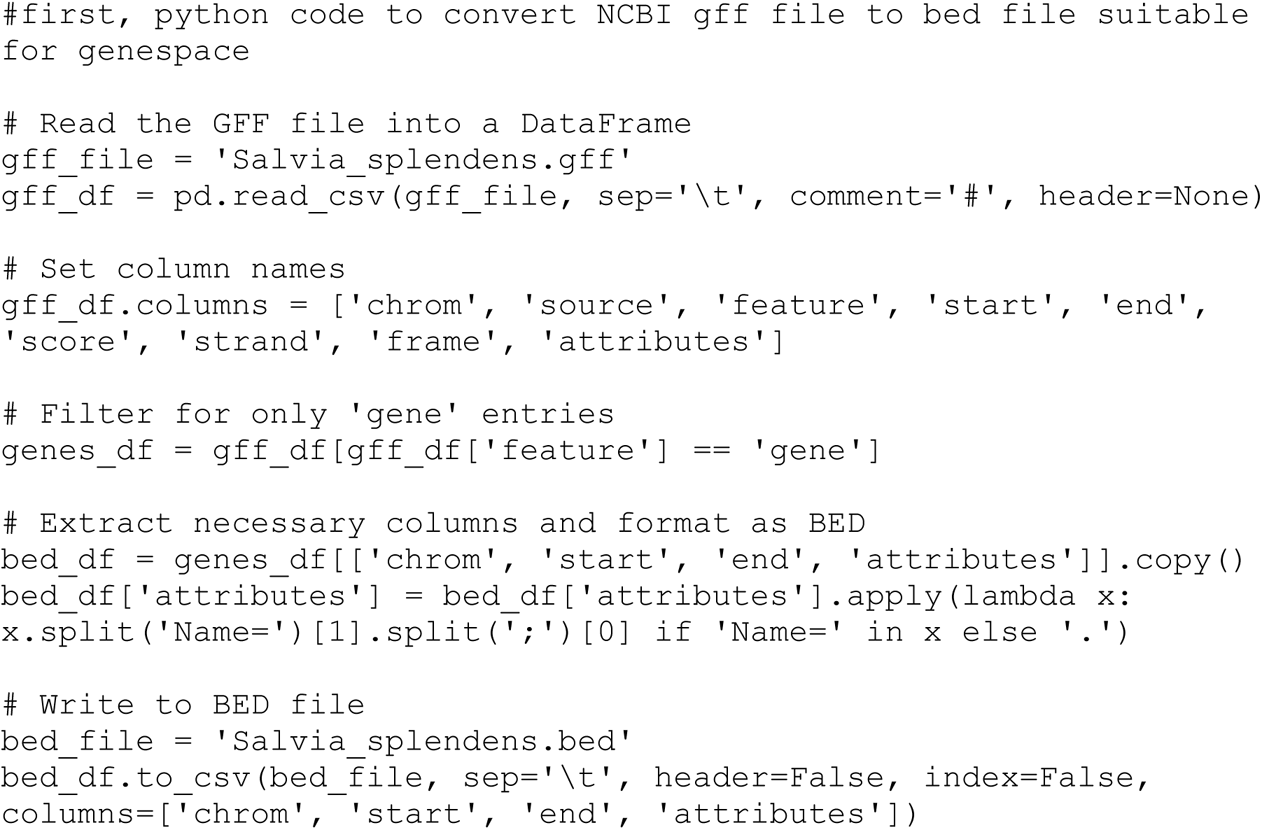

The protein fasta file header must exactly match the bed file. The following python code accomplishes this, ensuring only the longest isoform is retained.
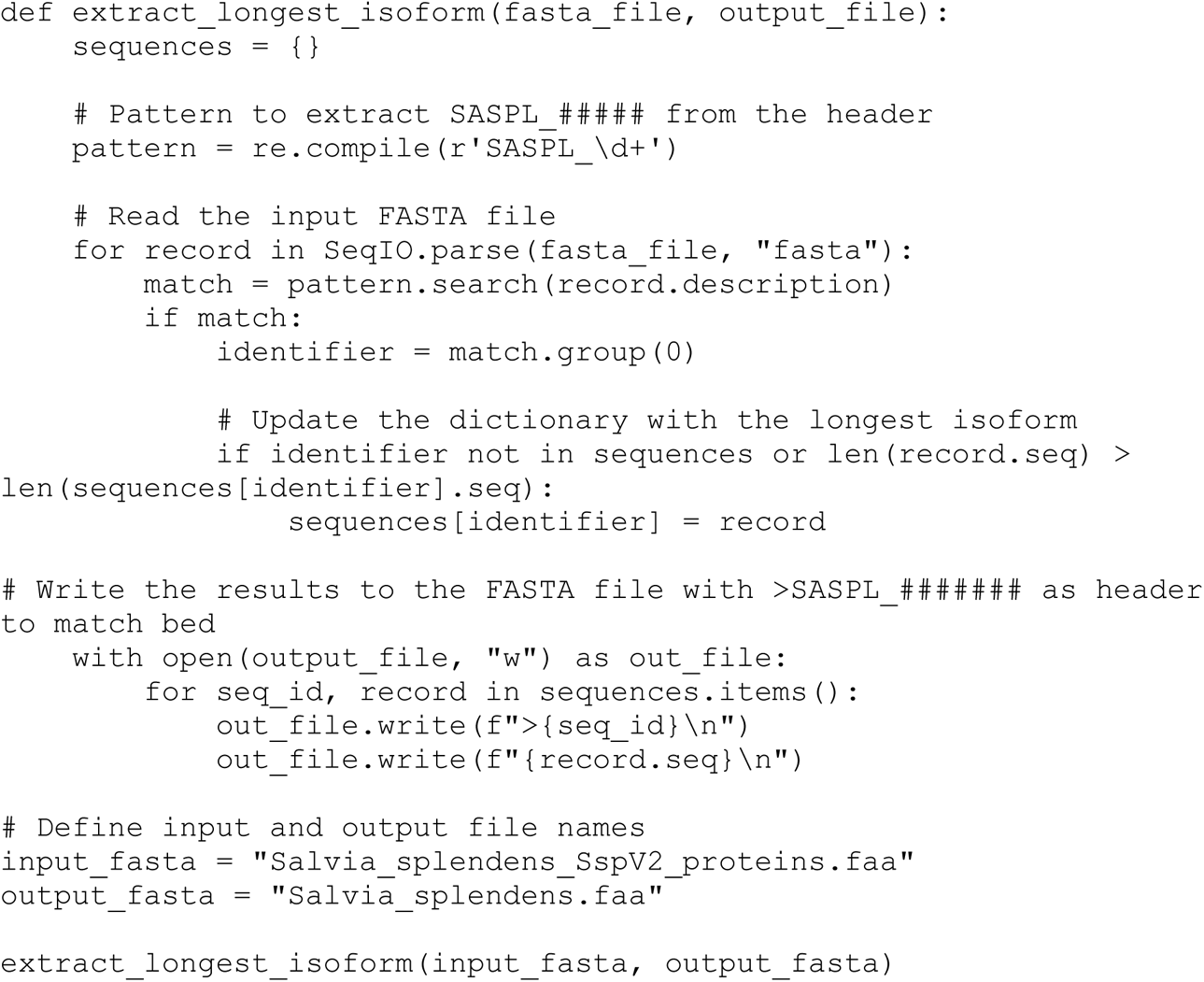

*S. divinorum* bed and corresponding protein fasta filtes are similarly obtained, starting with output from functional annotation. The ID column from the bed file exactly matches the fasta header. The files are called `S_divinorum.bed` and `S_divinorum.faa.`

To set up genespace directory, create a folder called bed and add `S_divinorum.bed` and `S_splendens.bed.` Create another folder called peptide and add `S_divinorum.faa` and `S_splendens.faa.`

Software: GENESPACE
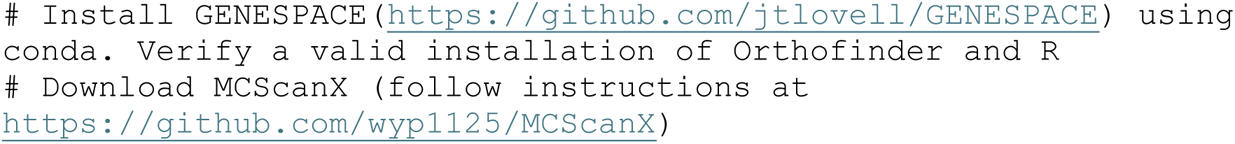

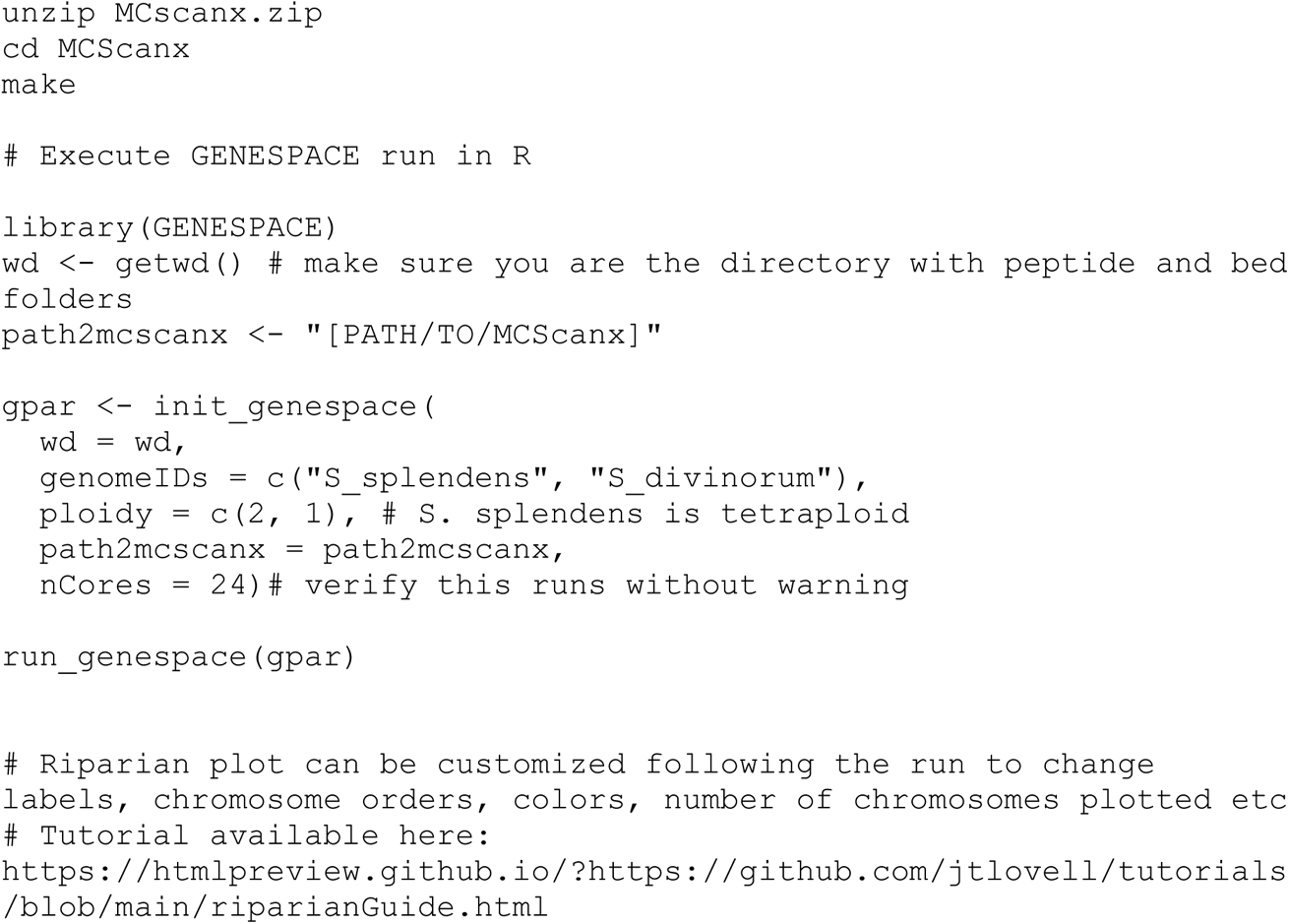

### 2. Find orthogroups and orthologs

Protein sequences (.faa) for *S. hispanica*, *S. bowleyana*, *S. miltiorrhiza*, *S. rosmarinus* and *Sesamum indicum* were downloaded from NCBI (Supplementary Data Set 1L for accession numbers). Fasta files were filtered to retain only the longest isoforms of each protein. Due to differences in the formatting of fasta headers and corresponding gff files, proteins of each species were filtered separately using in-house python scripts. Along with the longest protein isoforms of *S. divinorum* and *S. splendens* (above), they were placed in a folder we are calling `comparative_genomes` for simplicity. Ensure that each fasta file has a species-specific identifier in the header, as this will be important for downstream analyses of gene families.

Software: OrthoFinder
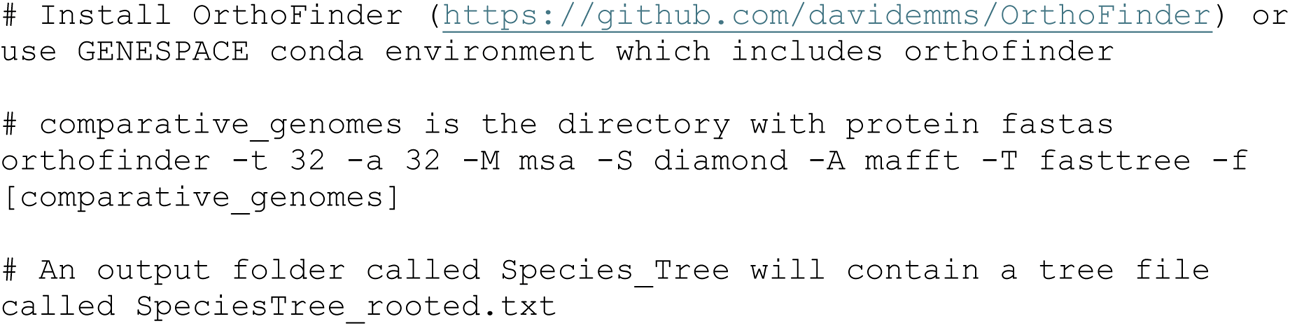

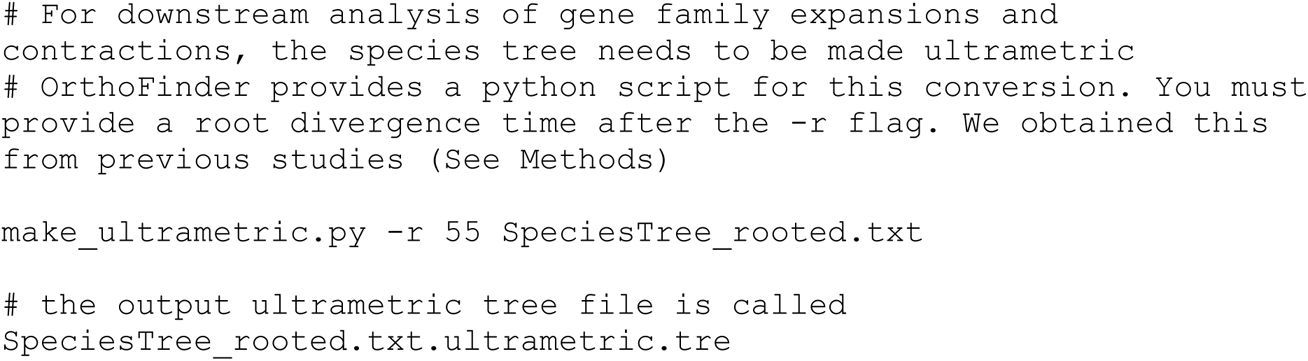

### 3. Gene family expansion and contraction

Gene family expansion and contraction analysis was done using CAFE5 (https://github.com/hahnlab/CAFE5), based on the ultrametric tree file `SpeciesTree_rooted.txt.ultrametric.tre` and orthogroup gene counts obtained from OrthoFiner (above). Install CAFE5 according to instructions provided (https://github.com/hahnlab/CAFE5).

Following OrthoFinder run, an Orthgroup gene count file `Orthogroups.GeneCount.tsv` will be in a directory called `Orthogroups`. This file must be slightly reformatted for use with CAFE5. Code for prepping `Orthogroups.GeneCount.tsv` is provided below based on a previously described pipeline (https://github.com/elsemikk/Willisornis_Genome_Assembly/blob/master/4.2_Gene_Family_Ex <u>pansions.md</u>). A detailed tutorial for CAFE5 is also available https://iu.app.box.com/v/cafetutorial-files.

Software: CAFE5
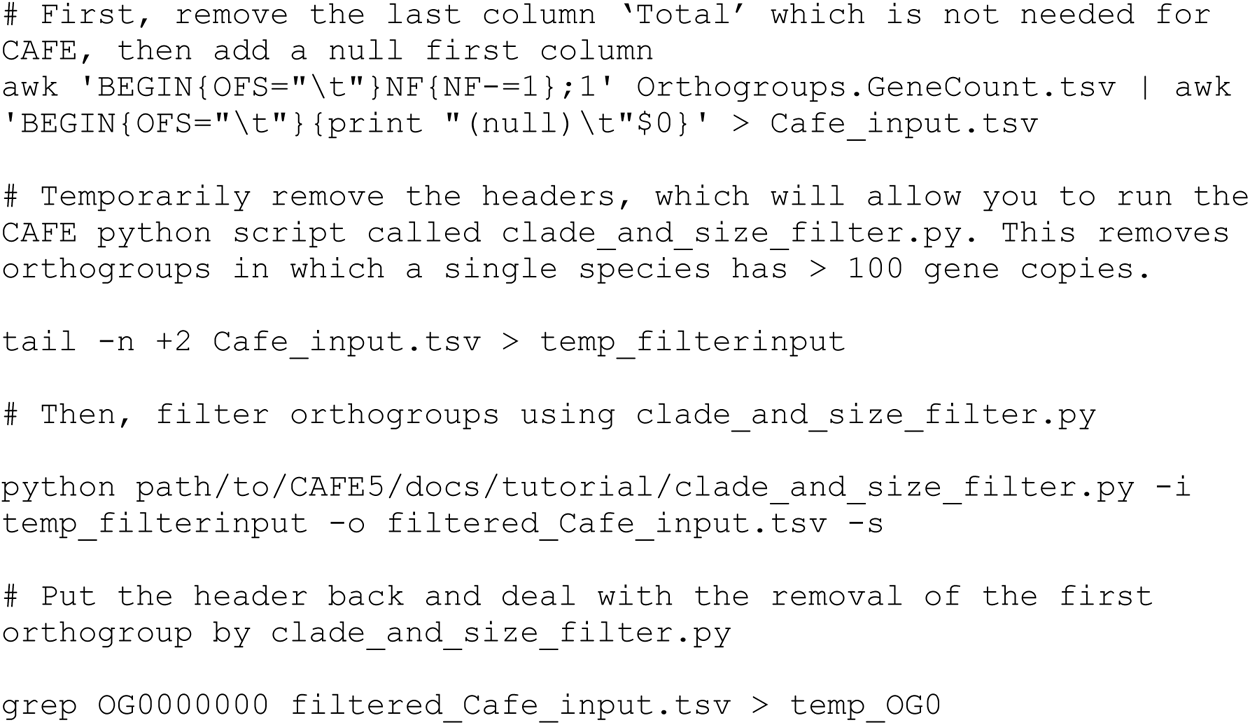

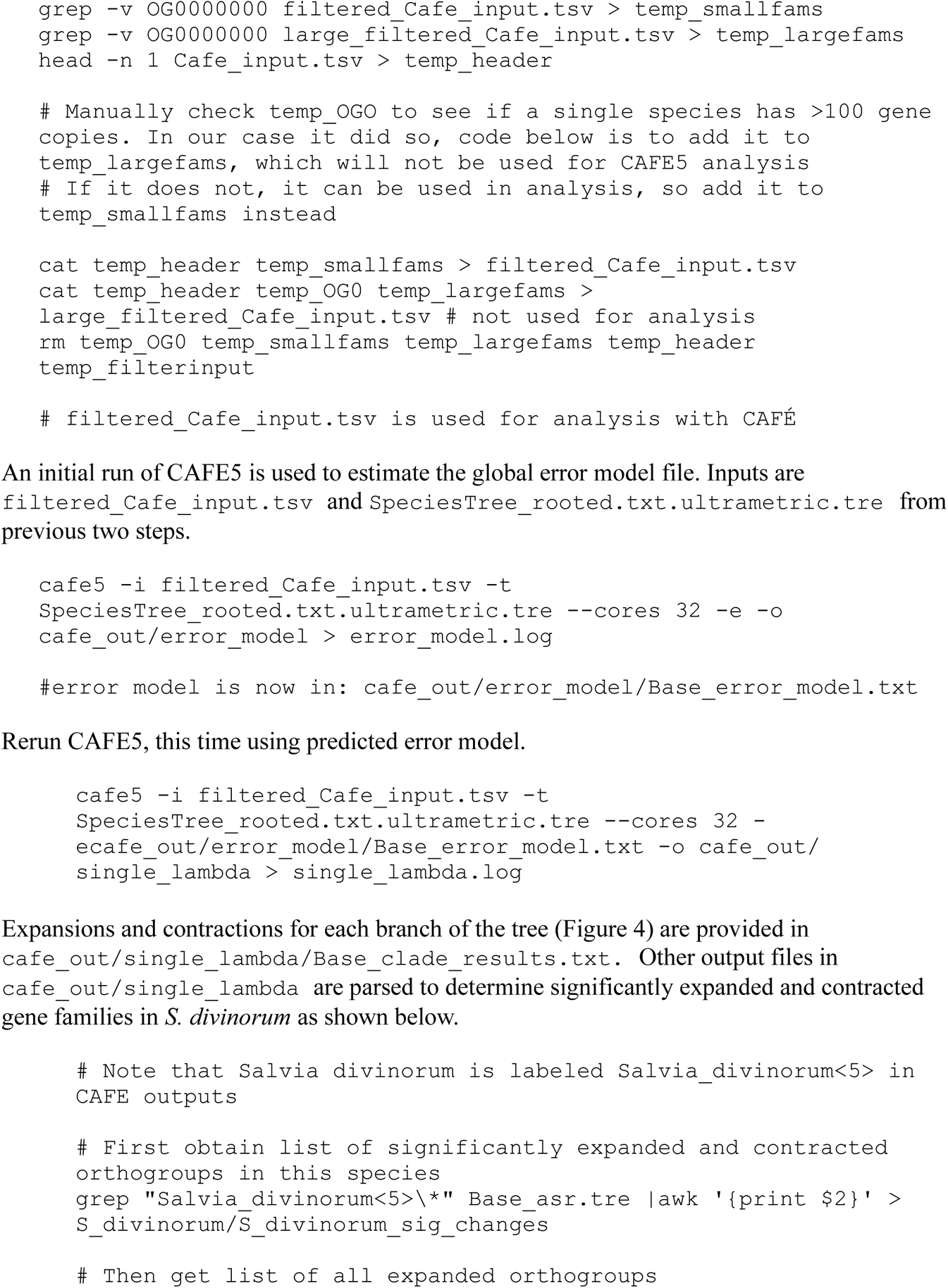

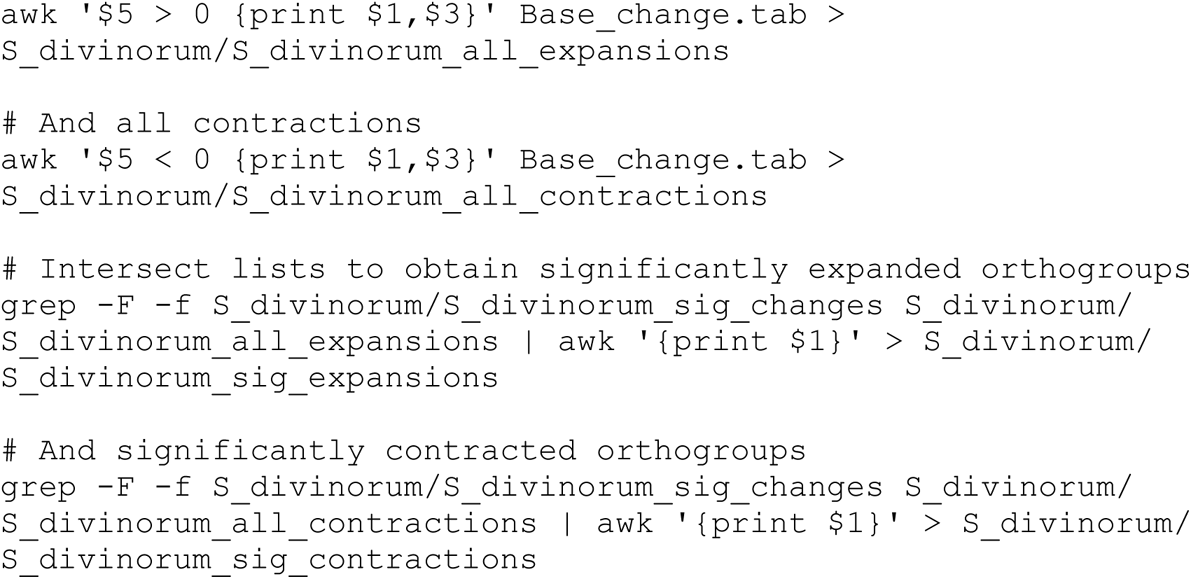

### 4. GO enrichment analysis of expanded gene families

We next extract fasta files of protein sequences in significantly expanded gene families of *S. divinorm*, from the OrthoFinder output folder `Orthogroup_Sequences`, using the bash command below. Once this is performed, orthogroup sequences are placed in a folder `expanded_Ogs`
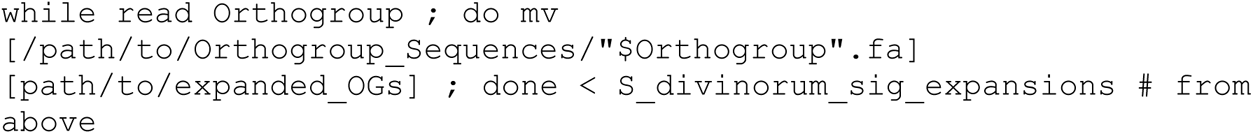

Each fasta file in `expanded_OGs` represents an expanded gene family of *S. divinorum*, but contains sequences from multiple species, so we need to extract the names of sequences corresponding specifically to *S. divinorum*. The headers of *S. divinorum* sequences have a unique locus tag `AAHA92`, which is used for extracting the names of genes in expanded orthogroups of *S. divinorum*.
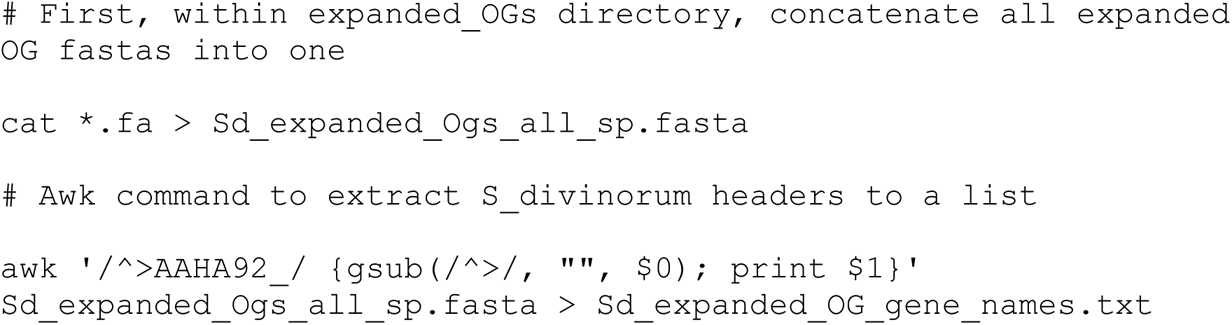

This provides a list of the names for proteins within significantly expanded gene families of *S. divinorum*. It is called `Sd_expanded_OG_gene_names.txt`, and is used as input for GO enrichment analysis using GOATOOLS (https://github.com/tanghaibao/goatools).

In addition to this list, GOATOOLS requires as input, a list of:

i. All gene names in the genome which have GO terms assigned, formatted as shown below. 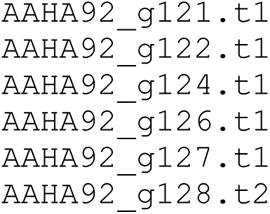
ii. GO terms for each gene in the genome formatted as shown below: 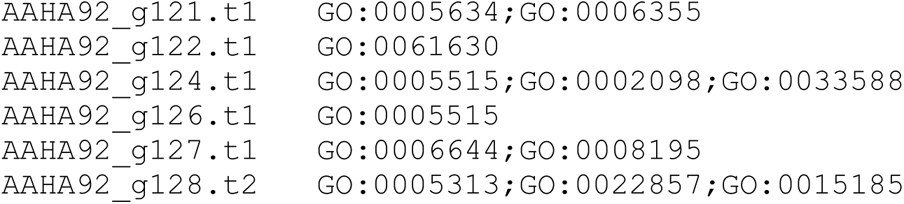
iii. All gene names in expanded gene families which have GO terms assigned, formatted as shown below.
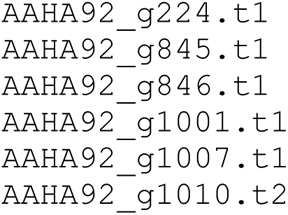

To generate these input files, we first extract names from the fasta file which contains longest isoform of each gene. We call this fasta file `longest_isoforms.faa` for simplicity.
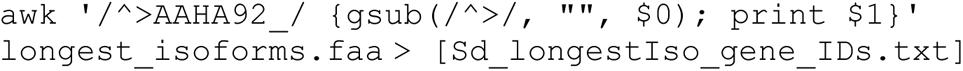

We then use the python code below to identify genes in Sd_longestIso_gene_IDs.txt that have assigned GO terms during functional annotation, and extract them to a line separated list. As input, we use the funannotate-generated gff file, which for simplicity we call `funannotate_out.gff.` As output we obtain the required list of genes which have GO terms assigned, and their corresponding GO terms formatted for input to GOATOOLS. We call this output file `all_genes_GO_terms.txt` as shown below.
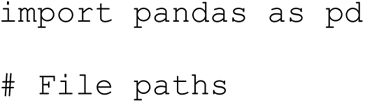

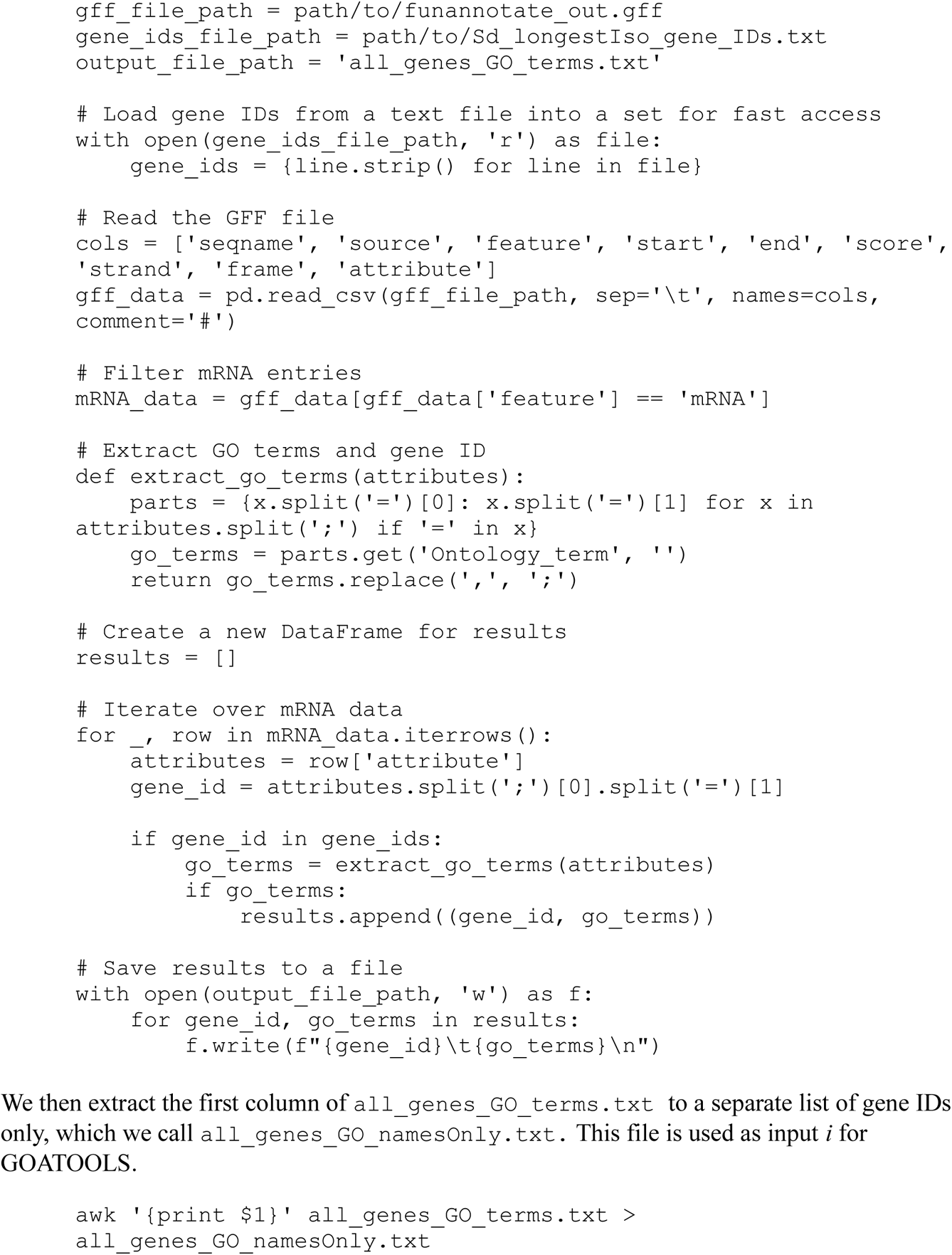

Finally, to generate input *iii* we filter `Sd_expanded_OG_gene_names.txt` such that it includes only genes annotated with GO terms. This can be done in python as below. We call the output `Sd_expanded_OG_gene_names_filtered.txt`
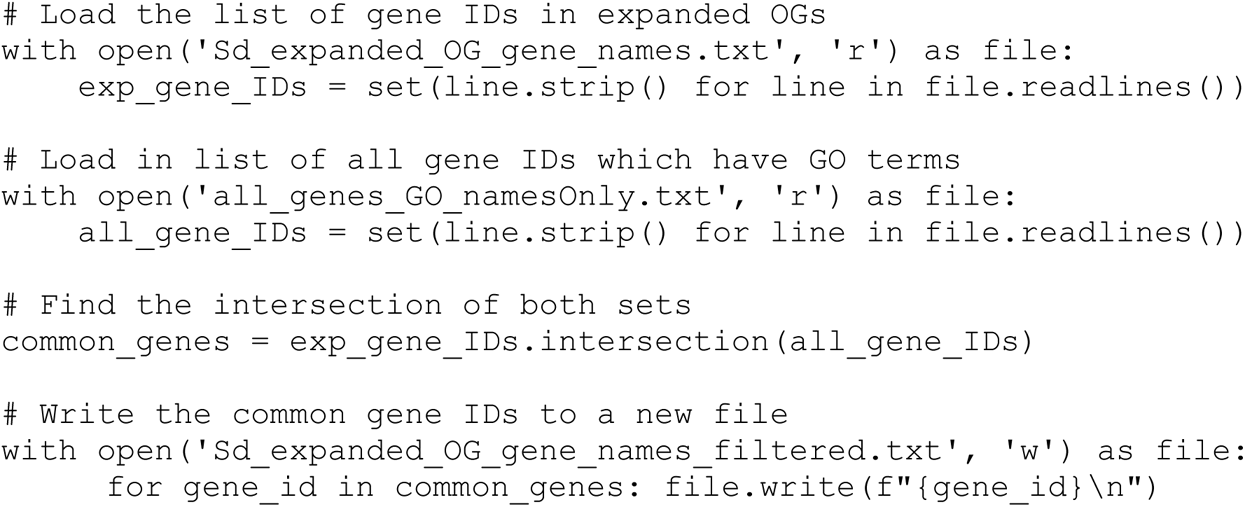

Now we have all required inputs for GOATOOLs. For instructions on installation, see (https://github.com/tanghaibao/goatools).

Software: GOATOOLS
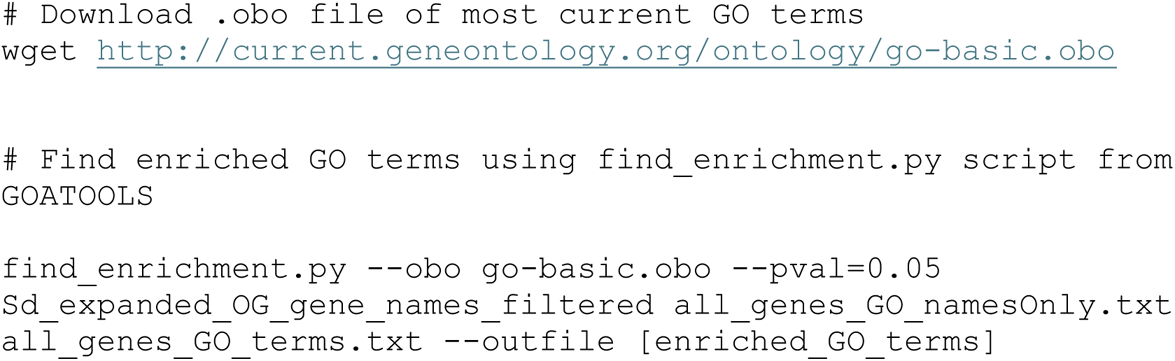

Output from GOATOOLs is used for plotting in Figure 4b and Supplemental Figure 5 after redundant GO terms are filtered using REVIGO (http://revigo.irb.hr/).

## G CIRCOS plotting

Figure 5 was generated using CIRCOS (https://circos.ca/). The command to generate the plot is below: 
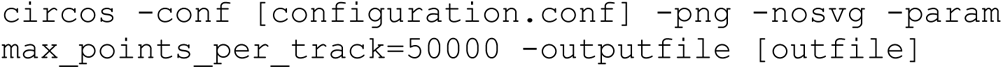

The configuration file `[configuration.conf]` is a text file with the following content

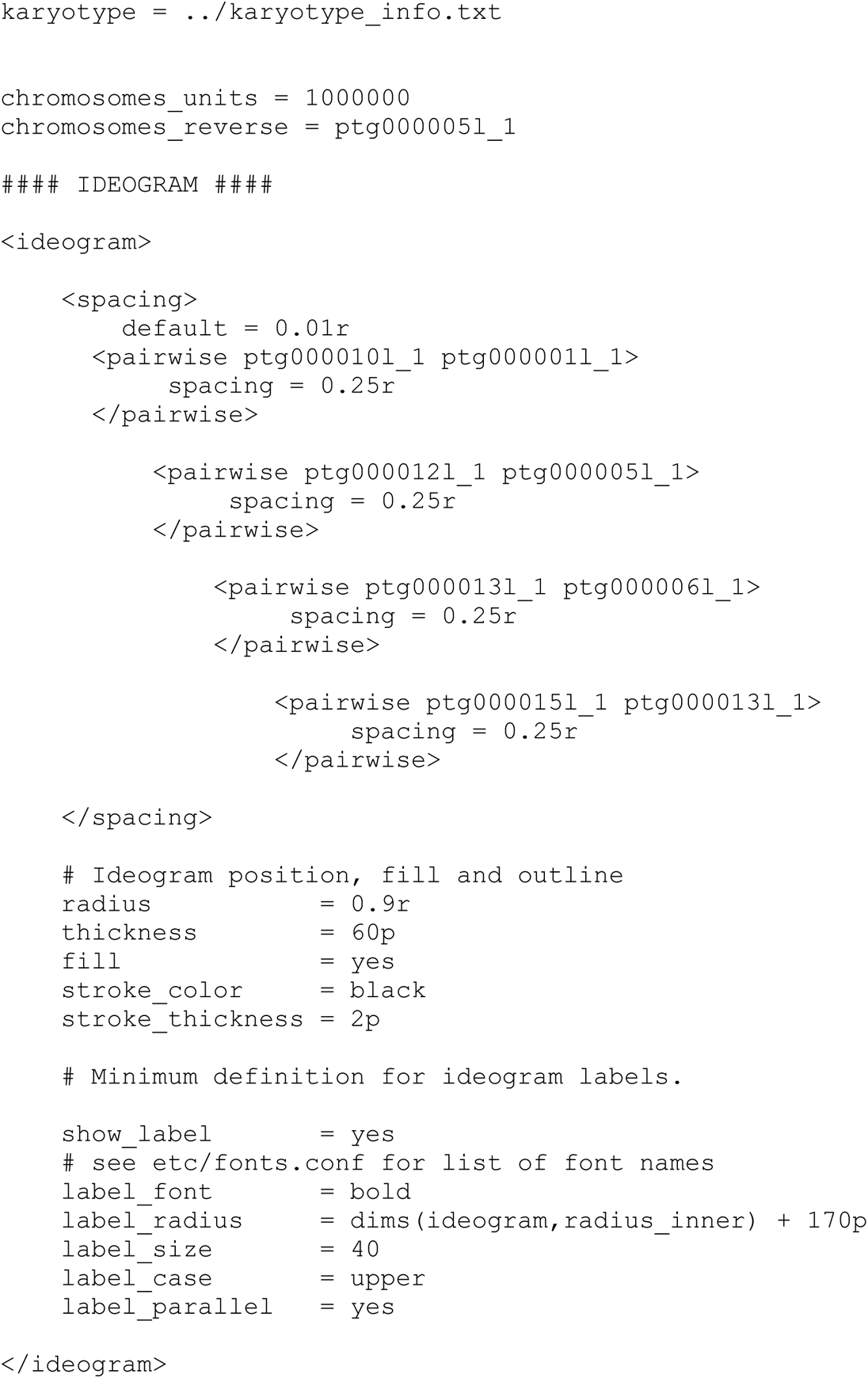

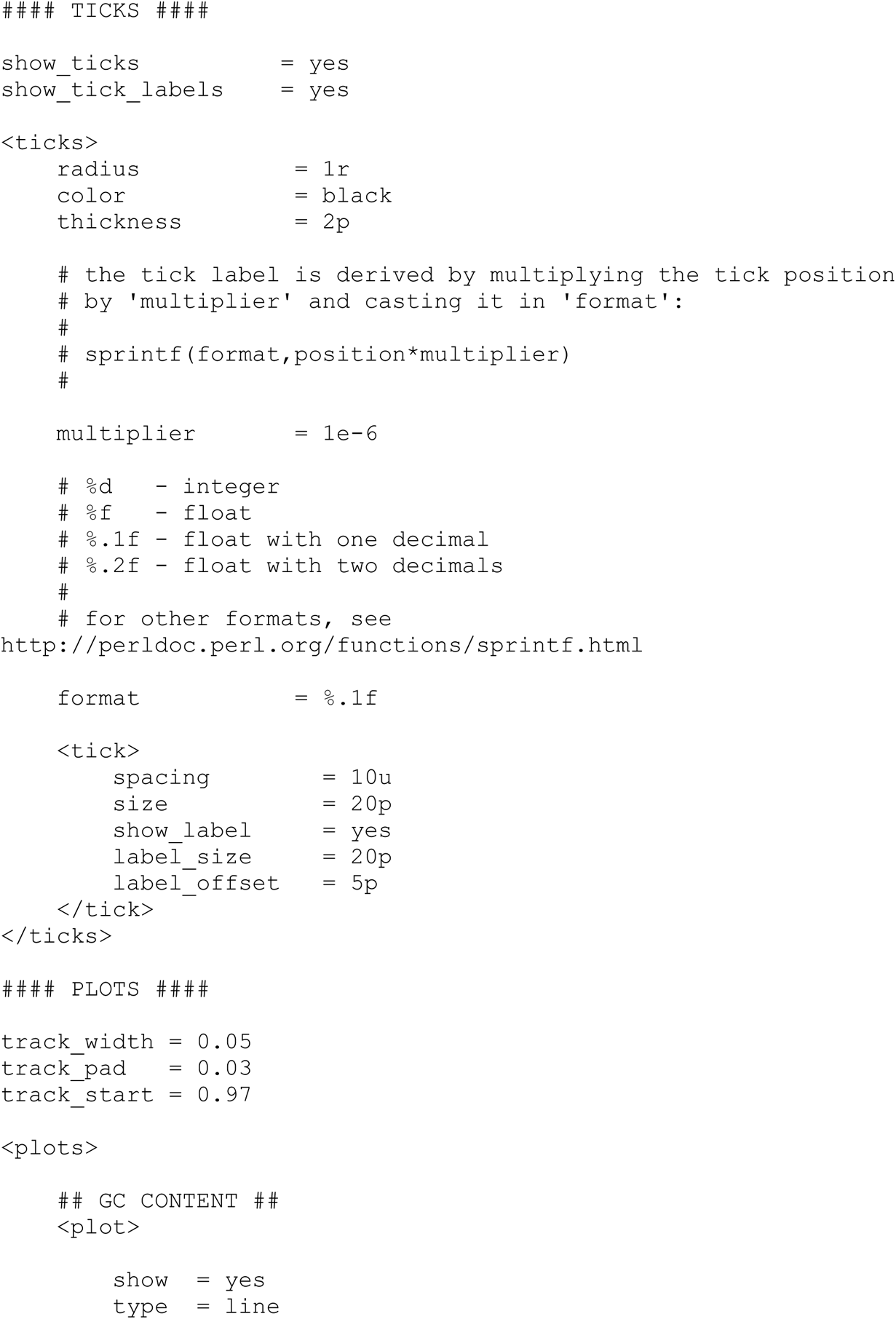

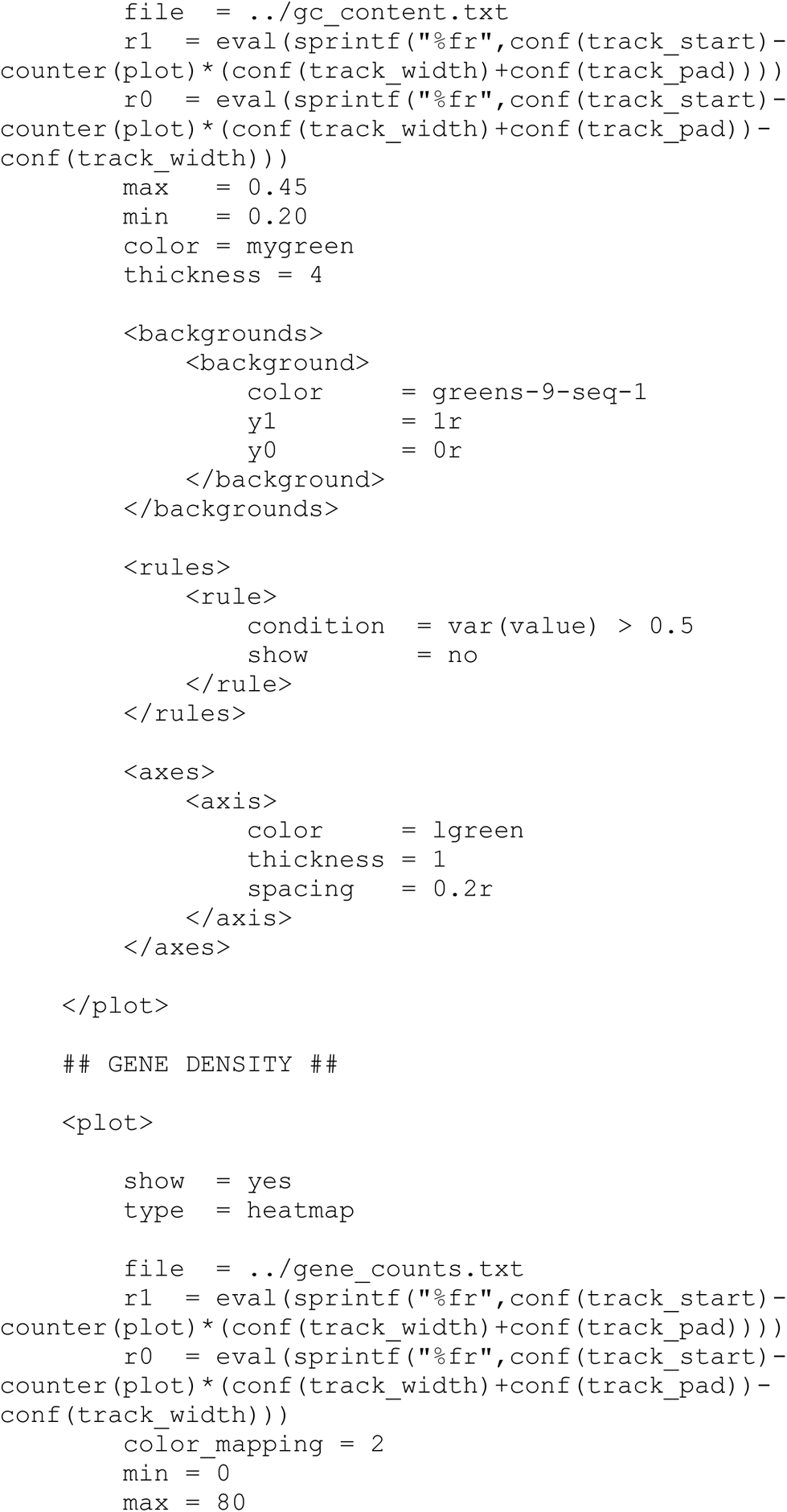

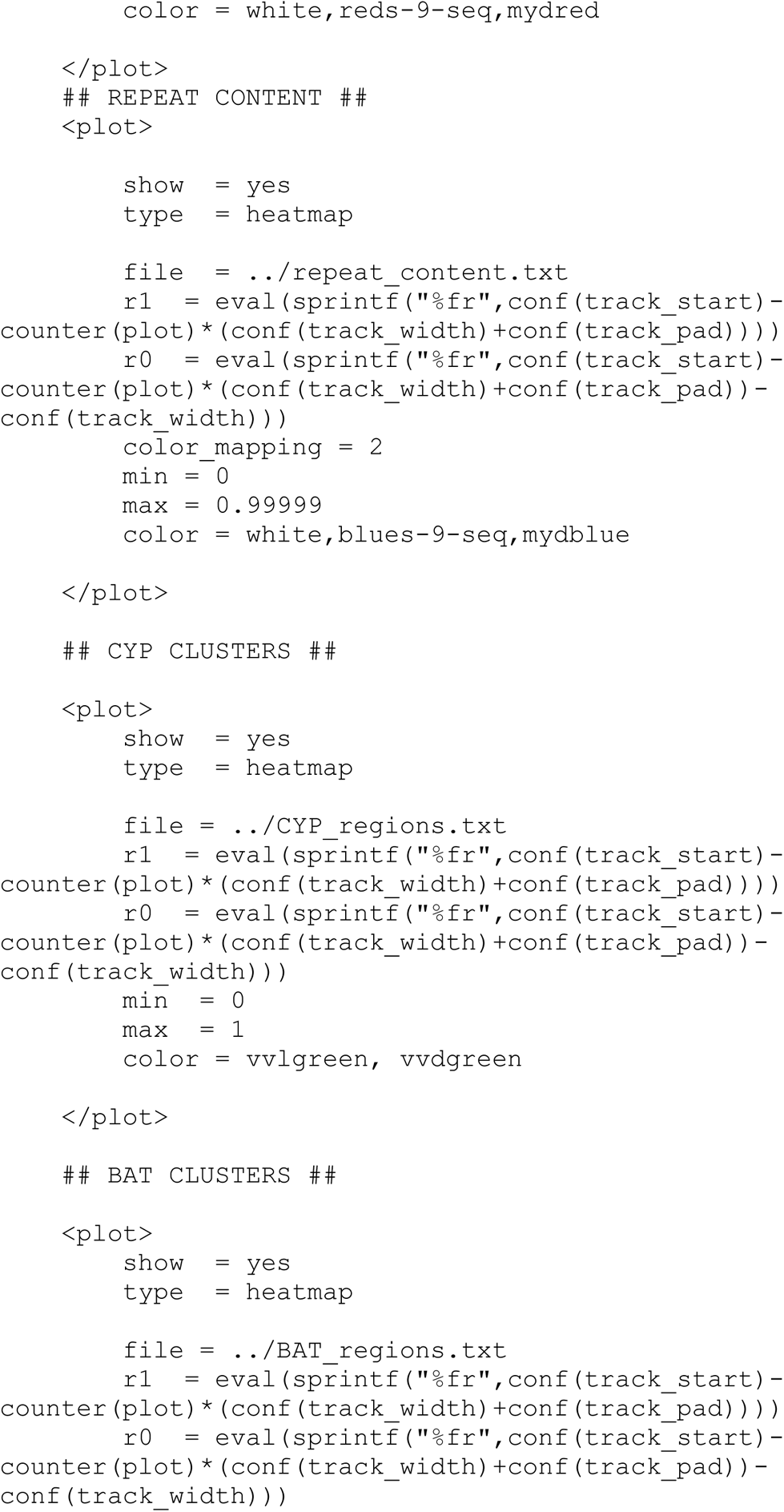

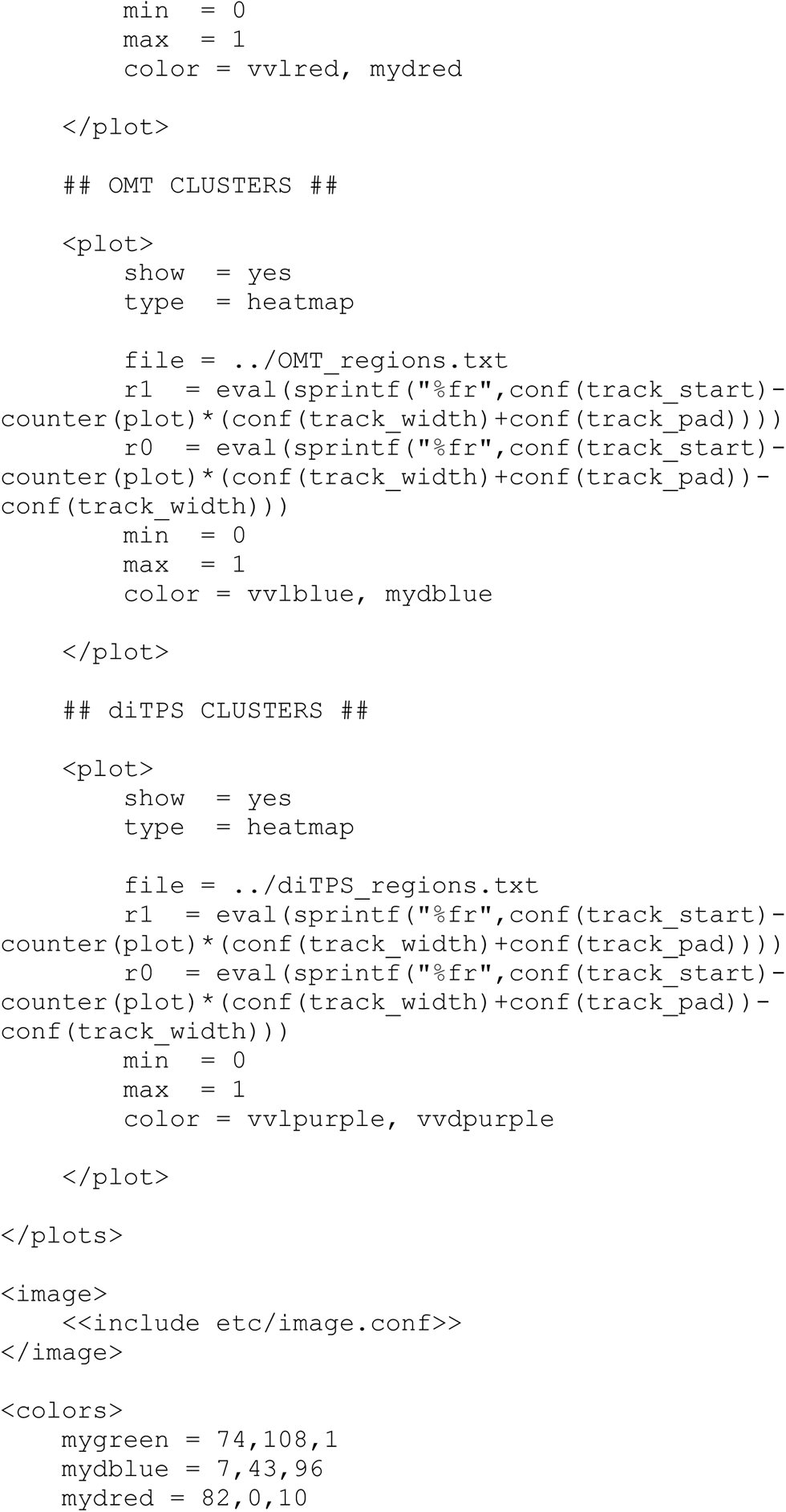

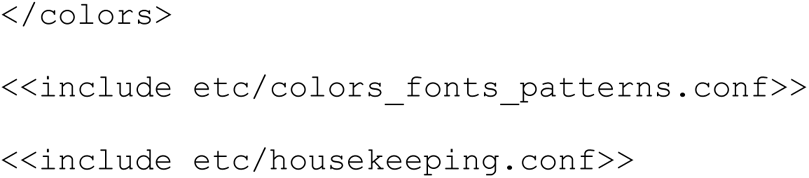

The karyotype file karyotype_info.txt is formatted as below:
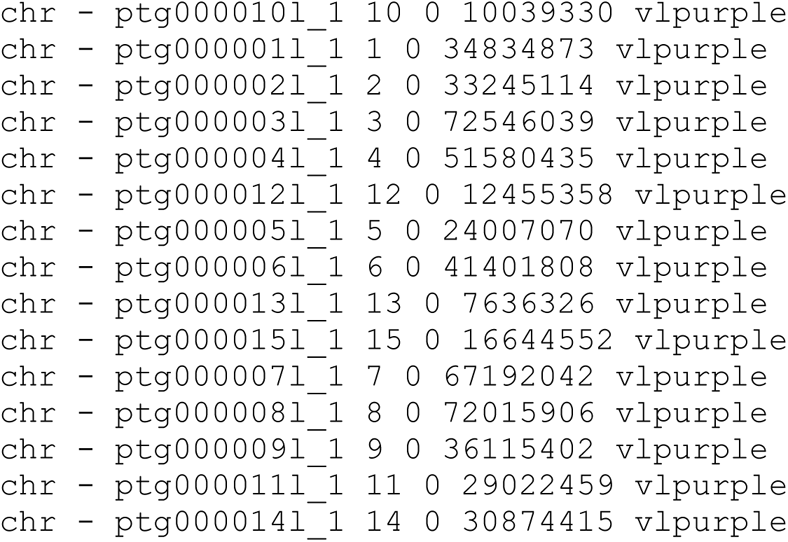

All files used for plotting in CIRCOS have 3 BED-formatted (ie., chromosome-start-end) and a statistic for plotting in the fourth column.

A subset of `gc_content.txt` is shown as below. This contains GC content averaged over 500 kb windows with a 100 kb step size.
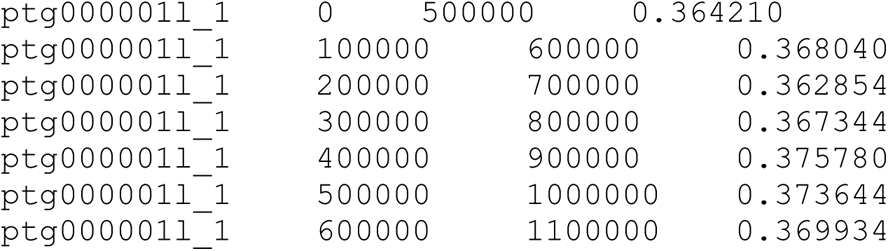

To generate `gc_content.txt` we used the following python functions. Use the assembly as `fasta_file` and assign output to `gc_content.txt`
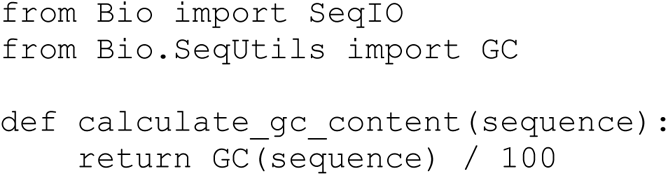

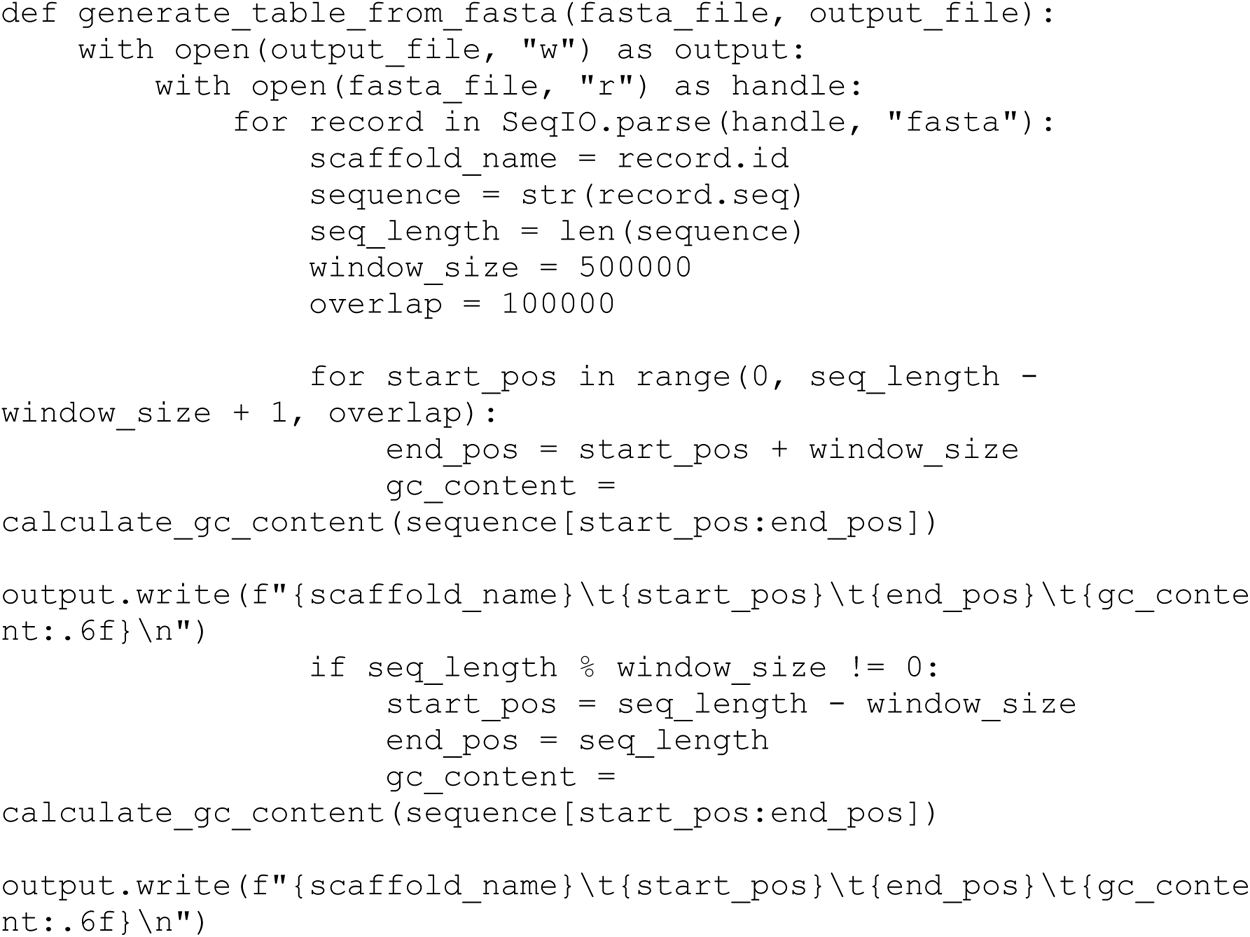

`repeat_content.txt` was similarly generated with the python functions below, using the softmasked genome (from RepeatMasker) as fasta_file and repeat_content.txt as output.
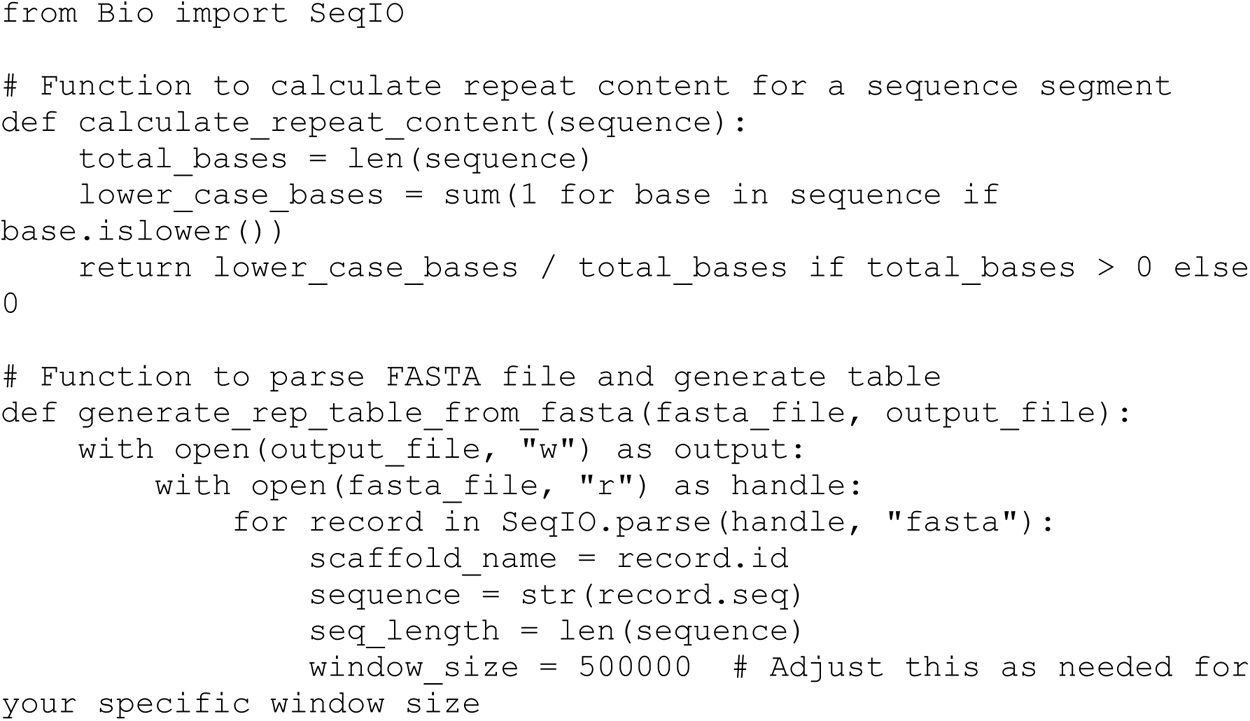

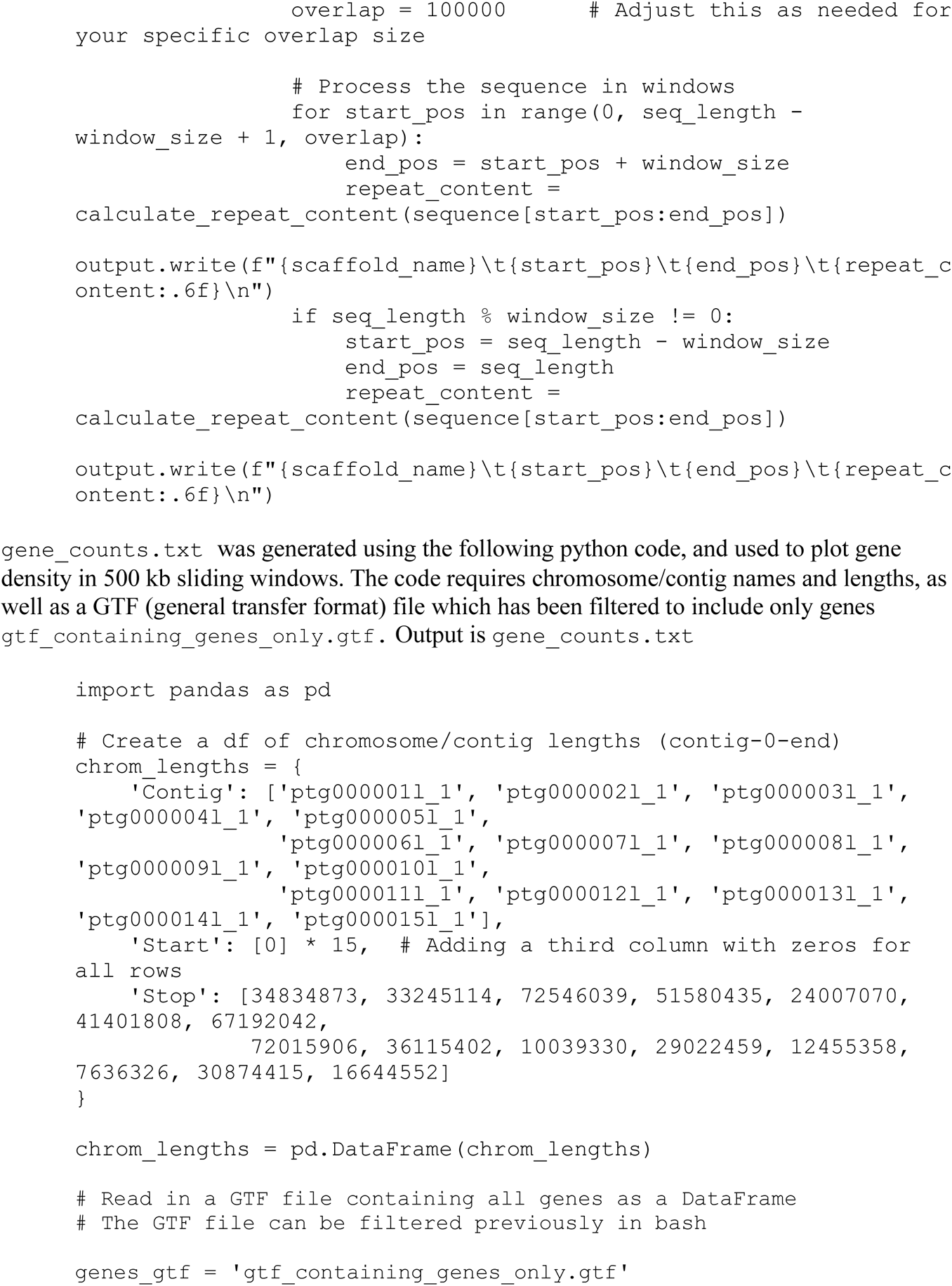

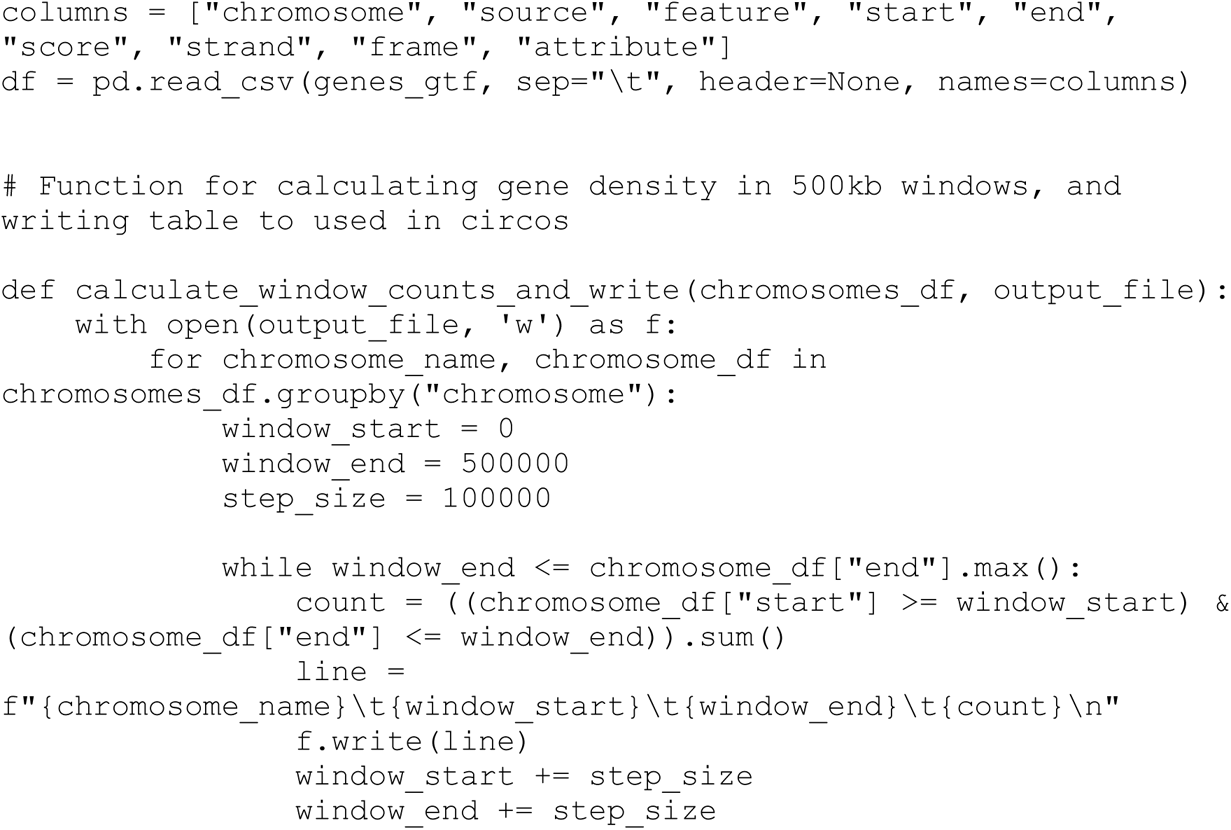

To plot clusters of genes encoding specific enzyme classes (inner rings), we generated files in where regions between the start and end of a gene were assigned 1 and all other regions of the genome were assigned 0. As an example, here are some BAHD acyl transferase (BAT) positions in chromosome 1 and 2:
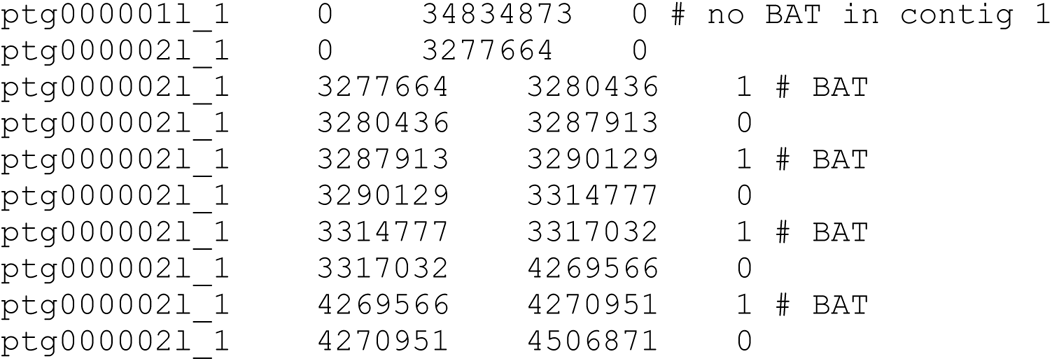

To generate gene region files (`CYP_regions.txt, BAT_regions.txt, OMT_regions.txt, diTPS_regions.txt`), we first obtained GTF files that were filtered to include only genes corresponding to each of the target classes. These were used in the following python code (using `BAT_genes.gtf to` generate `BAT_regions.txt` in this example):
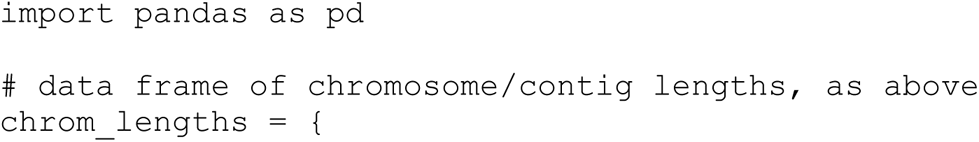

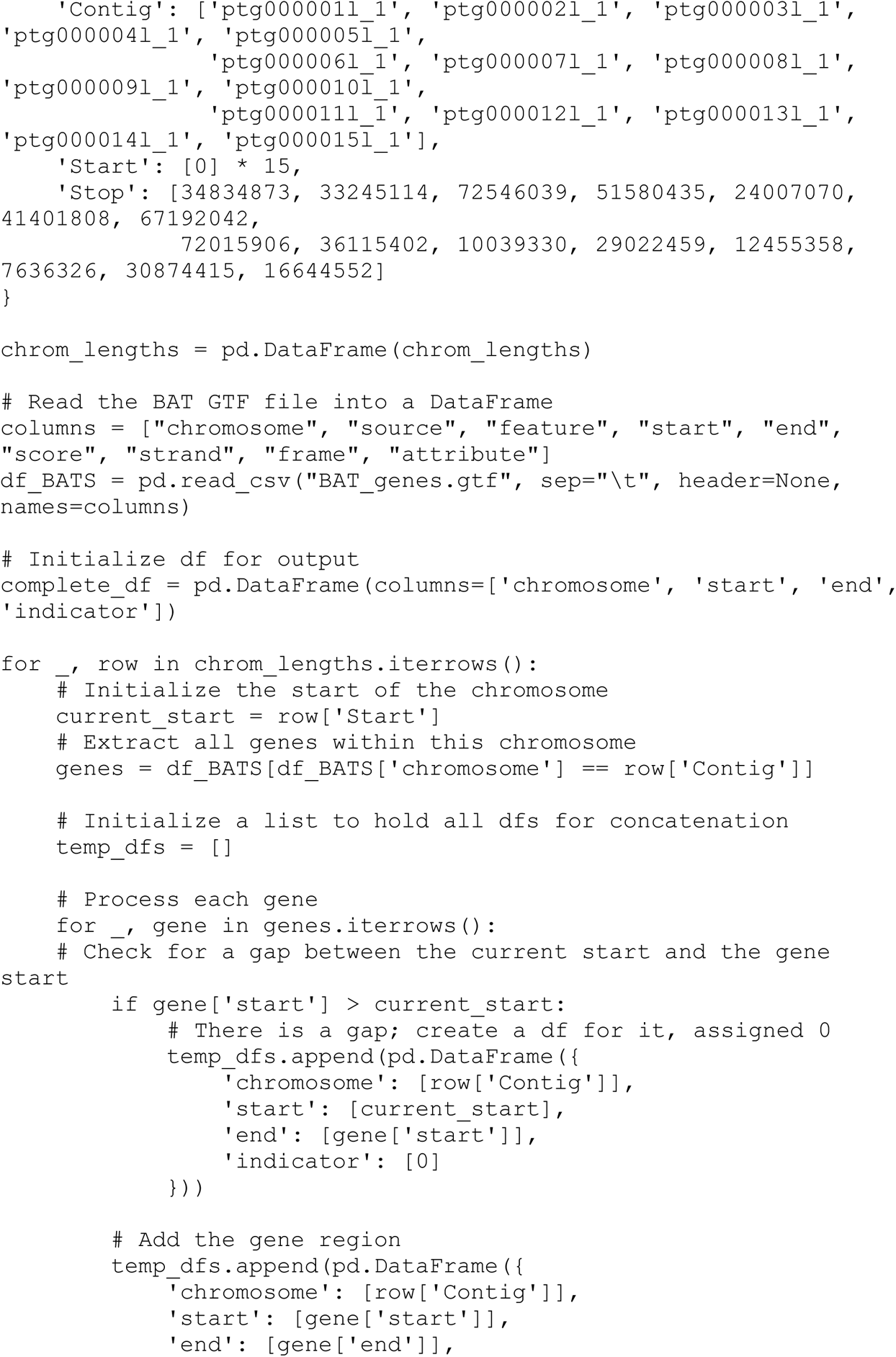

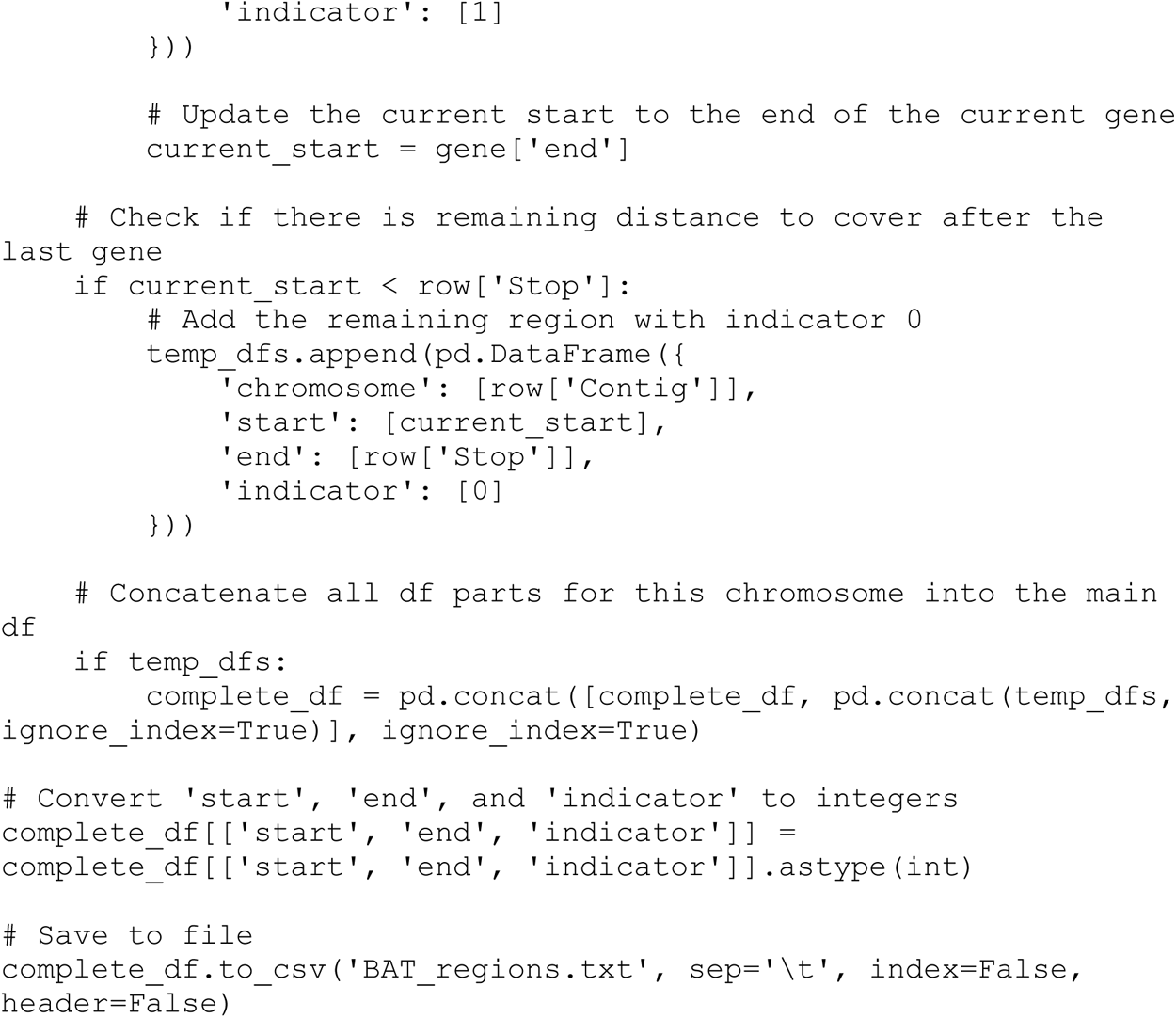

